# Engineering circular RNA expression systems to minimize contaminating linear RNA byproducts

**DOI:** 10.64898/2026.09.07.749908

**Authors:** Christopher J. Fields, Rina Fujiwara, Bradley W. Wright, Brett W. Stringer, Simon J. Conn, Jeremy E. Wilusz

## Abstract

Circular RNAs (circRNAs) are generated by backsplicing of eukaryotic protein-coding transcripts and can regulate microRNAs and RNA binding proteins, or serve as translation templates. Their covalently closed structure confers resistance to exonuclease-mediated degradation, extending their half-life and supporting their development as RNA therapeutics. However, existing overexpression methods often yield substantial contaminating linear RNAs, limiting their utility. Here, we systematically benchmarked plasmid-based circRNA overexpression strategies in human cells, comparing spliceosome- and ribozyme-based mechanisms across constructs incorporating widely used flanking sequences. The ribozyme-based Tornado system produced the highest circRNA yield but introduced extraneous “molecular scars” into the mature product. By contrast, spliceosome-mediated circularization using introns from the *Drosophila* Laccase2 gene, which contain imperfect complementary repeats, produced scarless circRNA with substantially lower linear RNA contamination. Linear RNA was further reduced by engineering the primary transcript to terminate in a non-polyadenylated end, increasing its susceptibility to exonucleases. Building on this optimized system, we developed a dual-output platform co-expressing a linear fluorescent reporter alongside a circRNA from a single promoter (CIRCUS, <u>circ</u>RNA and <u>u</u>pstream linear <u>s</u>ystem), enabling efficient screening of circRNA-driven cellular phenotypes, including site-specific A-to-I editing of target mRNAs. Together, this toolkit provides high-purity circRNA production suitable for mechanistic studies and circRNA-based therapeutic development.

## INTRODUCTION

Eukaryotic genes are typically interrupted by introns and can undergo alternative splicing to produce diverse RNA isoforms, including both linear mRNAs and circular RNAs (circRNAs). CircRNAs are covalently closed transcripts, and thousands have been identified across species [1–3]. They are generated through “backsplicing,” a noncanonical splicing reaction in which the spliceosome joins a downstream splice donor to an upstream splice acceptor—for example, linking the end of an exon to its beginning [4–6]. Although many circRNAs remain functionally uncharacterized, individual examples have been shown to regulate microRNA activity [3,7,8], interact with or scaffold RNA-binding proteins [9,10], modulate transcription [11,12], and, in some cases, serve as templates for protein translation [13–15]. As a class, circRNAs are more stable than linear mRNAs due to their intrinsic resistance to exonuclease-mediated degradation, allowing them to accumulate, in some cases, to levels more than tenfold higher than their corresponding linear mRNA isoforms [1,16,17]. This enhanced stability has driven growing interest in circRNAs as both endogenous regulators of physiological processes and as platforms for therapeutic development [18,19]. For example, engineered circRNAs have been designed to encode proteins for vaccines against infectious diseases [20,21], cancer immunotherapy [22], and protein replacement therapies; to act as microRNA sponges [23,24]; to function as aptamers that modulate inflammation [25,26]; and to serve as guide RNAs for site-specific DNA or RNA editing [27–29].

To facilitate their functional interrogation or therapeutic exploitation, circRNAs of interest can be produced either *in vitro* or in cells, with the latter leveraging intracellular machinery to transcribe and process circRNA-encoding vectors [19,30]. In cells, circRNA overexpression is commonly achieved using plasmids containing intronic sequences that promote backsplicing **(Figure 1A, left)**. Complementary intronic elements (e.g., Alu repeats) base-pair to bring splice sites into close proximity, thereby facilitating circularization. This process generates the mature circRNA along with a Y-shaped intermediate containing a 2’,5’ linkage, which is subsequently debranched and degraded [2,31–33]. Flanking exons are typically omitted from these constructs, limiting the production of alternatively spliced linear isoforms.

**Figure 1.**
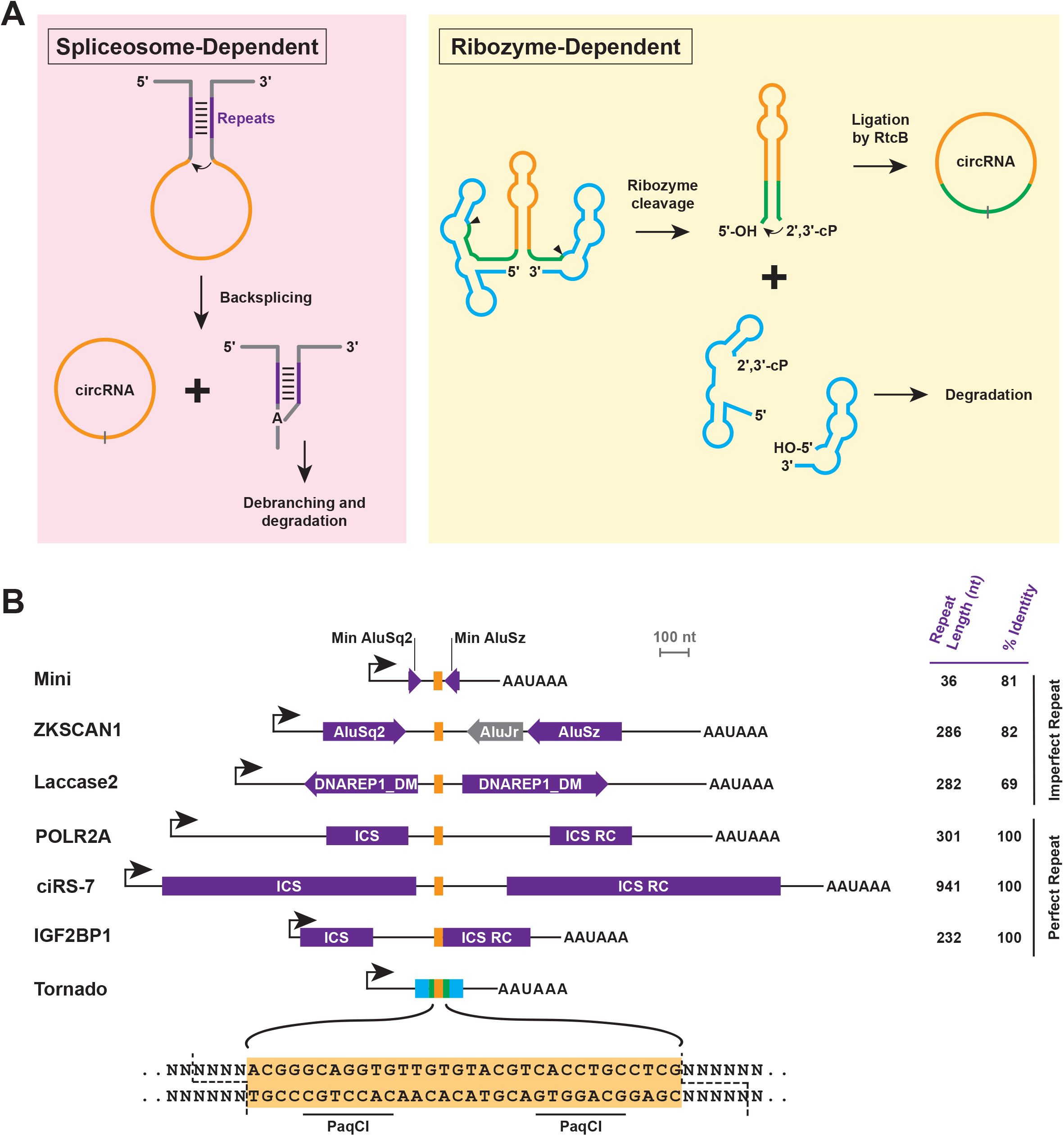
CircRNA overexpression strategies. **(A)** CircRNA overexpression in eukaryotic cells can be achieved using spliceosome- or ribozyme-dependent strategies. (Left) Complementary sequences (purple) within flanking introns promote backsplicing by the spliceosome, generating a Y-shaped byproduct that is subsequently debranched and degraded. (Right) In the Tornado system, a pair of self-cleaving Twister ribozymes (blue) enables ligation by the endogenous RtcB ligase to produce mature circRNA. Ribozyme cleavage sites are indicated by arrowheads, and the 47-nt “scar” (green) at the circRNA junction is shown. Upstream and downstream fragments generated by ribozyme cleavage are targeted for degradation; however, the 2’,3’-cyclic phosphate (2’,3’-cP) limits efficient removal of the upstream fragment. **(B)** A common multiple cloning site (MCS) containing a pair of PaqCI sites (orange) was inserted between intronic sequence pairs capable of driving circRNA production via spliceosome-dependent (Mini, ZKSCAN1, Laccase2, POLR2A, ciRS-7, IGF2BP1) or ribozyme-dependent (Tornado) mechanisms. Complementary sequences are denoted in purple, along with their lengths and percent sequence identity. ICS, intronic complementary sequence; RC, reverse complement.

Multiple intronic sequence pairs capable of driving backsplicing have been described, differing in intron length and degree of repeat complementarity **(Figure 1B)**. For example, introns from the human *ZKSCAN1* gene are sufficient to support backsplicing of the endogenous intervening exons as well as heterologous exonic sequences, and we previously defined minimal versions (denoted “Mini” in **Figure 1B**) that retain only splice sites and short (∼36 nt) inverted Alu repeats [32,34]. Introns containing longer, imperfect repeats—such as those derived from the *Drosophila Laccase2* gene [34]—as well as engineered, perfectly complementary sequences from human *POLR2A* [33], *ciRS-7* (*CDR1as*) [7], and *IGF2BP1* [35], have also been widely used to promote circRNA production **(Figure 1B)**. Despite their broad adoption, the full spectrum of transcripts generated from each of these vectors—particularly unintended linear byproducts—has not been systematically characterized. This gap has led to ongoing debate about whether phenotypes observed in cells can be attributed specifically to circRNAs or to other vector-derived transcripts [36–42].

Beyond spliceosome-mediated approaches, self-cleaving ribozymes provide an alternative strategy for circRNA generation in cells **(Figure 1A, right)** [43–45]. In the Tornado (<u>T</u>wister-<u>o</u>ptimized <u>RNA</u> for <u>d</u>urable <u>o</u>verexpression) system, Twister ribozymes flank the sequence of interest and self-cleave to generate termini (a 5’ hydroxyl and a 2’,3’-cyclic phosphate) that are compatible with ligation by the endogenous RtcB ligase [43]. Efficient ligation is facilitated by inclusion of a stem structure that resembles endogenous RtcB substrates. However, this approach introduces a 47-nt junction “scar” in the resulting circRNA **(Figure 1A, right** and **Supplementary Figure S1)**. While such scars are undesirable when studying endogenous circRNA function [46], they can be advantageous for distinguishing plasmid-derived transcripts from endogenous RNAs. Consistent with this rationale, in our prior work we introduced a multiple cloning site (MCS) between flanking intronic sequences to facilitate sequence insertion and enable discrimination of vector-derived transcripts from their endogenous counterparts **(Supplementary Figure S2A-B)** [32,34].

Here, we systematically evaluated widely used strategies for circRNA overexpression in human cells and developed improved plasmids that yield substantially reduced linear RNA byproducts. We benchmarked a panel of commonly used flanking intronic sequences, quantifying circRNA production alongside unintended linear transcripts, and observed marked variability across constructs, with some vectors producing substantially more linear byproduct than circRNA. Notably, abundant byproducts were easily missed depending on the detection approach used, underscoring the need for multiple orthogonal validation methods. We also observed that linear byproduct levels varied significantly by cell line. For high-performing designs—including those incorporating *Laccase2* introns—we further optimized vector architecture to increase the susceptibility of linear RNAs to exonucleolytic degradation while maintaining efficient circularization. To then enable easy screening of circRNA-driven cellular phenotypes, we generated a dual-output plasmid that cleanly co-expresses a linear fluorescent reporter alongside a circRNA from a single promoter. Collectively, this work provides a toolkit for producing high-purity circRNAs suitable for mechanistic studies and circRNA-based therapeutic development.

## MATERIALS AND METHODS

### Cell culture

HEK293FT cells were cultured in Dulbecco’s modified Eagle’s medium (DMEM) containing high glucose (Thermo Fisher Scientific 11995065) supplemented with 10% (v/v) fetal bovine serum (Thermo Fisher Scientific A5256701), 1% (v/v) non-essential amino acids (Thermo Fisher Scientific 11140050), 1% (v/v) GlutaMax (Thermo Fisher Scientific 35050061), and 1% (v/v) G418 Sulfate (Corning 30-234-CR). HeLa cells were cultured in DMEM containing high glucose (Thermo Fisher Scientific 11995065) supplemented with 10% (v/v) fetal bovine serum and 1% (v/v) penicillin-streptomycin (Thermo Fisher Scientific 15140122). HCT116 cells were cultured in DMEM without sodium pyruvate (Thermo Fisher Scientific 11965092) supplemented with 5% (v/v) fetal bovine serum and 1% (v/v) penicillin-streptomycin. SH-SY5Y cells were cultured in DMEM/Nutrient Mixture F-12 (Thermo Fisher Scientific 11320033) supplemented with 10% (v/v) fetal bovine serum and 1% (v/v) penicillin-streptomycin. All cells were cultured at 37°C and 5% CO_2_.

### Expression plasmids

To generate circRNA expression plasmids, the indicated sequences were inserted into pcDNA3.1(+) (Thermo Fisher Scientific V79020) between the CMV immediate early promoter and the bGH poly(A) signal. Intronic sequences derived from the *ZKSCAN1* [32,34], *Laccase2* [34], *POLR2A* [33], *ciRS-7* [7], and *IGF2BP1* [35] genes, as well as the Tornado system [43], have been previously described. Full details of the cloning procedures, including complete plasmid sequences, are provided in the **Supplementary Material**. To generate plasmids that transcribe unstable, non-polyadenylated linear RNAs **(Figure 5)**, the bGH poly(A) signal was replaced with wildtype or mutant versions of mouse mascRNA [47,48]. The guide RNA sequence for targeted A-to-I editing of the RAB7A 3’ UTR was designed using the GuideRNA-Forge tool [28]. All plasmids have been deposited at Addgene.

### Transfections and RNA isolation

HEK293FT, HeLa cells, and HCT116 (1 x 10^6^ per well) and SH-SY5Y cells (1.5 x 10^6^ per well) were seeded in 6-well plates in complete media and cultured overnight. The following day, cells were transfected with 1 μg of plasmid DNA using Lipofectamine 2000 (HEK293FT, HeLa, HCT116; Thermo Fisher Scientific 11668019) or 2.5 μg of plasmid DNA using Lipofectamine 3000 (SH-SY5Y; Thermo Fisher Scientific L3000008) according to the manufacturer’s instructions. Total RNA was extracted 24 h post-transfection using TRIzol reagent (Thermo Fisher Scientific 15596018) following the manufacturer’s instructions, except for experiments examining site-specific A-to-I editing **(Figure 7)** that were analyzed 48 h post-transfection.

### Northern blotting

Northern blots were performed using 1.2% denaturing formaldehyde agarose gels and NorthernMax reagents (Thermo Fisher Scientific), as previously described in detail [49]. RNA was transferred to a Biodyne A nylon membrane (Cytiva 60106) overnight by capillary transfer. Membranes were UV crosslinked (254 nm) and hybridized overnight with ^32^P-labeled oligonucleotide probes in ULTRAhyb-Oligo (Thermo Fisher Scientific AM8663) at 42°C, except for the mouse mascRNA-specific probe, which was hybridized overnight at 50°C. Blots were washed twice with 2x SSC, 0.5% SDS, then imaged using an Amersham Typhoon scanner (Cytiva). Signal quantification was performed using ImageQuant software (Cytiva).

Oligonucleotide probe sequences are provided in **Supplementary Table S1**. For detection of mascRNA and the Tornado upstream fragment, 8% polyacrylamide gels were used instead of denaturing formaldehyde agarose gels and transferred using a Trans-Blot SD Semi-Dry Transfer Cell (Bio-Rad 1703940) to Hybond N+ membrane (Cytiva RPN303B).

### RT-qPCR

5 μg of total RNA (quantified by NanoDrop) was treated with TURBO DNase (Thermo Fisher Scientific AM2238) in a 20 μL reaction following the manufacturer’s protocol. DNase was inactivated by the addition of EDTA (15 mM final concentration) and incubation at 75°C for 10 min. 1 μg of DNase-treated RNA was reverse transcribed in a 20 μL reaction using iScript Reverse Transcription Supermix for RT-qPCR (Bio-Rad 1708841) or HiScript IV RT SuperMix for qPCR (Vazyme R423-01) containing a mix of random primers and oligo(dT), according to the manufacturer’s instructions. cDNA was diluted up to 1:20 in nuclease-free H_2_O, and qPCR was performed using Power SYBR Green PCR Master Mix (Thermo Fisher Scientific 4368708).

Each 15 μL reaction contained 1.5 μL diluted cDNA, 7.5 μL 2x Power SYBR Green PCR Master Mix, and 6 μL of 1.5 μM gene-specific primer pairs. Primer sequences are provided in **Supplementary Table S2**. qPCR was performed on a QuantStudio 3 Real-Time PCR System (Thermo Fisher Scientific A28566) using clear plates (Thermo Fisher Scientific 4346907) with the following cycling conditions: 95°C for 10 min; 40 cycles of 95°C for 15 s and 60°C for 1 min; followed by a final cycle of 95°C for 10 s, 65°C for 1 min, and 97°C for 1 s. Melt curve analysis confirmed single amplicons. Threshold cycle (Ct) values were determined using the QuantStudio 3 system, and relative transcript levels (normalized to GAPDH) were calculated using the 2^-ΔΔCt^ method. RT-qPCR was performed using three independent biological replicates, each with two technical replicates.

To define the sequence present at the backsplice junction of eGFP circRNAs, cDNA was PCR amplified using the following forward (5’-ACCCTGAAGTTCATCTGCACC) and reverse (5’-GTCACGAACTCCAGCAGGAC) primers and then subjected to Sanger sequencing. To detect trans-spliced eGFP RNAs, cDNA was PCR amplified using the following forward (5’-CTACGTCCAGGAGCGCAC) and reverse (5’-GGGAGTGGCACCTTCCAG) primers and then subjected to Nanopore sequencing (Whole Amplicon Sequencing/Quintara Biosciences). Consensus sequences from Nanopore sequencing are provided in **Supplementary Table S3.**

### Western blotting

Cells were harvested 24 h after transfection using RIPA buffer (150 mM NaCl, 1% Triton X-100, 50 mM Tris-HCl, pH 7.5, 0.1% SDS, 0.5% sodium deoxycholate, 1% NP-40) supplemented with protease inhibitors (Roche 11836170001) and incubated on ice for 20 min. Lysates were clarified by centrifugation at 12,000 x *g* for 15 min at 4°C, and protein concentrations were determined using the Bio-Rad DC Assay (Bio-Rad 50000111). Equal amounts of protein (20 μg) were resolved on 4-12% Bis-Tris gels (Thermo Fisher Scientific NP0323) using MES SDS running buffer (Thermo Fisher Scientific NP0002) and transferred to PVDF membranes (Thermo Fisher Scientific 88520). Membranes were blocked in 5% (w/v) nonfat dry milk in TBST and incubated with primary antibodies diluted in blocking buffer overnight at 4°C. Membranes were washed with TBST (5 x 5 min) and incubated with HRP-conjugated secondary antibodies diluted in blocking buffer for 1 h at room temperature.

Following additional washes with TBST (5 x 5 min), proteins were detected using SuperSignal West Pico PLUS Chemiluminescent Substrate (Thermo Fisher Scientific PI34080) and imaged on an Amersham ImageQuant 800 system (Cytiva). Membranes were stripped using Restore PLUS Western Blot Stripping Buffer (Thermo Fisher Scientific 46430) and reprobed to detect loading controls. The intensity of protein bands was quantified using ImageQuant TL 10.2. The following antibodies were used: mouse anti-GFP (1:1000, Santa Cruz Biotechnology sc-9996), rabbit anti-mCherry (used to detect dTomato; 1:1000, Thermo Fisher Scientific PA5-34974), rabbit anti-GAPDH (1:5000, Proteintech 10494-1-AP), donkey anti-rabbit IgG/HRP (1:5000, Cytiva NA934), and sheep anti-mouse IgG/HRP (1:5000, Cytiva NA931).

### Ligation-based 3’ end cloning

To map the 3’ end of the Tornado upstream fragment, 10 μg of total RNA was incubated at 37°C for 1 h with T4 PNK (New England Biolabs, M0201) and then heat inactivated at 65°C for 20 min. A pre-adenylated oligo (miRNA Linker 3, IDT; 5’-rAppTTTAACCGCGAATTCCAG/3ddC/) was then ligated to the 3’ ends using the truncated form of T4 RNA Ligase 2 (New England Biolabs, M0242L). Ligation reactions were incubated at room temperature for 1 h followed by a phenol/chloroform extraction and ethanol precipitation.

Reverse transcription was then performed using iScript (Bio-Rad 1708841) as per the manufacturer’s instructions and 5 pmol of primer complementary to the 3’ linker sequence (5’-GACTAGCTGGAATTCGCGGTTAAA). cDNAs were amplified by PCR using AmpliTaq DNA Polymerase (Thermo Fisher Scientific N8080152) using the following forward (5’-GCTTACTGGCTTATCGAAAT) and reverse (5’-GACTAGCTGGAATTCGCGGTTAAA) primers and then subjected to Nanopore sequencing (Whole Amplicon Sequencing/Quintara Biosciences).

### FACS and Quantification of A-to-I Editing

HEK293FT cells (1 x 10^6^ per well) were seeded in 6-well plates and cultured overnight. Cells were transfected with 1 μg of CIRCUS plasmids containing either RAB7A or control guides using Lipofectamine 2000. At 48 h post-transfection, cells were washed with 1 mL PBS, incubated with 500 μL Trypsin for 5 min at 37°C, and then resuspended in 1 mL of DMEM supplemented with 2% FBS and 5 mM EDTA to prevent clumping. A subset of cells was saved (unsorted population) while the rest were sorted on the BD FACSAria II (100 µm nozzle) for dTomato expression (585 nm). Fluorescence was plotted on a log_10_ scale, and four cell populations were defined. The first population of cells exhibiting no dTomato expression was determined by a mock transfection control (“None”), and three non-overlapping sequential fluorescence populations were defined (Low, Medium, and High). Each cell population was collected. Total RNA was then extracted from each fraction, treated with DNase, and reverse transcribed as described above. cDNA was amplified using Q5 High-Fidelity DNA Polymerase (New England Biolabs M0491) using primers targeting RAB7A (Forward: 5’-AGTATGGCAGCAGGACAAGC and Reverse 5’-ACTCAGCCCACACCCTAGAATG) under the following cycling conditions: 98°C for 30 s; 35 cycles of 98°C for 30 s, 58°C for 30 s, and 72°C for 1 min; followed by a final cycle of 72°C for 2 min. Amplicons (∼1049 bp) were purified using the QIAquick PCR Purification Kit (Qiagen 28106) and then subjected to Nanopore sequencing (Whole Amplicon Sequencing/Quintara Biosciences). The frequency of A-to-G mutations, consistent with A-to-I RNA editing, was quantified at each adenosine in the PCR amplicon. Per base data from Nanopore sequencing are provided in **Supplementary Table S4**.

### Statistical analysis

For Northern blots, Western blots, RT-qPCRs and A-to-I editing analyses, statistical significance for comparison of means was assessed by unpaired t-test using GraphPad Prism. Statistical analyses and error bars are explained in the corresponding figure legends, when applicable.

## RESULTS

### Quantifying circRNA and linear byproducts from common circRNA overexpression strategies

To systematically compare the outputs of circRNA overexpression plasmids, we generated a panel of constructs incorporating commonly used intronic sequence pairs that promote backsplicing (*ZKSCAN1* [mini and long versions], *Laccase2*, *POLR2A*, *ciRS-7*, and *IGF2BP1*) [7,32–35] **(Figure 1B)**. All of these constructs were standardized using a shared PaqCI-based multiple cloning site (MCS) **(Figure 1B, orange)**, enabling seamless insertion of exonic sequences via restriction cloning or Gibson assembly without introducing extraneous nucleotides (“scars”) into the resulting circRNA [46] **(Supplementary Figure S2C)**. Although this design allows flexible cloning of diverse sequences, functional splice sites remain necessary to support efficient backsplicing. We also generated a construct containing mini *ZKSCAN1* introns flanking a previously described MCS **(Supplementary Figure S2A)** that introduces a junction scar into the mature circRNA [32] **(Supplementary Figure S2B)**, as well as a construct built on the ribozyme-based Tornado system [43] **(Figure 1B)**.

To assess circRNA production, we inserted into each construct an exon encoding a split eGFP open reading frame (ORF) together with the encephalomyocarditis virus (EMCV) internal ribosome entry site (IRES) **(Figure 2A)**. In linear (unspliced) transcripts, the ORF is disrupted because the start codon lies downstream of the stop codon, thereby preventing eGFP protein expression. In contrast, backsplicing restores a continuous ORF, enabling translation of full-length eGFP when the junction is scarless **(Figure 2A, right)**. However, junction “scars” that result from the previously described Mini MCS design **(Supplementary Figure S2B)** and the Tornado system **(Supplementary Figure S1B)** are expected to disrupt the ORF and prevent eGFP protein production.

**Figure 2.**
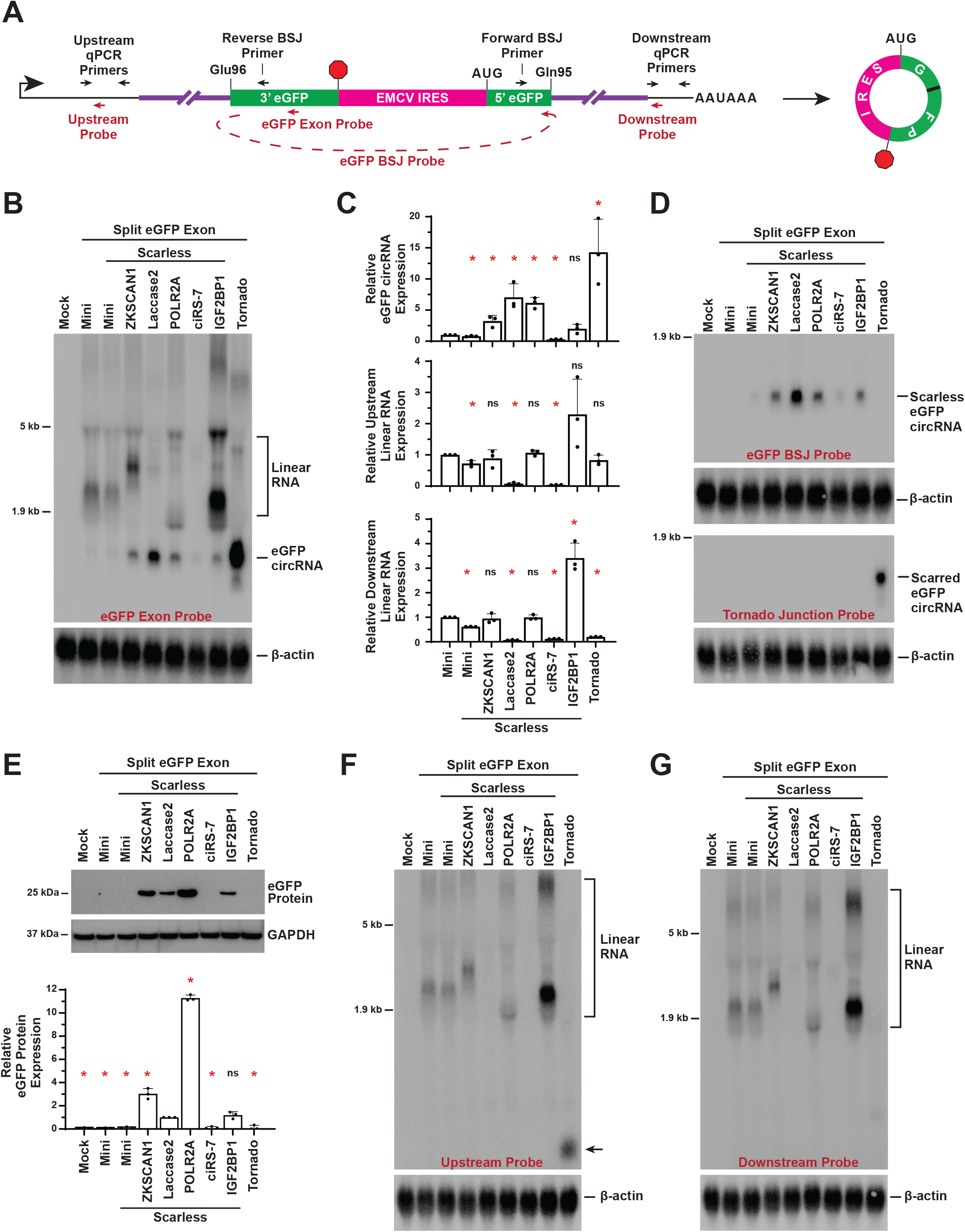
Comparison of circRNA and linear outputs of commonly used circRNA overexpression strategies in HEK293FT cells. **(A)** An exon encoding a split eGFP ORF together with the EMCV IRES was inserted between various intronic sequence pairs (purple). qPCR primers (black) targeting common regions upstream and downstream of the intronic sequences were used to quantify linear RNA levels, whereas primers spanning the backsplice junction (BSJ) were used to quantify circRNA. Oligonucleotide probes (red) for Northern blotting were similarly designed to detect transcripts containing the indicated complementary regions. **(B-G)** Plasmids expressing the split eGFP/EMCV IRES exon between the indicated intronic sequences were transfected into HEK293FT cells, and total RNA and protein were isolated after 24 h. **(B, D, F, G)** 18 μg of total RNA was analyzed by Northern blot using probes complementary to the **(B)** eGFP ORF, **(D)** backsplice or Tornado junction, **(F)** upstream common region, or **(G)** downstream common region. β-actin was used as a loading control. **(C)** RT-qPCR was used to quantify circRNA and linear RNA expression from each plasmid. Data are normalized to the Mini plasmid that produces a scarred BSJ **(Figure S2A)** and are presented as mean ± SD (N = 3). p values were generated with unpaired t-tests, comparing each construct to the Mini plasmid that produces a scarred BSJ. *p < 0.05; ns, not significant. Note that the junction scar increases the length of the PCR amplicon for circRNAs derived from the Mini and Tornado plasmids. **(E)** 20 μg of total protein was analyzed by immunoblotting using an anti-eGFP antibody. GAPDH was used as a loading control. Data are normalized to the *Laccase2* intron-containing construct and are presented as mean ± SD (N = 3). p values were generated with unpaired t-tests, comparing each construct to the *Laccase2* intron-containing construct. *p < 0.05; ns, not significant.

Each construct was transfected into HEK293FT cells, and RNA and protein outputs were analyzed 24 h post-transfection. Northern blotting using a probe complementary to the eGFP exon confirmed that all constructs produced mature eGFP circRNA **(Figure 2B** and **Supplementary Figure S3)**. The Tornado system yielded the highest circRNA levels **(Figure 2B-C)**; however, the presence of a junction scar—confirmed by Northern blotting **(Figure 2D)** and RT-PCR across the junction **(Supplementary Figure S1B)**—prevented expression of full-length eGFP protein **(Figure 2E)**. Among spliceosome-based constructs, introns derived from *ZKSCAN1*, *Laccase2*, and *POLR2A* supported the highest levels of circRNA production **(Figure 2B-C)**, and in each case, the absence of a junction scar **(Figure 2D** and **Supplementary Figure S2C)** enabled eGFP protein expression **(Figure 2E)**.

Notably, despite encoding identical circRNA sequences, these spliceosome-based constructs exhibited clear discrepancies between circRNA abundance and protein output (compare **Figure 2B-C** and **2E**), indicating that protein expression is not a reliable proxy for circRNA levels. Removal of one of the intronic repeat sequences required for backsplicing from each construct abolished circRNA accumulation **(Supplementary Figure S4A-B)** and markedly reduced eGFP protein expression **(Supplementary Figure S4C)**. However, residual protein production persisted for the *ZKSCAN1* intron construct. We reasoned this residual activity may reflect trans-splicing events that generate translatable exon concatemers [36,37] **(Supplementary Figure S5A)**, and RT-PCR revealed the presence of higher-molecular-weight RNA species consistent with linear, trans-spliced transcripts from many of our constructs **(Supplementary Figure S5B-C** and **Supplementary Table S3)**.

We next examined how repeat architecture influences backsplicing efficiency. Although perfectly complementary repeats are widely used to drive circRNA overexpression [7,33,35], *ciRS-7* (*CDR1as*) introns engineered to be perfectly complementary over 941 nt produced relatively low levels of eGFP circRNA **(Figure 2B-C)**. Substituting stronger splice sites [50] from *ZKSCAN1* or *Laccase2* **(Supplementary Figure S6A)** into the *ciRS-7* construct did not enhance circRNA production **(Supplementary Figure S6B)**, indicating that splice site strength is not the primary limiting factor in this context. Instead, these findings suggest that extensive perfect complementarity may limit backsplicing efficiency. In contrast, introns containing imperfect repeats, such as those derived from *ZKSCAN1* and *Laccase2* **(Figure 1B)**, were more effective at promoting circRNA production **(Figure 2B-C)**. Notably, constructs containing perfectly versus imperfectly complementary repeats or the Tornado sequences elicited no significant changes in well-established immune response genes, including RIG-I, MDA5, and OAS **(Supplementary Figure S7)**, indicating that these effects are unlikely to be driven by differential immune activation.

In addition to circRNA production, we assessed the extent of linear RNA byproducts by analyzing sequences common to all constructs **(Figure 2A)**. Northern blotting with a probe complementary to the eGFP ORF revealed abundant higher-molecular-weight RNA species for several constructs, in some cases (e.g., the Mini and *IGF2BP1* introns) exceeding circRNA levels **(Figure 2B)**. Many of these species were also detected using probes targeting the 5’ **(Figure 2F)** and 3’ regions **(Figure 2G)** of the pre-mRNA, consistent with unspliced linear transcripts. In contrast, fewer such species were observed for the Tornado system **(Figure 2B)**; however, a prominent smaller RNA species was detected with the upstream Northern blot probe **(Figure 2F)**. We reasoned that this species corresponds to the upstream fragment generated by ribozyme cleavage, whose 3’ end is partially protected from degradation by a 2’,3’-cyclic phosphate **(Figure 1A)**. Consistent with this model, a ∼183-nt RNA was detected using a Northern probe complementary to upstream but not downstream of the ribozyme cleavage site **(Supplementary Figure S8A-B)**. Treatment with T4 polynucleotide kinase (T4 PNK), which converts 2’,3’-cyclic phosphates to 3’-hydroxyl termini [51], increased ligation efficiency to this fragment **(Supplementary Figure S8C)**, and sequencing confirmed that it terminates at the cleavage site **(Supplementary Figure S8D)**.

Together, these analyses demonstrate that linear RNA byproducts from circRNA overexpression constructs can be substantial and are highly dependent on construct design. Their detection is further influenced by assay design, particularly the specific probes or primers used. Considering both eGFP circRNA yield and linear RNA contamination, we suggest that constructs containing *Laccase2* introns provided the most favorable balance of efficient backsplicing and minimal byproduct accumulation in HEK293FT cells **(Figure 2C)**.

### Internal circRNA introns are efficiently removed with spliceosome-based, but not ribozyme-based constructs

We next tested the ability of each pair of flanking sequences to drive overexpression of an endogenous circRNA **(Figure 3)**. The long noncoding RNA LINC00632 gives rise to ciRS-7 (also known as CDR1as), one of the best-characterized circRNAs, which contains ∼70 miR-7 binding sites and regulates neuronal activity [3,7,8,52,53]. The full-length ciRS-7 transcript is 1,485 nt; however, a subset of transcripts is shorter due to alternative splicing that removes an internal 184-nt intron [7,54] **(Figure 3A)**. The 1,485 nt ciRS-7 sequence was inserted into each circRNA overexpression construct and transfected into HEK293FT cells **(Figure 3B-C)**. All tested constructs resulted in >5-fold overexpression of ciRS-7 relative to endogenous levels **(Supplementary Figure S9A).** Consistent with the eGFP reporter results, the Tornado system produced the highest levels of ciRS-7, particularly the long isoform **(Figure 3B-C)**, but all transcripts contained a junction scar **(Supplementary Figure S9B-C)**. In addition, the Tornado upstream cleavage fragment accumulated in cells **(Supplementary Figure S9C)**, analogous to what we observed with the eGFP circRNA plasmid **(Figure 2F)**. In contrast, constructs containing *ZKSCAN1*, *Laccase2*, and *POLR2A* introns generated substantial levels of both long and short ciRS-7 isoforms (denoted ciRS-7 and ciRS-7 Δintron, respectively), all lacking junction scars **(Figure 3B-C** and **Supplementary Figure S9C)**. These spliceosome-based constructs also showed reduced accumulation of upstream linear RNA byproducts. However, substantial levels of downstream linear RNAs were detected for constructs containing *POLR2A* introns **(Figure 3C** and **Supplementary Figure S9C)**.

**Figure 3.**
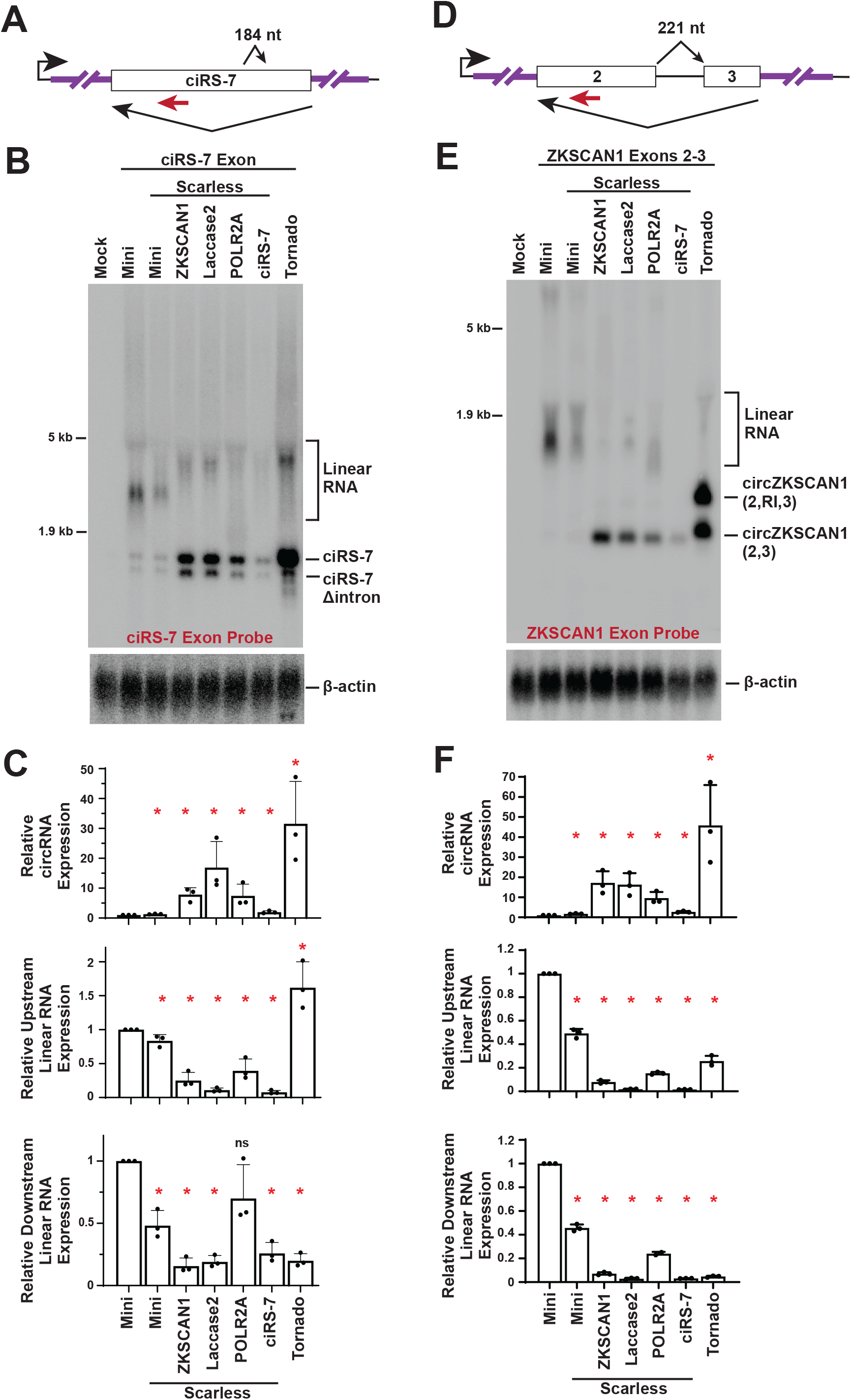
Overexpression of endogenous circRNAs.(A) Backsplicing of the ciRS-7 (also known as CDR1as) exon yields a 1,485-nt circRNA or a shorter isoform when an internal 184-nt intron is removed. The oligonucleotide probe used for Northern blotting is indicated in red. **(B-C)** Plasmids expressing the ciRS-7 exon flanked by the indicated intronic sequences were transfected into HEK293FT cells, and total RNA was isolated after 24 h. **(B)** 18 μg of total RNA was analyzed by Northern blot using a probe complementary to the ciRS-7 exon. β-actin was used as a loading control. **(C)** RT-qPCR was used to quantify circRNA (full-length ciRS-7 and ciRS-7 Δintron) and linear RNA expression from each plasmid. Data are normalized to the Mini plasmid that produces a scarred BSJ and are presented as mean ± SD (N = 3). p values were generated with unpaired t-tests, comparing each construct to the Mini plasmid that produces a scarred BSJ. *p < 0.05; ns, not significant. Note that the junction scar increases the length of the PCR amplicon for circRNAs derived from the Mini and Tornado plasmids. **(D)** Backsplicing of exons 2 and 3 of human ZKSCAN1 generates circZKSCAN1(2,3), a 668-nt circRNA. The 221-nt internal intron is typically constitutively removed; its retention would instead produce an 889-nt isoform, circZKSCAN1(2,RI,3). The oligonucleotide probe used for Northern blotting is indicated in red. RI, retained intron. **(E-F)** Plasmids expressing ZKSCAN1 exons flanked by the indicated intronic sequences were transfected into HEK293FT cells, and total RNA was isolated after 24 h. **(E)** 18 μg of total RNA was analyzed by Northern blot using a probe complementary to ZKSCAN1 exon 2. β-actin was used as a loading control. **(F)** RT-qPCR was used to quantify circRNA (circZKSCAN1(2,3) and circZKSCAN1(2,RI,3)) and linear RNA expression from each plasmid. Data are normalized to the Mini plasmid that produces a scarred BSJ and presented as mean ± SD (N = 3). p values were generated with unpaired t-tests, comparing each construct to the Mini plasmid that produces a scarred BSJ. *p < 0.05. As above, the junction scar increases the length of circRNA PCR amplicons derived from the Mini and Tornado constructs.

To further evaluate the ability of each system to generate a multi-exon circRNA requiring constitutive intron removal, we examined exons 2 and 3 of ZKSCAN1, which are spliced together to produce a 668-nt circRNA (circZKSCAN1(2,3)) [1–3,32] **(Figure 3D)**. The intervening 221-nt intron was efficiently removed by spliceosome-based constructs but was frequently retained when using the Tornado system **(Figure 3E)**, suggesting that ribozyme cleavage can precede and/or interfere with spliceosome-mediated removal of the internal intron. Constructs containing *ZKSCAN1*, *Laccase2*, and *POLR2A* introns generated comparable levels of scarless circZKSCAN1(2,3) **(Figure 3F** and **Supplementary Figure S9A)**; however, the *POLR2A* introns again produced higher levels of linear RNA byproducts **(Figure 3F** and **Supplementary Figure S9D-E)**.

### Tornado more effectively generates small circRNAs

We previously noted that circRNAs <400 nt in length can be difficult to accurately overexpress from plasmids [32,34], and we therefore examined whether any flanking sequences could provide an effective strategy to remedy this. The endogenous human NFASC gene generates a 270-nt circRNA from exons 26 and 27 (circNFASC(26,27)), and we found that very little circRNA was produced when these exons were cloned between the mini, *POLR2A*, or *ciRS-7* introns **(Supplementary Figure S10)**. Scarless circNFASC(26,27) was produced from introns derived from *ZKSCAN1* and *Laccase2*; however, a ladder of circular RNAs with repetitive exons (circular concatemers [55]) was also generated. In contrast, the Tornado system generated a single prominent circRNA species **(Supplementary Figure S10)**, indicating that it may represent a strategy to more efficiently produce short circRNAs, albeit with the caveat that a scar is present at the junction.

### The extent of linear RNA byproducts is cell-type specific

To assess the generality of circRNA overexpression strategies across cell types, we examined three additional human cell lines: HCT116 colorectal carcinoma **(Figure 4A-C)**, HeLa cervical adenocarcinoma **(Figure 4D-F)**, and SH-SY5Y neuroblastoma **(Figure 4G-I)**. Each cell line was transfected with the split eGFP constructs **(Figure 2A)**, and RNA and protein outputs were analyzed 24 h post-transfection, as had been done for HEK293FT cells. In HCT116 cells, most constructs surprisingly produced comparable levels of eGFP circRNA, apart from the ciRS-7 intron-containing construct, which was markedly less efficient **(Figure 4A-B)**. In HeLa **(Figure 4D-E)** and SH-SY5Y cells **(Figure 4G-H)**, the Tornado system and the construct containing *Laccase2* introns consistently yielded high levels of eGFP circRNA, with Tornado producing a junction scar and the *Laccase2* introns producing scarless circRNA. However, in contrast to HEK293FT cells—where Tornado was approximately twofold more efficient **(Figure 2B-C)**—circRNA levels produced by these two systems were similar in HCT116 **(Figure 4B)**, HeLa **(Figure 4E)**, and SH-SY5Y cells **(Figure 4H)**.

**Figure 4.**
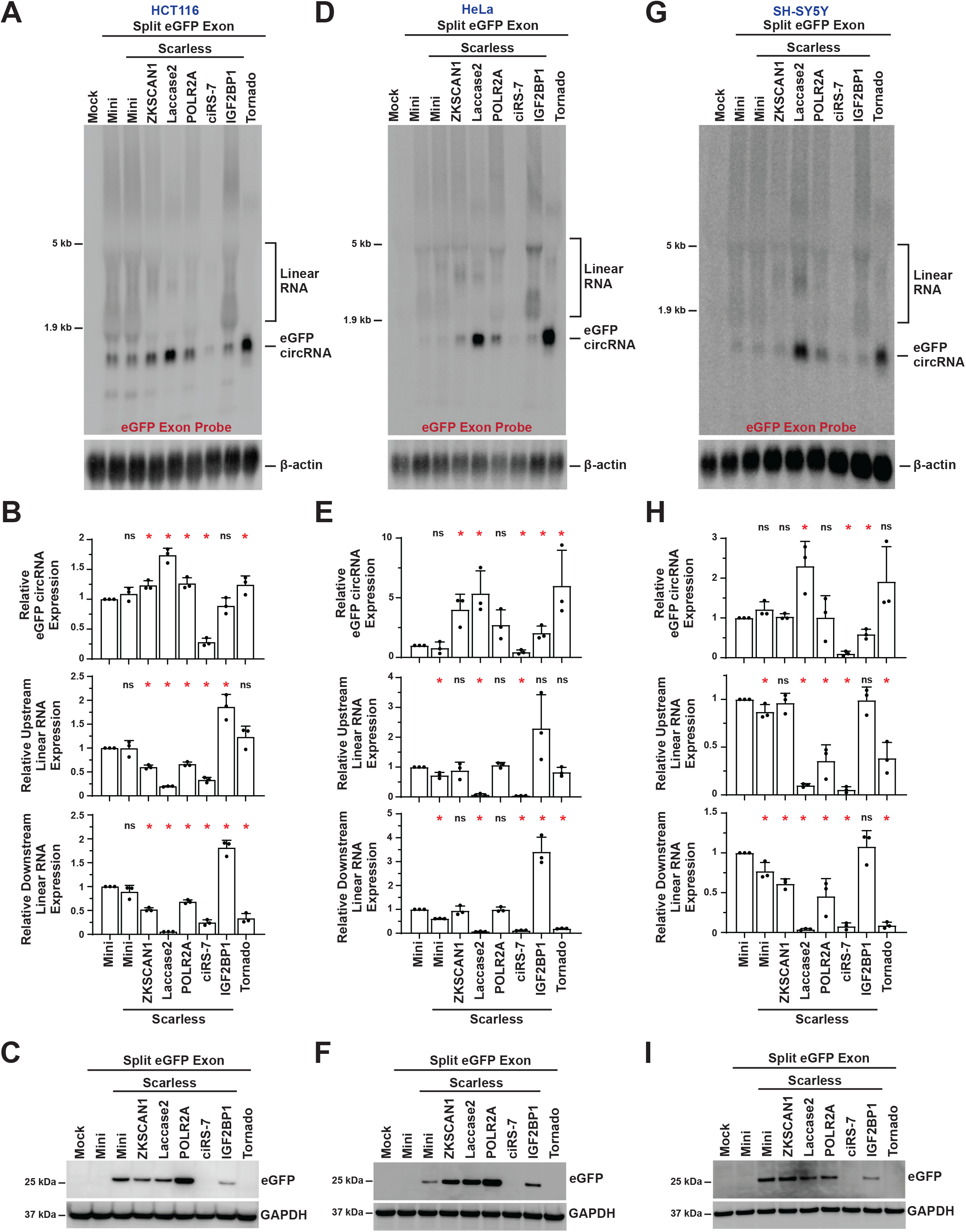
Comparison of circRNA expression strategies across cell types. Plasmids expressing the split eGFP/EMCV IRES exon flanked by the indicated intronic sequences were transfected into HCT116 **(A-C)**, HeLa **(D-F)**, and SH-SY5Y cells **(G-I)**. Total RNA and protein were harvested 24 h post-transfection. **(A, D, G)** 18 μg of total RNA was analyzed by Northern blot using a probe complementary to the eGFP exon. β-actin was used as a loading control. **(B, E, H)** circRNA and linear RNA levels were quantified by RT-qPCR. Data were normalized to the Mini plasmid that produces a scarred BSJ and are presented as mean ± SD (N = 3). p values were generated with unpaired t-tests, comparing each construct to the Mini plasmid that produces a scarred BSJ. *p < 0.05; ns, not significant. Note that the junction scar increases the length of the PCR amplicon for circRNAs derived from the Mini and Tornado plasmids. **(C, F, I)** 20 μg of total protein was analyzed by immunoblotting using an anti-eGFP antibody. GAPDH served as a loading control.

Most notably, the relative abundance of linear RNA byproducts (compared to eGFP circRNA levels) was markedly reduced in all three cell lines compared to HEK293FT cells (compare **Figure 4A, D, G** with **Figure 2B**), indicating that accumulation of linear species is strongly influenced by cellular context. Some degree of linear byproducts remained detectable for most constructs, but those containing *Laccase2* introns consistently produced fewer such species **(Figures 4B, E, H** and **Supplementary Figure S11)**. eGFP protein could further be detected from most of the scarless expression constructs, but, like in HEK293FT cells, eGFP protein expression was not a reliable proxy for circRNA levels **(Figure 4C, F, I)**. Collectively, these findings indicate that linear byproduct accumulation is not an intrinsic, fixed property of a given overexpression strategy but is instead shaped by cellular context, underscoring the importance of empirically validating circRNA purity in each experimental system of interest.

### Linear RNA byproducts can be reduced by destabilizing their 3’ ends

To reduce the accumulation of linear RNA byproducts from circRNA overexpression constructs, we hypothesized that recruitment of endogenous microRNAs [56,57] and/or RNA decay factors could suppress their levels post-transcriptionally. We first focused on constructs containing the *IGF2BP1* introns, which generate high levels of bystander linear RNAs in HEK293FT cells **(Figure 2B-C)**, and introduced perfectly complementary target sites for miR-92a—an abundant microRNA in HEK293FT cells [58]—near the 3’ end, 5’ end, or both ends of the pre-mRNA **(Supplementary Figure S12A)**. This design was intended to enable miR-92a to function in a siRNA-like manner, promoting cleavage of bystander linear RNAs while sparing mature circRNAs. Consistent with this model, inclusion of miR-92a target sites—but not control versions with mutated seed regions—reduced bystander linear RNAs by ∼50% **(Supplementary Figure S12B-C)**. However, circRNA levels were also reduced to a similar degree **(Supplementary Figure S12B-C)**. Because substantial bystander linear byproducts persisted and the target sites could sequester miR-92a, potentially interfering with repression of endogenous targets [59], we next explored an alternative strategy to selectively destabilize linear RNAs from circRNA overexpression constructs.

Specifically, we sought to destabilize the 3’ ends of linear transcripts by replacing the poly(A) signal with mascRNA (MALAT1-associated small cytoplasmic RNA), a 58-nt tRNA-like structure recognized by RNase P that mediates 3’ end processing of the MALAT1 long noncoding RNA [47] **(Figure 5A)**. Following RNase P cleavage, mature MALAT1 is stabilized by a 3’ triple-helix structure, and disruption of this structure promotes rapid decay of the long noncoding RNA [48,60,61]. Based on this, we inserted mouse mascRNA—without the stabilizing MALAT1 triple helix—at the 3’ end of the circRNA expression constructs **(Figure 5A)** and measured RNA levels 24 h post-transfection in HEK293FT cells. Importantly, mouse mascRNA contains four sequence differences from the human homolog, thereby allowing plasmid-derived and endogenous mascRNA transcripts to be distinguished by Northern blot [48]. Replacing the poly(A) signal with wildtype (WT) mouse mascRNA significantly reduced linear bystander RNAs, particularly transcripts retaining downstream intron sequences, which were no longer detectable by RT-qPCR, even from constructs containing the *IGF2BP1* introns **(Figure 5B-C** and **Supplementary Figure S13)**. CircRNA levels were also reduced (often by more than 50%); however, the substantially greater reduction in linear byproducts improved the overall circRNA-to-linear RNA ratio.

Although mascRNA insertion reduced linear RNA levels, it also resulted in accumulation of mouse mascRNA itself as a byproduct **(Figure 5B)**. To address this, we introduced mutations into the mouse mascRNA acceptor stem designed to destabilize the RNA and promote its post-transcriptional degradation, either through uridylation (mascRNA Mut 7) or CCACCA addition to the 3’ end by the CCA-adding enzyme (mascRNA Mut 10) [48,62,63] **(Figure 5A)**. These mutant mouse mascRNA transcripts minimally accumulated in cells, but the output of the plasmids otherwise largely mirrored that seen with the wildtype mascRNA sequence **(Figure 5B-C)**.

Together, these results indicate that 3’ end engineering can strongly reduce linear RNA byproducts while still allowing circRNA expression. More broadly, these approaches highlight the importance of post-transcriptional RNA stability in shaping the outputs of circRNA expression systems, offering a generalizable strategy that could be readily adapted to other overexpression platforms where linear byproduct contamination limits interpretability or therapeutic utility.

### A strategy for bicistronic expression of circRNA and a selectable marker mRNA

The strategies pursued thus far have focused on expressing circRNAs alone. However, in some applications, coordinated expression of a linear mRNA and a circRNA is desirable. For example, co-expression of a fluorescent reporter enables tracking and isolation of circRNA-expressing cells. To address this, we designed a bicistronic expression cassette (CIRCUS, <u>circ</u>RNA and <u>u</u>pstream linear <u>s</u>ystem) that generates both a linear mRNA and a circRNA from a single CMV promoter **(Figure 6A)**. This construct first produces a translatable linear mRNA by repurposing the MALAT1 3’ end processing mechanism. Specifically, mouse mascRNA and the upstream triple-helix structure are used to generate a stable, non-polyadenylated transcript that is efficiently exported to the cytoplasm and translated [47,48,60,61]. Downstream of this module, a circRNA expression cassette is positioned, in which circularization is driven either by the spliceosome (using *ZKSCAN1* or *Laccase2* introns) or by the Tornado system. As in **Figure 5**, either wildtype mouse mascRNA **(Supplementary Figure S14)** or a destabilized mutant (Mut 7) **(Figure 6A)** was included to modulate mascRNA accumulation.

**Figure 5.**
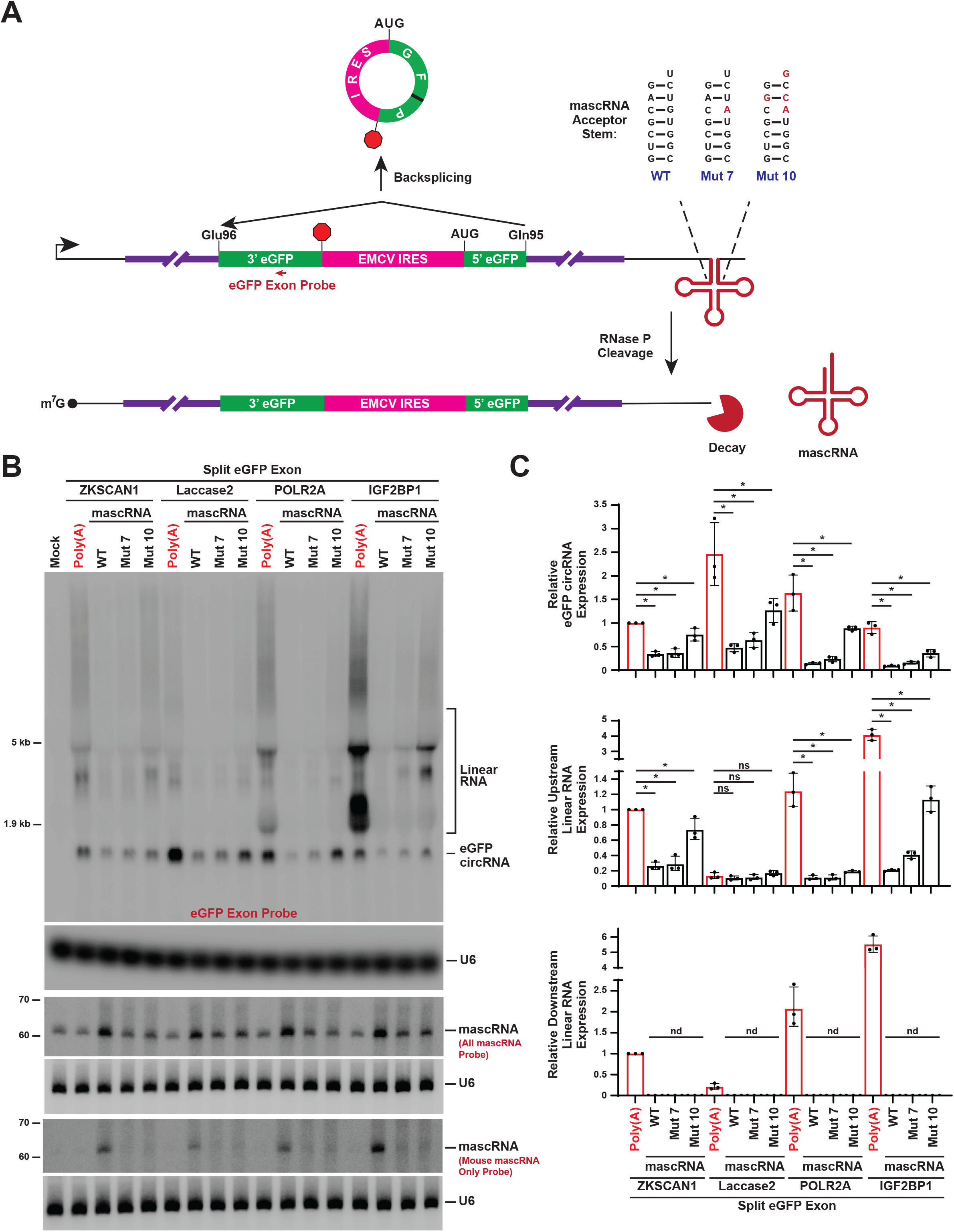
Destabilization of the pre-mRNA 3’ end reduces bystander linear RNA levels. **(A)** The split eGFP/EMCV IRES exon expression plasmids from Figure 2A were altered by replacing the downstream poly(A) signal with mouse mascRNA. This tRNA-like structure can be recognized by RNase P, resulting in production of a mature linear RNA susceptible to decay due to its unprotected 3’ end. In addition to wildtype (WT) mascRNA, variants with the indicated mutations in the acceptor stem (naming scheme as per [62]) were tested. **(B-C)** Plasmids expressing the split eGFP/EMCV IRES exon between the indicated intronic sequences and terminating in a poly(A) signal or a mascRNA variant were transfected into HEK293FT cells, and total RNA was isolated after 24 h. **(B)** 18 μg of total RNA was analyzed by Northern blots using probes complementary to the eGFP exon and mascRNA. β-actin and U6 snRNA were used as loading controls. **(C)** RT-qPCR was used to quantify circRNA and linear RNA expression from each plasmid. Data are normalized to the plasmid containing ZKSCAN1 introns and a poly(A) signal and presented as mean ± SD (N = 3). p values were generated with unpaired t-tests, comparing each construct to the equivalent plasmid ending in a poly(A) signal. *p < 0.05; ns, not significant; nd, not detectable.

**Figure 6.**
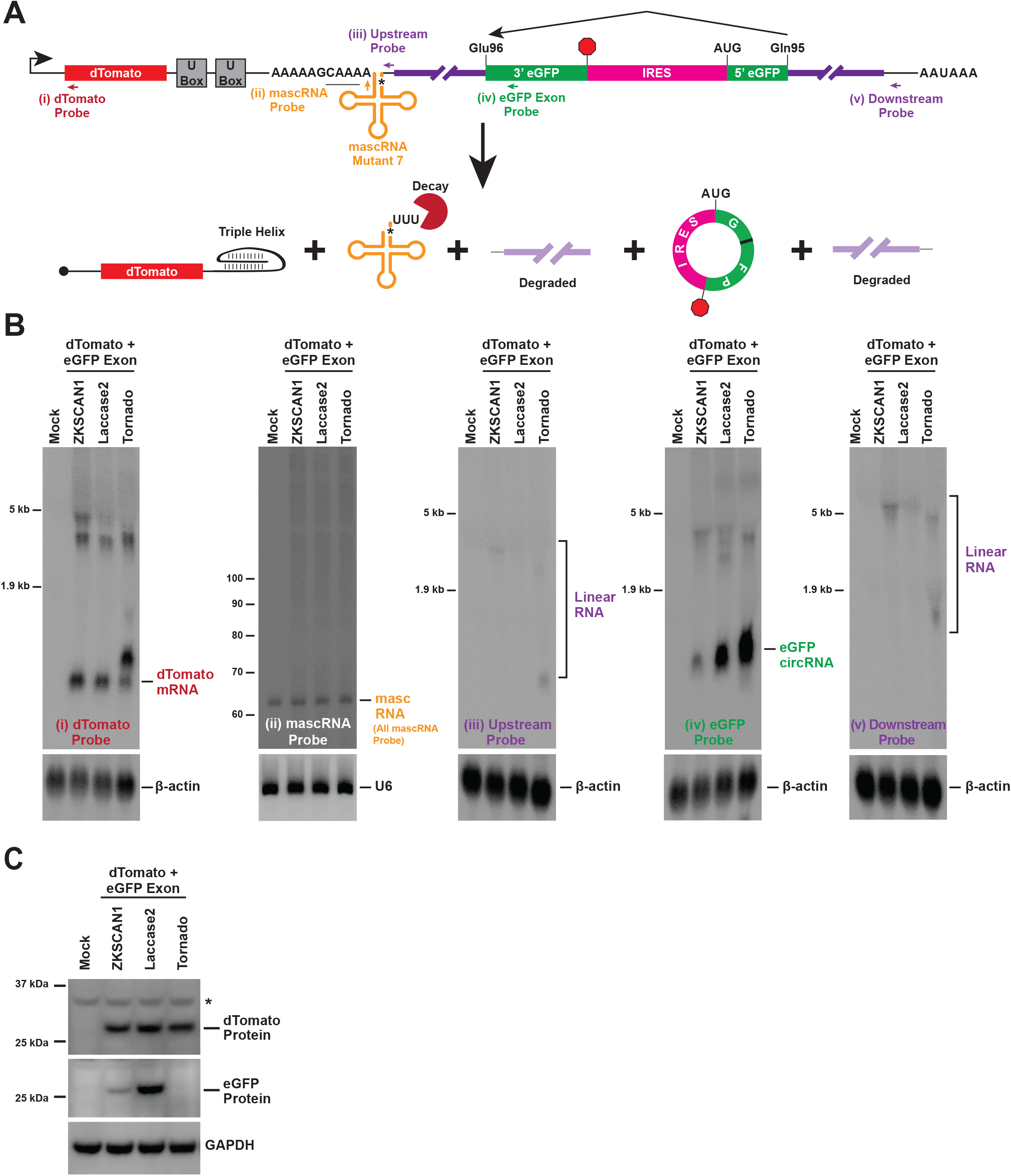
Coordinated expression of a linear mRNA and a circRNA from a single promoter. **(A)** Schematic of the CIRCUS platform enabling co-expression of a linear mRNA encoding dTomato with a circRNA encoding eGFP. RNase P cleavage enables production of a stable dTomato linear mRNA ending in the MALAT1 triple helix, while the mutant mascRNA transcript is targeted for post-transcriptional decay following RNase Z cleavage. The intronic sequence pairs (purple) facilitate production of the eGFP circRNA, before themselves being targeted for degradation. Oligonucleotide probes used for Northern blotting are indicated. **(B)** The CIRCUS plasmids with the indicated sequences flanking the split eGFP/EMCV IRES exon were transfected into HEK293FT cells, and total RNA was isolated after 24 h. 18 μg of total RNA was analyzed by Northern blots with the indicated probes. β-actin and U6 snRNA were used as loading controls. **(C)** Total protein was likewise isolated 24 h post-transfection. 20 μg of total protein was analyzed by immunoblotting using an anti-eGFP antibody. GAPDH was used as a loading control. *, non-specific band.

To evaluate the CIRCUS platform, we first aimed to co-express a linear mRNA encoding dTomato with a circRNA encoding eGFP **(Figure 6A** and **Supplementary Figure S14)**. At 24 h post-transfection in HEK293FT cells, Northern blot analysis confirmed robust expression of the dTomato mRNA terminating in the MALAT1-derived triple helix **(Figure 6B)**, accompanied by dTomato protein production **(Figure 6C)**. The eGFP circRNA was also efficiently generated, with Tornado producing higher levels than spliceosome-mediated approaches **(Figure 6B)** but no eGFP protein due to the junction scar **(Figure 6C)**, consistent with earlier results **(Figure 2B-C)**. Wildtype mascRNA accumulated as expected, confirming proper processing **(Supplementary Figure S14B)**, whereas the mascRNA Mut 7 variant was rapidly degraded following processing **(Figure 6B)**, thereby enabling an overall cleaner expression system.

Notably, the CIRCUS construct containing *Laccase2* introns showed strongest eGFP protein expression **(Figure 6C)** along with minimal accumulation of upstream or downstream intronic regions **(Figure 6B** and **Supplementary Figure S14D)**, indicating that the intended linear mRNA and circRNA species predominate. When using Tornado, mascRNA cleavage was less efficient, resulting in two dTomato mRNA isoforms: the intended triple-helix-terminated transcript and a less abundant, longer species terminating at the first ribozyme cleavage site **(Figure 6B**, dTomato blot). Similar levels of dTomato protein were nonetheless detected regardless of the sequences used to drive downstream circRNA production **(Figure 6C)**. Collectively, these findings demonstrate that pairing a linear reporter with an optimized circRNA expression cassette, separated by a destabilized mascRNA structure, allows both products to be generated efficiently and with minimal byproduct accumulation, addressing a key limitation of existing circRNA overexpression tools, which lack a built-in means of tracking or enriching expressing cells.

### Use of the CIRCUS platform to mark cells with high efficiency of site-specific A-to-I editing

Beyond expressing proteins, circRNAs can be designed to modulate microRNA activity [23,24], function as aptamers [25,26], or serve as guide RNAs for site-specific DNA or RNA editing [27–29]. To test the utility of the CIRCUS platform for tracking and isolating cells that have been successfully modulated by a circRNA of interest, we replaced the eGFP-encoding circRNA from **Figure 6** with a circular guide RNA designed to recruit endogenous ADAR (adenosine deaminase acting on RNA) for site-specific adenosine-to-inosine (A-to-I) editing **(Figure 7A)**. ADAR can hydrolytically deaminate an A within a region of double-stranded RNA, converting it to an I that is biochemically interpreted as a G by many cellular processes [64].

**Figure 7.**
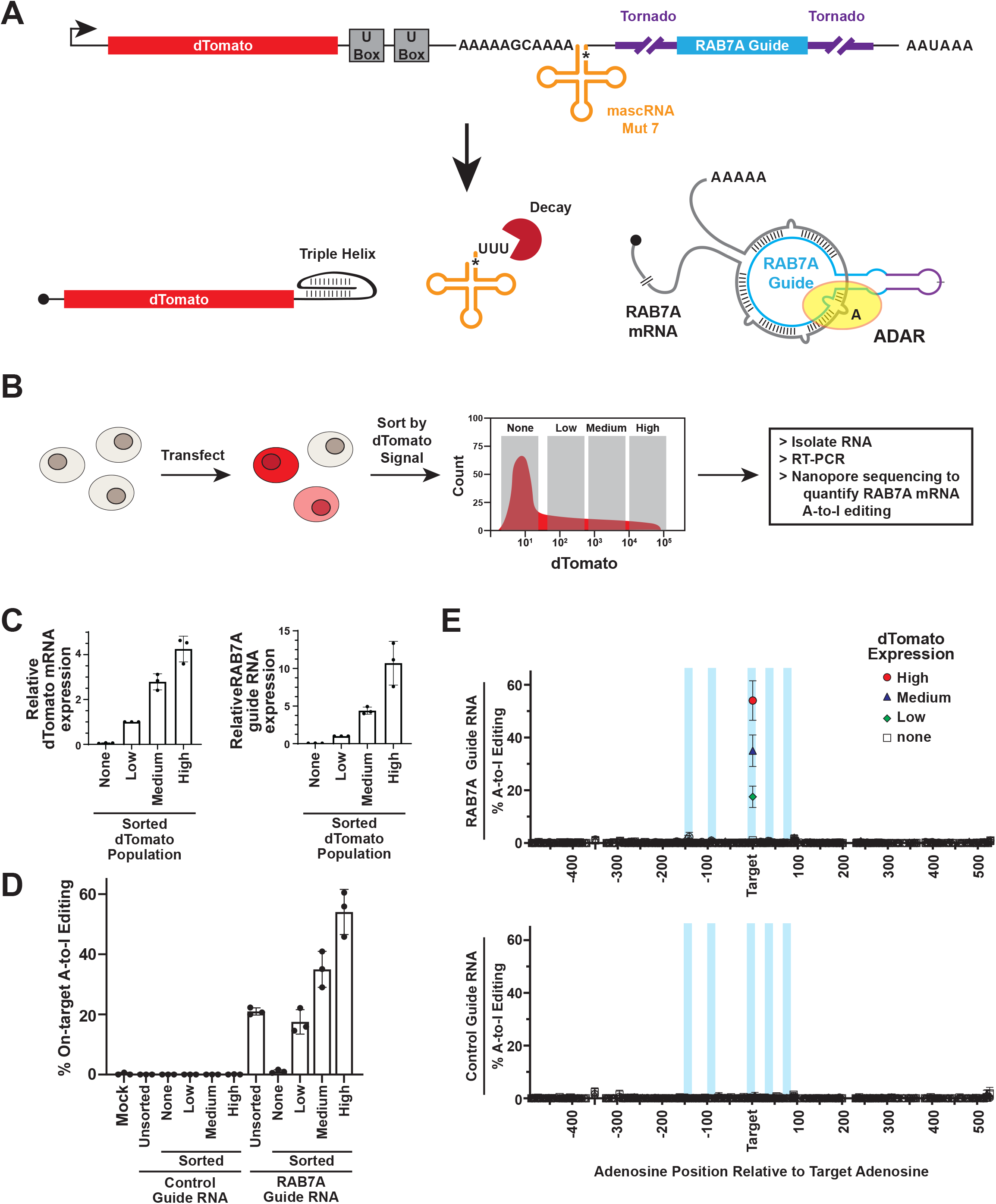
The CIRCUS platform enables identification of cells with high circRNA-driven, site-specific A-to-I editing. **(A)** Schematic of the CIRCUS platform enabling co-expression of a linear mRNA encoding dTomato with a circRNA that directs site-specific A-to-I editing of the RAB7A 3’ UTR. RNase P cleavage enables production of a stable dTomato linear mRNA ending in the MALAT1 triple helix, while the mutant mascRNA transcript (orange) is targeted for post-transcriptional decay following RNase Z cleavage. The Tornado platform facilitates production of a circRNA (blue) with multiple regions of complementarity to the 3’ UTR of RAB7A mRNA, enabling ADAR recruitment and site-specific A-to-I editing. **(B)** HEK293FT cells were transfected with the CIRCUS platform expressing the RAB7A guide RNA or a control guide RNA, and cells were sorted after 24 h into four bins based on dTomato signal (none, low, medium, and high). Total RNA was isolated from each fraction, reverse transcribed, and the 3’ UTR of RAB7A was PCR amplified and subjected to Nanopore sequencing to quantify A-to-I editing events. **(C)** RT-qPCR was used to quantify expression levels of dTomato mRNA and the RAB7A guide RNA in each sorted cell population. **(D)** The level of A-to-I edits at the target adenosine in the 3’ UTR of RAB7A mRNA, as determined by A-to-G sequence changes, in unsorted or sorted cells transfected with the CIRCUS platform expressing the RAB7A guide RNA or a control guide RNA. **(E)** The level of A-to-I edits at all adenosines present in the RAB7A 3’ UTR amplicon. Light blue shading indicates regions of complementarity between the RAB7A guide RNA and its target. Full Nanopore sequencing data are provided in **Supplementary Table S4.**

There has thus been increasing therapeutic interest in inducing specific A-to-I editing events through purposeful expression of guide RNAs complementary to a transcript of interest [65]. Here, we used the CLUSTER approach [28,66] to design a circRNA complementary to the 3’ untranslated region (UTR) of the RAB7A mRNA, aiming to convert a single A to an I. This circular guide binds RAB7A in a multivalent fashion: there are five distinct regions (20-nt each) with complementarity to the RAB7A 3’ UTR, with an A:C mismatch at the desired editing site, consistent with the preferred substrate configuration for ADAR [67] **(Figure 7A** and **Supplementary Figure S15A)**. In addition, the circRNA contains an ADAR-recruiting domain (R/G motif) modified from a well-characterized editing site in the GluR2 (GluR-B) transcript [68] to increase editing efficiency. Given the small size (220 nt) of the RAB7A guide RNA, we chose to employ the Tornado approach to drive its production based on the results in **Supplementary Figure S10**.

As the CIRCUS platform generates both dTomato and this circRNA **(Figure 7A)**, we reasoned that the efficiency of A-to-I editing within the RAB7A 3’ UTR should positively correlate with dTomato expression levels. We thus transfected HEK293FT cells with a CIRCUS construct expressing the RAB7A guide RNA or a control guide RNA and used fluorescence-activated cell sorting (FACS) to sort cells after 48 h into four bins based on dTomato fluorescence signal (none, low, medium, and high) **(Figure 7B** and **Supplementary Figure S15B)**. Unsorted cells were also retained for comparison. Total RNA was isolated from each cell population, and we first used RT-qPCR to quantify expression of dTomato mRNA and the RAB7A guide RNA **(Figure 7C)**. As expected, little dTomato mRNA or guide RNA was detected in cells lacking dTomato fluorescence, but their levels progressively increased with dTomato fluorescence intensity. RT-PCR was then used to amplify the 3’ UTR of RAB7A, and the products were subjected to Nanopore sequencing to quantify A-to-I editing events (as measured by A-to-G mutations). The control guide RNA resulted in no on-target RNA editing, whereas the RAB7A guide resulted in ∼20% editing in the unsorted population **(Figure 7D)**. Upon examining cells sorted by dTomato fluorescence, a progressive increase in on-target editing was observed, with A-to-I editing levels reaching nearly 60% in cells expressing the highest levels of dTomato— representing an ∼3-fold enrichment in editing efficiency achieved simply by sorting for reporter expression. This suggests that A-to-I editing efficiency scales with circular guide RNA abundance. Notably, editing was highly site-specific, with negligible editing of other adenosines within a ∼1 kb region of the RAB7A 3’ UTR flanking the target editing site **(Figure 7E** and **Supplementary Table S4)**. Together, these results demonstrate that dTomato fluorescence serves as a reliable, real-time proxy for circRNA-guided A-to-I editing efficiency, enabling enrichment of highly edited cells by simple fluorescence-based sorting rather than laborious clonal screening. More broadly, this approach establishes a generalizable strategy for isolating or tracking cells with high activity of a circRNA of interest, applicable to potentially any phenotype (editing, translation, or otherwise) that is driven by circRNA expression from the CIRCUS platform.

## DISCUSSION

CircRNAs have been increasingly investigated and deployed both as experimental tools and therapeutic candidates, yet the field has largely lacked a systematic, quantitative comparison of the overexpression strategies on which this work depends. Here, we benchmarked widely used strategies for circRNA overexpression in human cells and developed an improved toolkit that substantially reduces contaminating linear RNA byproducts while preserving efficient circRNA production. Our results reveal that no single overexpression strategy is universally optimal: circRNA yield, junction scarring, and linear byproduct accumulation all vary substantially across constructs, underscoring the importance of empirically validating circRNA purity in any given experimental system rather than assuming the results obtained with one platform or in one cell type will generalize to another.

Among the strategies tested, the ribozyme-based Tornado system consistently produced the highest circRNA yields and proved particularly effective for generating small circRNAs that are otherwise difficult to overexpress efficiently. However, this approach comes with two notable drawbacks: the introduction of a 47-nt junction scar into the mature circRNA precludes use in applications requiring a native, unmodified circRNA junction (e.g., studying endogenous circRNA-derived protein products) and the upstream cleavage fragment accumulates due to its 3’ end being protected by a 2’,3’-cyclic phosphate **(Figure 2)**. By contrast, spliceosome-mediated circularization using introns containing imperfect complementary repeats—most notably those derived from the *Drosophila Laccase2* gene—produced scarless circRNA with a favorable balance of yield and purity across multiple exons and cell types examined **(Figures 2-4)**. Interestingly, we found that extensive perfect complementarity between flanking repeats, as used in multiple widely adopted constructs [7,33,35], may limit backsplicing efficiency, independent of splice site strength. This observation suggests that repeat architecture, not merely repeat presence, is a key design parameter for circRNA overexpression tools.

Our work further highlights that circRNA overexpression constructs can generate substantial quantities of unintended linear RNA species—including unspliced pre-mRNAs and trans-spliced exon concatemers—that are easily overlooked depending on the detection strategy employed. This has direct implications for interpreting circRNA-driven phenotypes: without explicit characterization of these byproducts, effects attributed to a circRNA of interest may instead reflect the activity of a contaminating linear species, a concern that has fueled ongoing debate in the field, especially regarding whether circRNAs are translated [36–42]. Given that even low-abundance linear contaminants may be efficiently translated via canonical cap-dependent mechanisms, we suggest that protein products derived from circRNA overexpression plasmids should be interpreted cautiously. Our finding that linear byproduct accumulation is strongly cell-type dependent **(Figures 2 and 4)** further complicates this picture, as a construct validated as “clean” in one cell line cannot be assumed to behave identically in another.

To reduce these linear byproducts, we engineered the primary transcript to terminate in a non-polyadenylated 3’ end using mascRNA, a tRNA-like structure recognized by RNase P [47], rendering unwanted linear RNAs substantially more susceptible to exonucleolytic decay **(Figure 5)**. Further engineering of the mascRNA acceptor stem to promote its own post-transcriptional degradation minimized accumulation of the tRNA-like small RNA [48,62], yielding an overall cleaner expression system. This mascRNA-based strategy is mechanistically distinct from microRNA-mediated cleavage of linear transcripts [56,57], though the two approaches are complementary. Combining both approaches may help further reduce linear RNAs from circRNA overexpression plasmids, especially from the Tornado platform where the 2’,3’-cyclic phosphate present on the end of the upstream linear RNA limits its decay by exonucleases.

Building on our optimized systems for expressing a circRNA, we further developed CIRCUS, a bicistronic platform that co-expresses a linear fluorescent reporter alongside a circRNA of interest from a single promoter **(Figure 6)**. This design addresses a practical limitation of existing circRNA overexpression tools, which generally lack a built-in means of identifying or enriching cells that express the circRNA at high levels. Using CIRCUS to drive expression of an ADAR-recruiting circular guide RNA, we found that reporter fluorescence intensity correlated strongly with on-target A-to-I editing efficiency **(Figure 7)**. Because this relationship reflects the shared transcriptional origin of the reporter and the circRNA, this strategy should be generalizable to other circRNA-driven phenotypes, including microRNA sponging [23,24] or aptamer-based signaling [25,26], providing a broadly applicable tool for enriching or tracking cells with high circRNA activity. In addition, the fluorescent reporter ORF can easily be swapped with any sequence of interest, thereby allowing bicistronic expression of nearly any linear and circRNA combination from a single promoter, saving space in vectors like adeno-associated virus (AAV) where cargo packaging capacity is at a premium [69].

Several limitations should be noted. Our comparisons were performed using transient plasmid transfection in a limited set of human cell lines, and extrapolation to other expression systems, primary cells, or in vivo delivery contexts will require further validation. Additionally, although mascRNA-based 3’ end engineering substantially reduced linear byproducts, some residual contamination persisted across all constructs tested, and further optimization may be needed for applications demanding the highest achievable purity. Collectively, this work provides a systematically benchmarked and optimized toolkit for circRNA overexpression in human cells, offering improved tools for both dissecting circRNA biology and advancing circRNA-based therapeutic development.

## ACKNOWLEDGMENTS

We thank all members of the Wilusz and Conn labs as well as Bill Lagor for discussions and advice. This project was supported by the Cytometry and Cell Sorting Core at Baylor College of Medicine and the assistance of Joel M. Sederstrom.

## AUTHOR CONTRIBUTIONS

C.J.F., S.J.C., and J.E.W. conceived and designed the project. B.W.S. and S.J.C. conceptualized the idea of the scarless circRNA expression strategy. C.J.F., R.F., B.W.W., and B.W.S. performed experiments and analyzed data. C.J.F. and J.E.W. wrote the manuscript with input from all authors.

## DATA AVAILABILITY

All data supporting this study are available within the figures and supplementary data files.

