## Supplementary Figures, Methods, and Tables S1-3 for "Engineering circular RNA expression systems to minimize contaminating linear RNA byproducts"

#### Figure S1

**A**

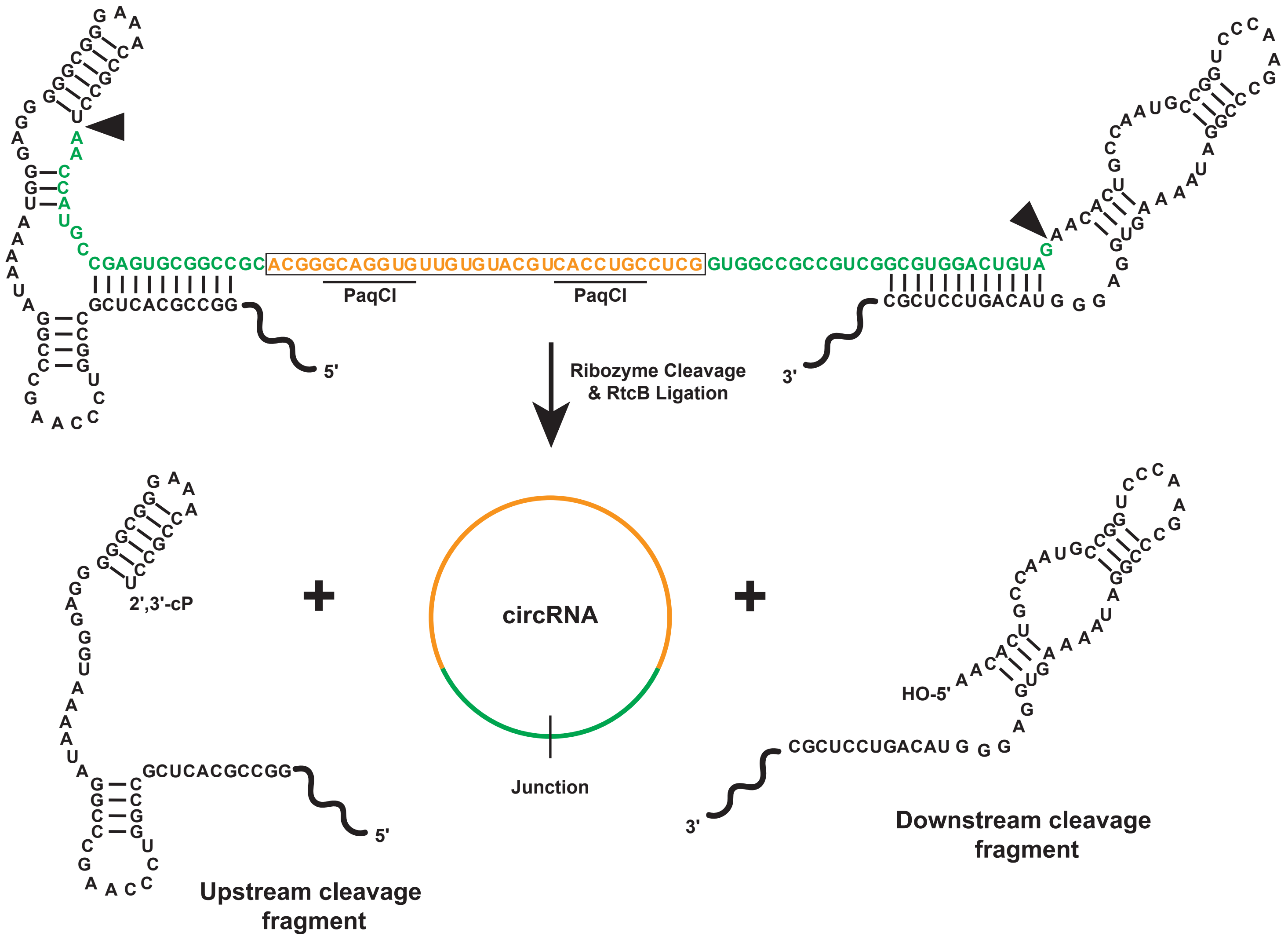

CGAGUGCGGCCGC ACGGGCAGGUGUUGUGUACGUCACCUGCCUCG GUGGCCGCCGUCGGCGUGGACUGUA
GCUCACGCCGG
CGCUCCUGACAU G G G
PaqCI
PaqCI
5'
3'
Ribozyme Cleavage & RtcB Ligation
2',3'-cP
circRNA
Junction
HO-5'
Upstream cleavage fragment
Downstream cleavage fragment


**B**

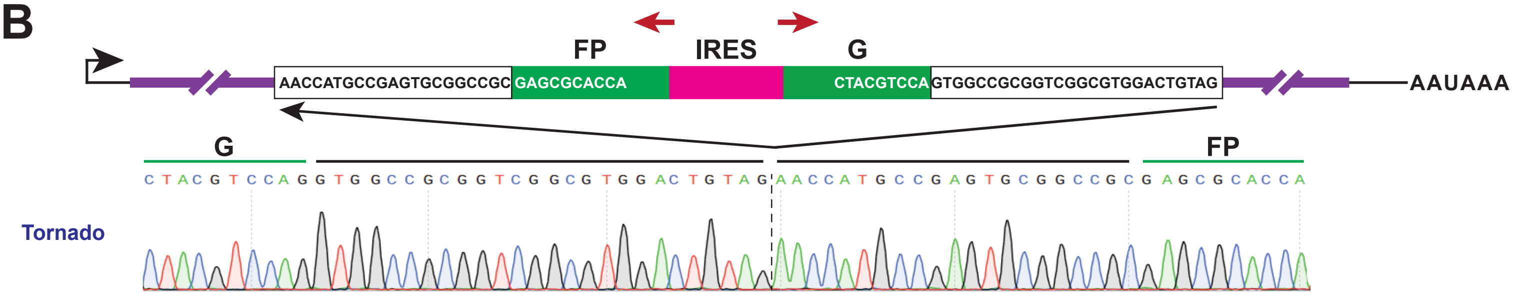

FP
IRES
G
AACCATGCCGAGTGCGGCCGC
GAGCGCACCA
CTACGTCCA
GTGGCCGCGGTCGGCGTGGACTGTAG
AAUAAA
G
FP
CTACGTCCAGGTGGCCGCGGTCGGCGTGGACTGTAGAACCATGCCGAGTGCGGCCGCGAGCGCACCA
Tornado

**Supplementary Figure S1. The Tornado system generates circRNAs with a 47-nt sequence “scar”.**

**(A)** A sequence of interest can be inserted between two Twister ribozymes using PaqCI sites. Each ribozyme self-cleaves at the indicated arrowhead, producing termini (a 5' hydroxyl and a 2',3'-cyclic phosphate [2',3'-cP]) that are compatible with ligation by the endogenous RtcB ligase. This process yields a mature circRNA containing a 47-nt “scar” (green) at the junction, along with upstream and downstream linear fragments.

**(B)** An exon encoding a split eGFP ORF together with the EMCV IRES was cloned into the PaqCI sites of the Tornado construct and transfected into HEK293FT cells. Total RNA was isolated, reverse transcribed using random hexamers, and PCR was performed with the indicated forward and reverse primers (red). PCR products were analyzed by Sanger sequencing. As expected, the resulting mature circRNA contained a 47-nt “scar” at the junction that disrupts the eGFP ORF.

### Figure S2

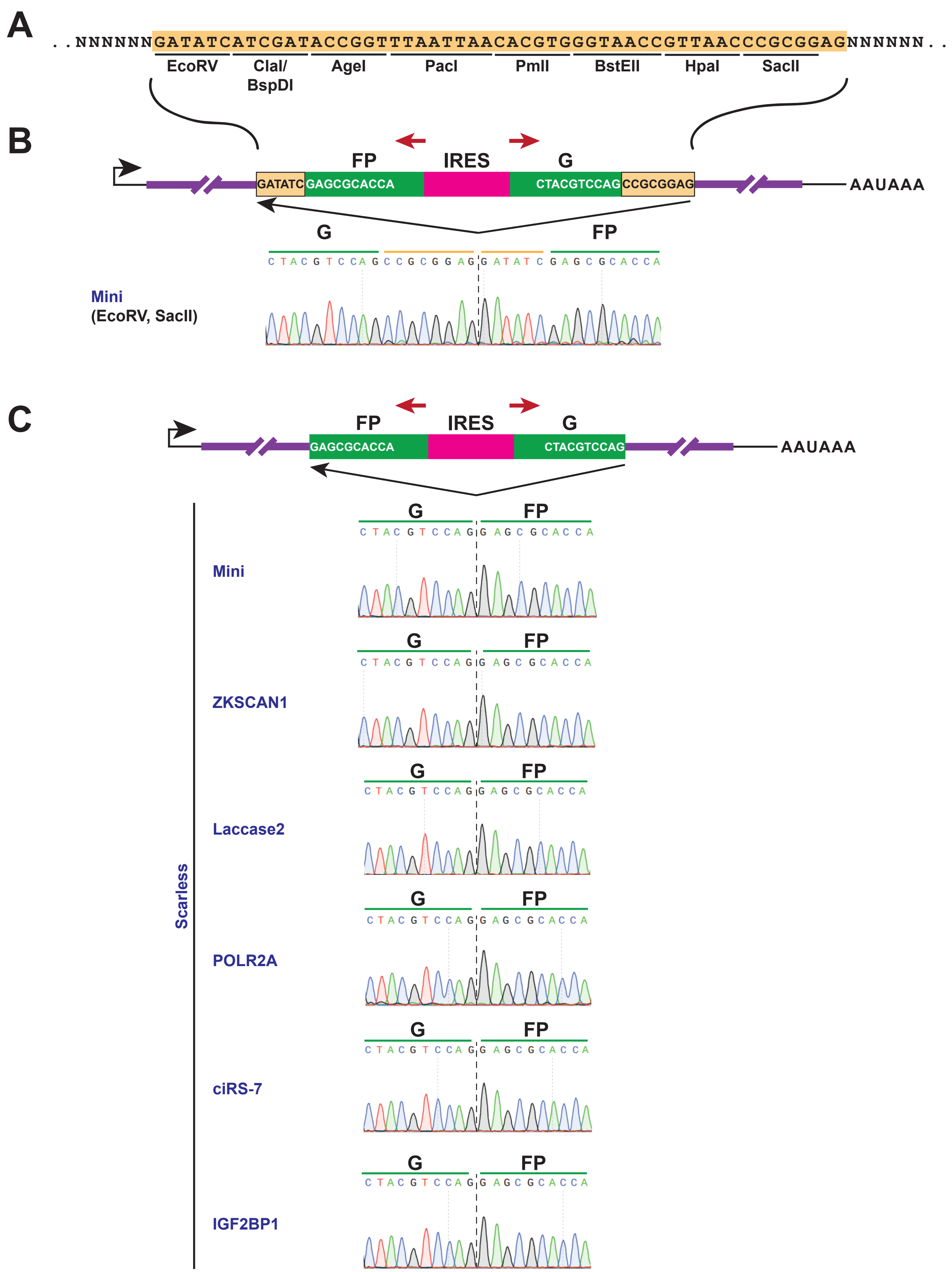

A
. . NNNNNNGATATCATCGATACCGGTTTAATTAACACGTGGGTAACCGTTAACCCGCGGAGNNNNNN . .
EcoRV
ClaI/ BspDI
AgeI
PacI
PmlI
BstEII
HpaI
SacII
B
FP
IRES
G
GATATC GAGCGCACCA
CTACGTCCAG CCGCGGAG
AAUAAA
G
FP
C T A C G T C C A G C C G C G G A G G A T A T C G A G C G C A C C A
Mini
(EcoRV, SacII)
C
FP
IRES
G
GAGCGCACCA
CTACGTCCAG
AAUAAA
Scarless
Mini
ZKSCAN1
Laccase2
POLR2A
ciRS-7
IGF2BP1
C T A C G T C C A G G A G C G C A C C A

**Supplementary Figure S2. Precise production of scarred or scarless circRNA backsplice junctions depending on the cloning vector employed.**

**(A)** The previously described CircRNA Mini Vector ("Mini") contains an artificial 51-nt exon (orange) composed of restriction enzyme sites flanked by minimal introns from the human *ZKSCAN1* gene [32].

**(B)** An exon encoding a split eGFP ORF together with the EMCV IRES was cloned between the EcoRV and SacII sites of the Mini construct and transfected into HEK293FT cells. Total RNA was isolated, reverse transcribed using random hexamers, and PCR was performed with the indicated forward and reverse primers (red). PCR products were analyzed by Sanger sequencing. As expected, the resulting mature circRNA contained a 14-nt "scar" (derived from the restriction sites) that disrupts the eGFP ORF at the backsplice junction.

**(C)** The split eGFP/EMCV IRES exon was cloned analogously into the PaqCI sites of the indicated circRNA overexpression plasmids shown in **Figure 1B** and transfected into HEK293FT cells. RT-PCR confirmed precise backsplicing, with no sequence scar at the backsplice junction.

### Figure S3

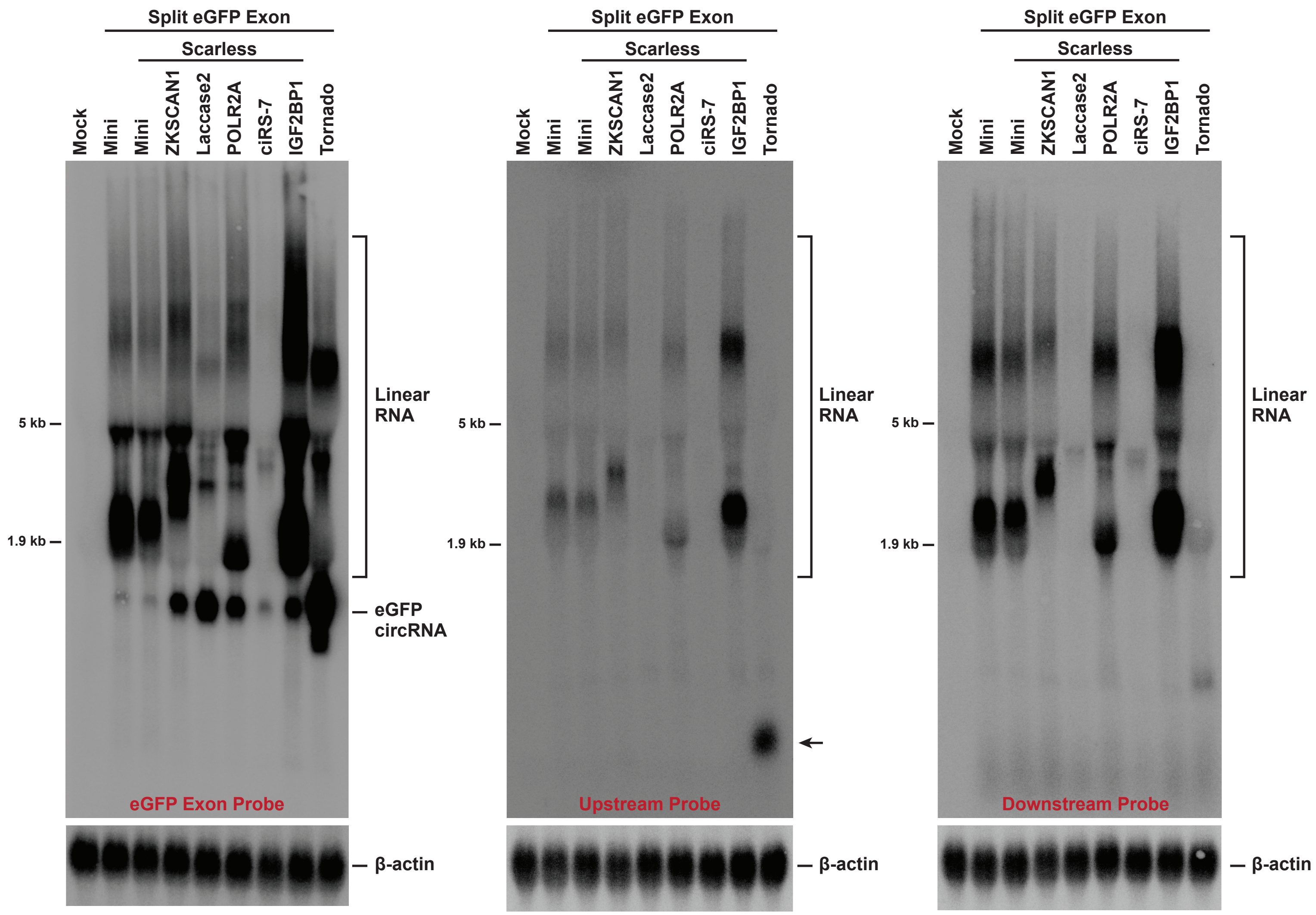
Split eGFP Exon
Scarless
Mock
Mini
Mini
ZKSCAN1
Laccase2
POLR2A
ciRS-7
IGF2BP1
Tornado
5 kb
1.9 kb
Linear RNA
eGFP circRNA
eGFP Exon Probe
Upstream Probe
Downstream Probe
β-actin

**Supplementary Figure S3. High-contrast images of Northern blots from Figure 2.**

Gel images from Figures 2B, 2F, and 2G are shown at high contrast to facilitate visualization of less abundant transcripts. β-actin was used as a loading control, and these images are identical to those shown in Figure 2.

### Figure S4

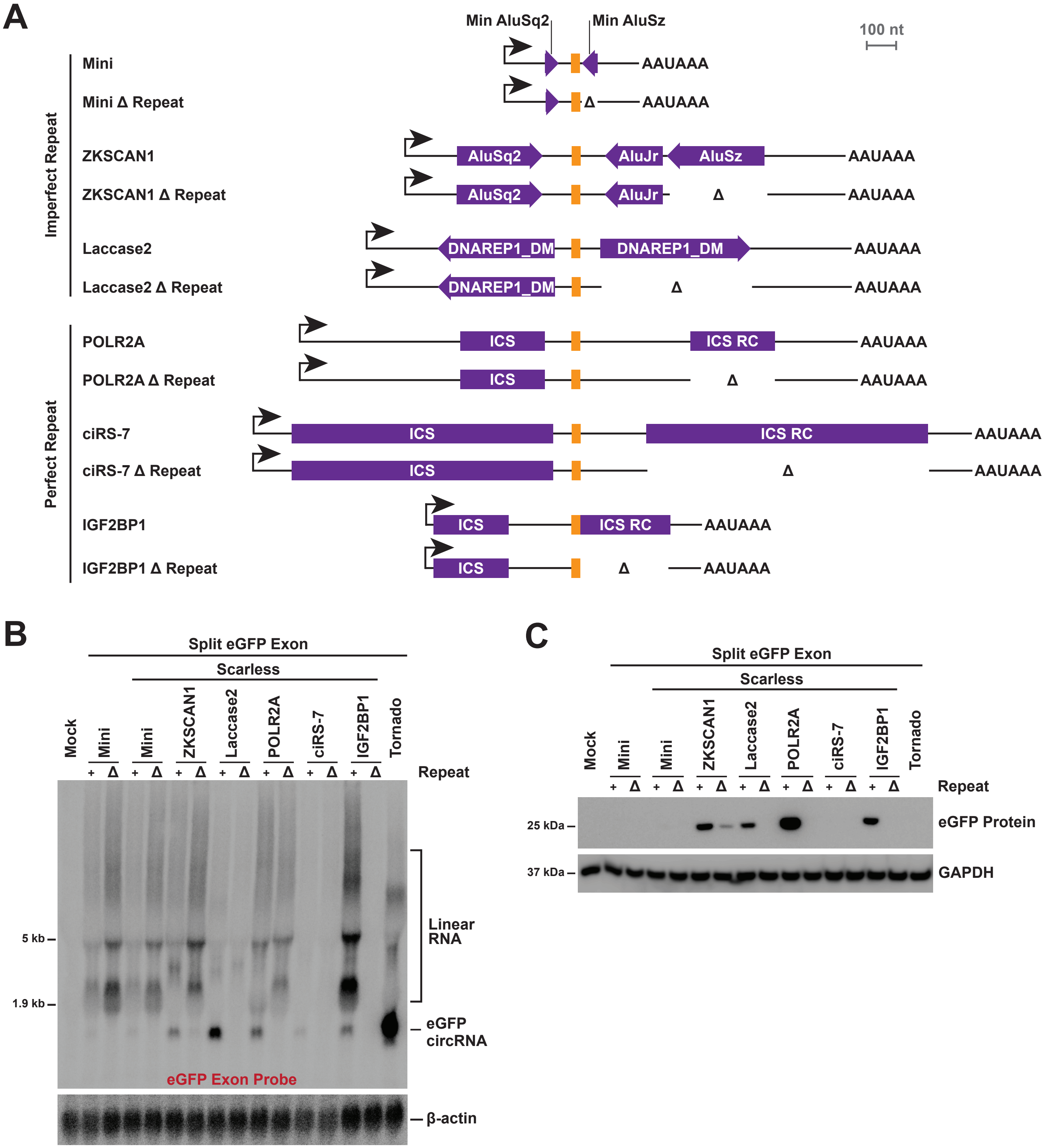
A
Min AluSq2
Min AluSz
100 nt
Imperfect Repeat
Mini
Mini Δ Repeat
ZKSCAN1
ZKSCAN1 Δ Repeat
Laccase2
Laccase2 Δ Repeat
AluSq2
AluJr
AluSz
DNAREP1_DM
AAUAAA
Δ
Perfect Repeat
POLR2A
POLR2A Δ Repeat
ciRS-7
ciRS-7 Δ Repeat
IGF2BP1
IGF2BP1 Δ Repeat
ICS
ICS RC
B
Split eGFP Exon
Scarless
Mock
Mini
Mini
ZKSCAN1
Laccase2
POLR2A
ciRS-7
IGF2BP1
Tornado
+ Δ
Repeat
5 kb
1.9 kb
Linear RNA
eGFP circRNA
eGFP Exon Probe
β-actin
C
Split eGFP Exon
Scarless
Mock
Mini
Mini
ZKSCAN1
Laccase2
POLR2A
ciRS-7
IGF2BP1
Tornado
+ Δ
Repeat
25 kDa
eGFP Protein
37 kDa
GAPDH

**Supplementary Figure S4. Deletion of flanking repeat sequences influences backsplicing efficiency.**

**(A)** Deletions were introduced into each expression plasmid to remove the repeat sequence from the downstream intron.

**(B)** The split eGFP/EMCV IRES exon was cloned into each of the indicated plasmids and transfected into HEK293FT cells. “+” and “Δ” indicate the presence or absence of the downstream repeat sequence, respectively. 18 μg of total RNA was analyzed by Northern blot using a probe complementary to the eGFP ORF. β-actin was used as a loading control.

**(C)** 20 μg of total protein was analyzed by immunoblotting using an anti-eGFP antibody. GAPDH was used as a loading control.

### Figure S5

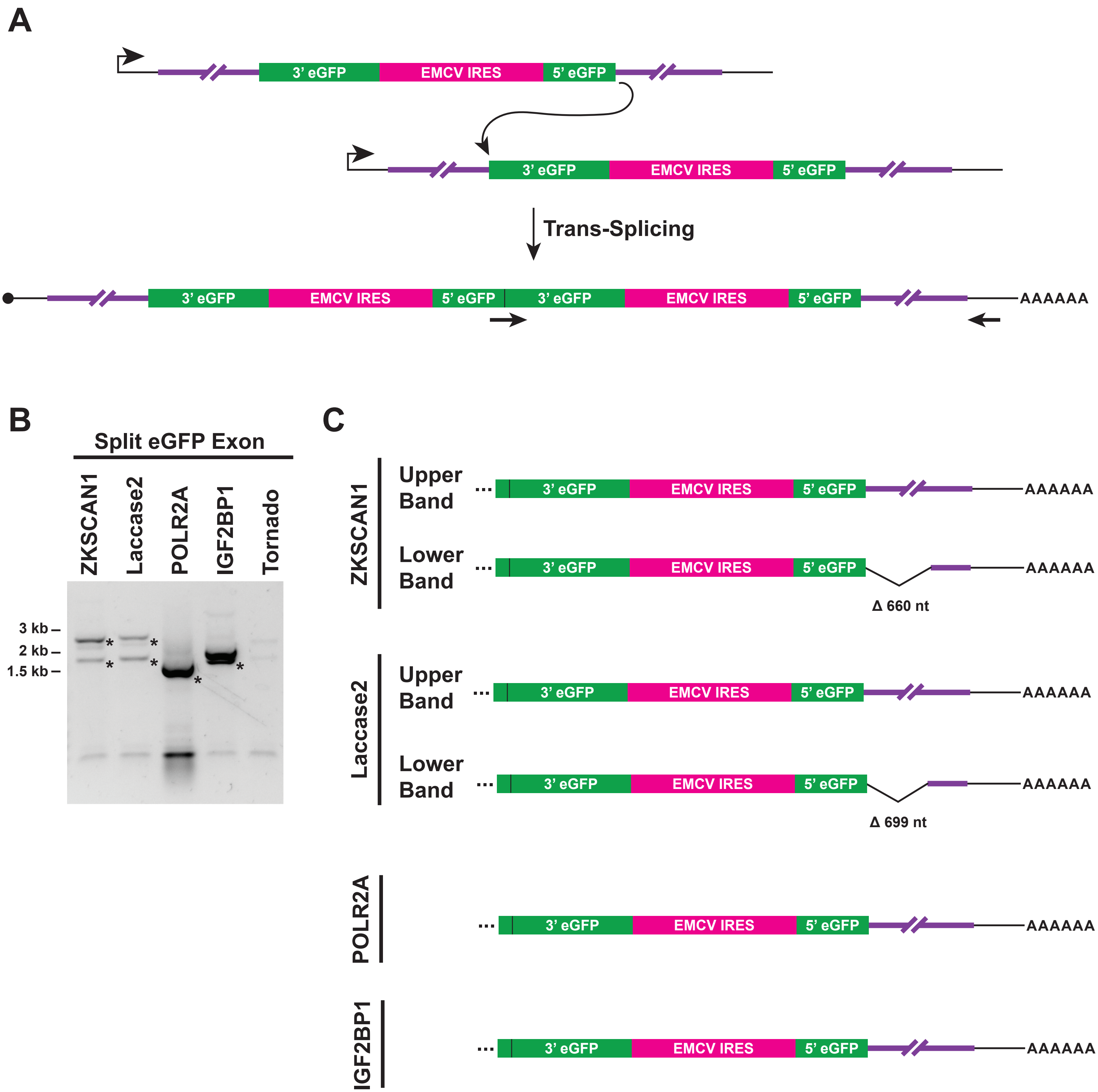

A
3' eGFP
EMCV IRES
5' eGFP
Trans-Splicing
AAAAAA
B
Split eGFP Exon
ZKSCAN1
Laccase2
POLR2A
IGF2BP1
Tornado
3 kb
2 kb
1.5 kb
C
ZKSCAN1
Upper Band
Lower Band
Δ 660 nt
Laccase2
Upper Band
Lower Band
Δ 699 nt
POLR2A
IGF2BP1

**Supplementary Figure S5. Trans-spliced RNAs can be generated from circRNA overexpression plasmids.**

**(A)** Trans-splicing to join the 5' splice site from one split eGFP/EMCV IRES containing transcript to the 3' splice site of another could yield a long linear mRNA with an uninterrupted eGFP that could be translated. PCR primers used to detect putative trans-spliced RNAs are indicated in black. The forward primer spans the backsplice/trans-splice junction (and thus should not be able to bind unspliced pre-mRNAs), while the reverse primer is complementary to the 3' region of linear transcripts. Only transcripts consistent with trans-splicing should be amplified with these primers.

**(B)** To test for the presence of such trans-spliced RNAs, the indicated expression plasmids were transfected into HEK293FT cells. Total RNA was isolated, reverse transcribed using random hexamers, and PCR was performed using the primers from above. * indicates bands that were gel purified.

**(C)** PCR products were individually gel purified and their consensus sequences identified using Nanopore sequencing. All detected products are consistent with trans-splicing, with the smaller transcripts generated by alternative splicing events in the 3' region. Full consensus sequences of all amplicons are provided in **Supplementary Table S3**. Note that the downstream intron sequences (purple) differ in length across constructs, thereby explaining the observed size differences in PCR amplicons, but are drawn as identical lengths here to facilitate visualization.

### Figure S6

**A**

| | 3' Splice Site | | 5' Splice Site | |
|---|---|---|---|---|
| ciRS-7-eGFP | 5.41 | ACAATGTTTCCAATGTCCAGGAG | 6.49 | CAGGTATTC |
| ZKSCAN1-eGFP | 8.92 | ACTTTTTTTTTATACTTCAGGAG | 10.77 | CAGGTAAGA |
| Laccase2-eGFP | 8.10 | ATTTTTTTTATTTTATGCAGGAG | 10.86 | CAGGTAAGT |

**B**

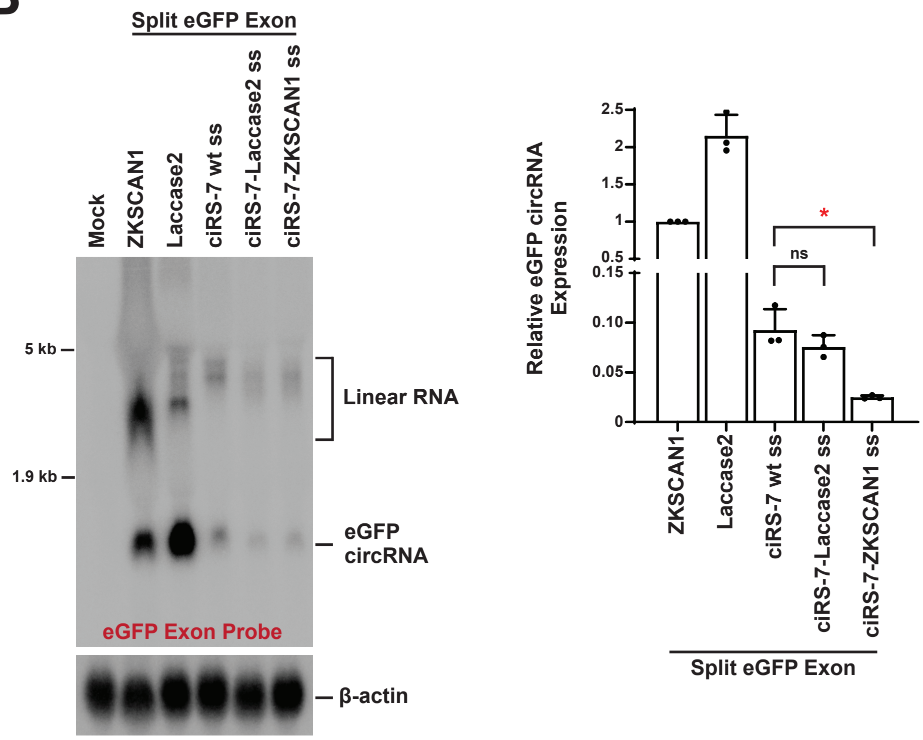

**Supplementary Figure S6. Improving splice site strength does not increase circRNA output from the *ciRS-7* introns.**

**(A)** Splice site strengths were evaluated using MaxEntScan [50] for constructs in which the split eGFP/EMCV IRES exon was inserted between the *ciRS-7*, *ZKSCAN1*, and *Laccase2* introns. Exon sequences are shown in orange.

**(B)** The split eGFP/EMCV IRES exon was cloned between the *ZKSCAN1*, *Laccase2*, or *ciRS-7* introns, as well as into modified *ciRS-7* introns in which the splice sites (ss) were replaced with those from *Laccase2* or *ZKSCAN1* (sequences shown in **A**). Plasmids were transfected into HEK293FT cells and analyzed by Northern blotting (left) and RT-qPCR (right) to quantify circGFP expression. Data are presented as mean ± SD (N = 3). p values were generated with unpaired t-tests, comparing each construct to the plasmid with wildtype *ciRS-7* splice sites. $^*p < 0.05$; ns, not significant.

### Figure S7

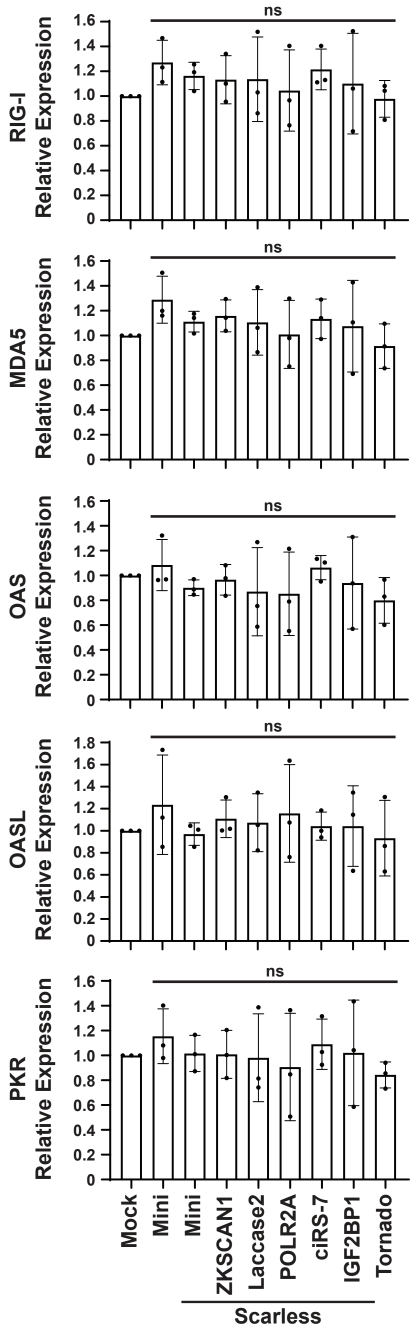

RIG-I Relative Expression
MDA5 Relative Expression
OAS Relative Expression
OASL Relative Expression
PKR Relative Expression
ns
0
0.2
0.4
0.6
0.8
1.0
1.2
1.4
1.6
1.8
Mock
Mini
Mini
ZKSCAN1
Laccase2
POLR2A
ciRS-7
IGF2BP1
Tornado
Scarless

**Supplementary Figure S7. None of the plasmids induce significant immune responses in HEK293FT cells.**

Plasmids expressing the split eGFP/EMCV IRES exon between the indicated intronic sequences were transfected into HEK293FT cells, and total RNA was isolated after 24 h. RT-qPCR was used to quantify expression of well-established immune response genes. Data are normalized to mock transfected cells and are presented as mean ± SD (N = 3). p values were generated with unpaired t-tests, comparing each construct to mock transfected cells. ns, not significant.

### Figure S8

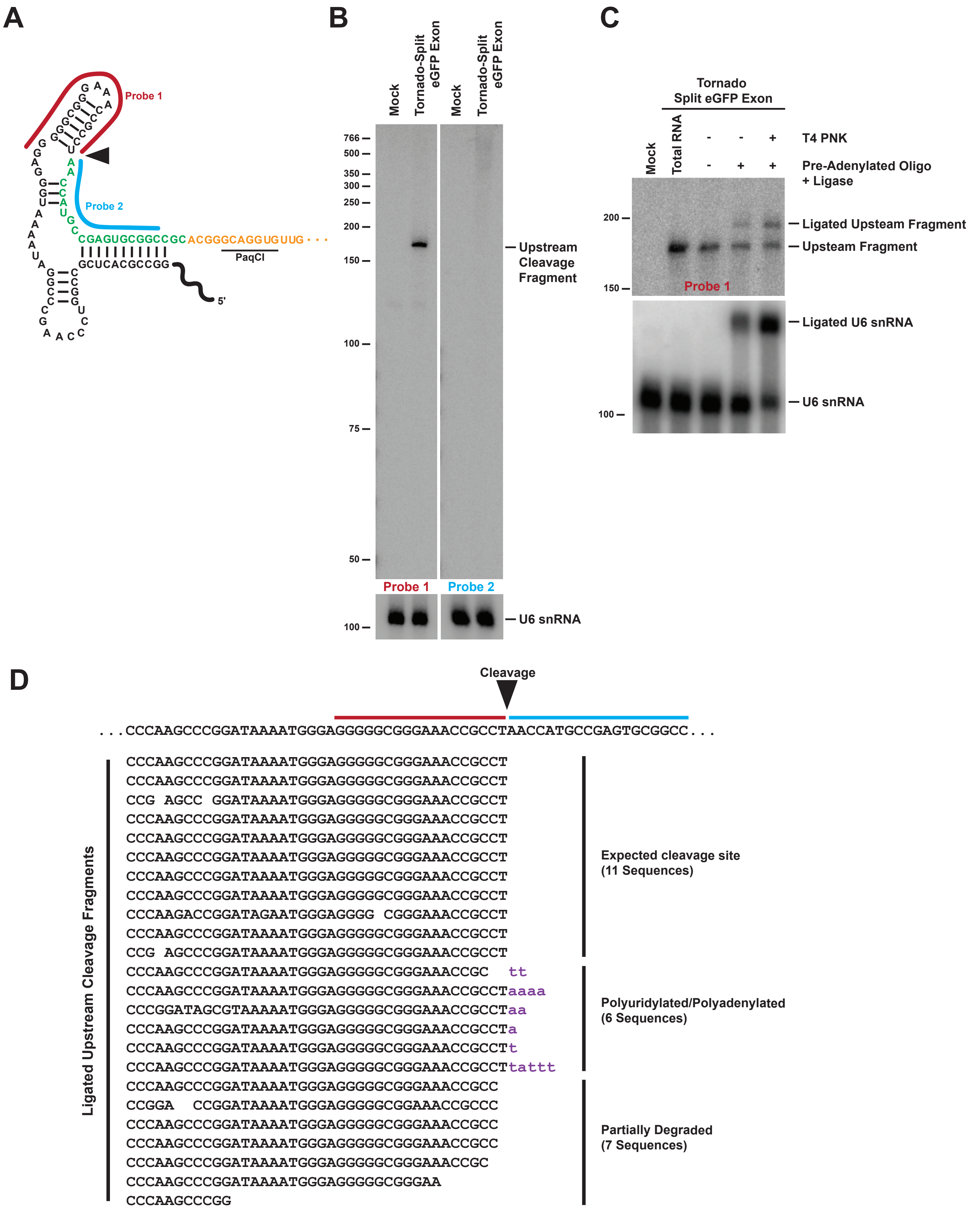

A
Probe 1
Probe 2
CGAGUGCGGCCGC
ACGGGCAGGUGUUG
GCUCACGCCGG
PaqCI
5'
B
Mock
Tornado-Split eGFP Exon
Mock
Tornado-Split eGFP Exon
766
500
350
300
250
200
150
100
75
50
Upstream Cleavage Fragment
Probe 1
Probe 2
100
U6 snRNA
C
Tornado Split eGFP Exon
Mock
Total RNA
T4 PNK
Pre-Adenylated Oligo + Ligase
200
150
100
Ligated Upsteam Fragment
Upsteam Fragment
Probe 1
Ligated U6 snRNA
U6 snRNA
D
Cleavage
...CCCAAGCCCGGATAAAATGGGAGGGGGCGGGAAACCGCCTAACCATGCCGAGTGCGGCC...
Ligated Upstream Cleavage Fragments
CCCAAGCCCGGATAAAATGGGAGGGGGCGGGAAACCGCCT
CCCAAGCCCGGATAAAATGGGAGGGGGCGGGAAACCGCCT
CCG AGCC GGATAAAATGGGAGGGGGCGGGAAACCGCCT
CCCAAGCCCGGATAAAATGGGAGGGGGCGGGAAACCGCCT
CCCAAGCCCGGATAAAATGGGAGGGGGCGGGAAACCGCCT
CCCAAGCCCGGATAAAATGGGAGGGGGCGGGAAACCGCCT
CCCAAGCCCGGATAAAATGGGAGGGGGCGGGAAACCGCCT
CCCAAGCCCGGATAAAATGGGAGGGGGCGGGAAACCGCCT
CCCAAGACCGGATAGAATGGGAGGGG CGGGAAACCGCCT
CCCAAGCCCGGATAAAATGGGAGGGGGCGGGAAACCGCCT
CCG AGCCCGGATAAAATGGGAGGGGGCGGGAAACCGCCT
Expected cleavage site (11 Sequences)
CCCAAGCCCGGATAAAATGGGAGGGGGCGGGAAACCGC tt
CCCAAGCCCGGATAAAATGGGAGGGGGCGGGAAACCGCCTaaaa
CCCGGATAGCGTAAAAATGGGAGGGGGCGGGAAACCGCCTaa
CCCAAGCCCGGATAAAATGGGAGGGGGCGGGAAACCGCCTa
CCCAAGCCCGGATAAAATGGGAGGGGGCGGGAAACCGCCTt
CCCAAGCCCGGATAAAATGGGAGGGGGCGGGAAACCGCCTtattt
Polyuridylated/Polyadenylated (6 Sequences)
CCCAAGCCCGGATAAAATGGGAGGGGGCGGGAAACCGCC
CCGGA CCGGATAAAATGGGAGGGGGCGGAAACCGCCC
CCCAAGCCCGGATAAAATGGGAGGGGGCGGGAAACCGCC
CCCAAGCCCGGATAAAATGGGAGGGGGCGGGAAACCGCC
CCCAAGCCCGGATAAAATGGGAGGGGGCGGGAAACCGC
CCCAAGCCCGGATAAAATGGGAGGGGGCGGGAA
CCCAAGCCCGG
Partially Degraded (7 Sequences)

**Supplementary Figure S8. The Tornado upstream cleavage fragment accumulates in cells.**

**(A)** Schematic of the 5' portion of the Tornado system, highlighting the ribozyme self-cleavage site (arrowhead) and the oligonucleotide probes used for Northern blotting.

**(B)** A plasmid expressing the split eGFP/EMCV IRES exon using the Tornado system was transfected into HEK293FT cells, and total RNA was isolated after 24 h. 18 μg of total RNA was analyzed by Northern blot using the indicated probes. U6 snRNA was used as a loading control.

**(C)** 10 μg of total RNA from HEK293FT cells transfected with the Tornado-driven split eGFP/EMCV IRES construct was treated with T4 polynucleotide kinase (T4 PNK) and then ligated to a pre-adenylated oligonucleotide, as indicated. "-" denotes control reactions containing buffer only. 5 μg of treated RNAs were then analyzed by Northern blot using probes complementary to the Tornado upstream cleavage fragment (Probe 1) or U6 snRNA, which is known to terminate in a 2',3'-cyclic phosphate [70].

**(D)** After employing T4 PNK and the ligation-based approach described in **C**, the 3' end of the Tornado upstream cleavage fragment was PCR amplified using a primer complementary to the ligated oligonucleotide. PCR products were then subjected to Nanopore sequencing. The adaptor sequence is not shown for clarity. We detected many transcripts terminating at the expected ribozyme cleavage site, including ones that have been uridylated or adenylated, likely as part of the degradation process.

### Figure S9

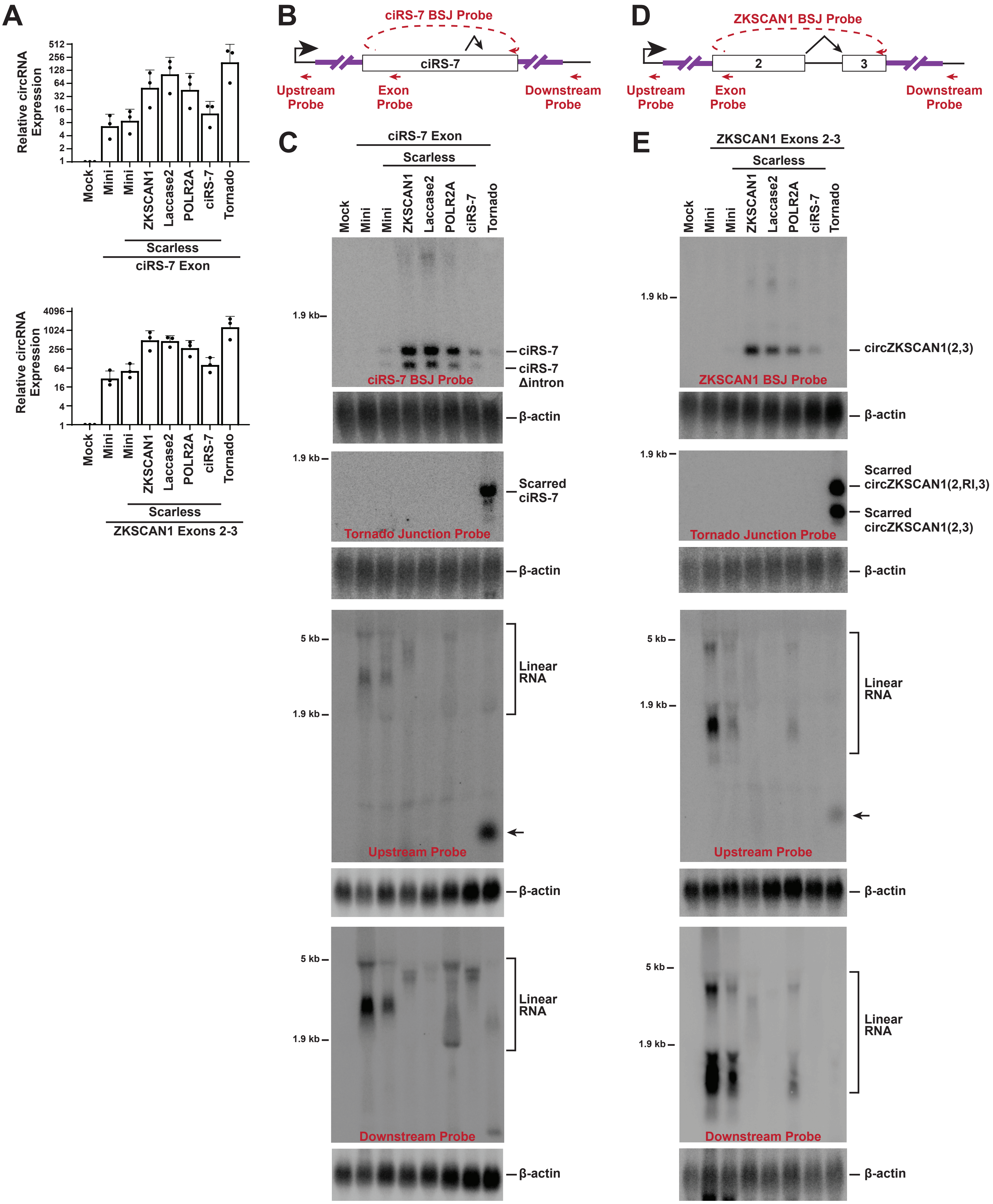

A
Relative circRNA Expression
512
256
128
64
32
16
8
4
2
1
Mock
Mini
Mini
ZKSCAN1
Laccase2
POLR2A
ciRS-7
Tornado
Scarless
ciRS-7 Exon
Relative circRNA Expression
4096
1024
256
64
16
4
1
Scarless
ZKSCAN1 Exons 2-3
B
ciRS-7 BSJ Probe
ciRS-7
Upstream Probe
Exon Probe
Downstream Probe
D
ZKSCAN1 BSJ Probe
2
3
C
ciRS-7 Exon
E
ZKSCAN1 Exons 2-3
1.9 kb
5 kb
ciRS-7
ciRS-7 Δintron
ciRS-7 BSJ Probe
β-actin
Scarred ciRS-7
Tornado Junction Probe
Linear RNA
Upstream Probe
Downstream Probe
circZKSCAN1(2,3)
ZKSCAN1 BSJ Probe
Scarred circZKSCAN1(2,RI,3)
Scarred circZKSCAN1(2,3)

**Supplementary Figure S9. Outputs of ciRS-7 and circZKSCAN1(2,3) expression plasmids.**

**(A)** Plasmids expressing the ciRS-7 exon (top) or exons 2 and 3 of ZKSCAN1 (bottom) flanked by the indicated intronic sequences were transfected into HEK293FT cells. Total RNA was isolated 24 h post-transfection, and circRNA levels were quantified by RT-qPCR. Data were normalized to mock transfected cells and are presented as mean ± SD (N = 3).

**(B)** Schematic of ciRS-7 expression vectors and oligonucleotide probes used for Northern blotting. The upstream and downstream probes target regions common to all vectors.

**(C)** Plasmids expressing the ciRS-7 exon flanked by the indicated intronic sequences were transfected into HEK293FT cells, and total RNA was isolated 24 h post-transfection. 18 μg of total RNA was analyzed by Northern blot using the indicated probes. β-actin was used as a loading control.

**(D)** Schematic of circZKSCAN1(2,3) expression vectors and oligonucleotide probes used for Northern blotting.

**(E)** Plasmids expressing ZKSCAN1 exons flanked by the indicated intronic sequences were transfected into HEK293FT cells, and total RNA was isolated 24 h post-transfection. 18 μg of total RNA was analyzed by Northern blot using the indicated probes. β-actin was used as a loading control. RI, retained intron.

### Figure S10

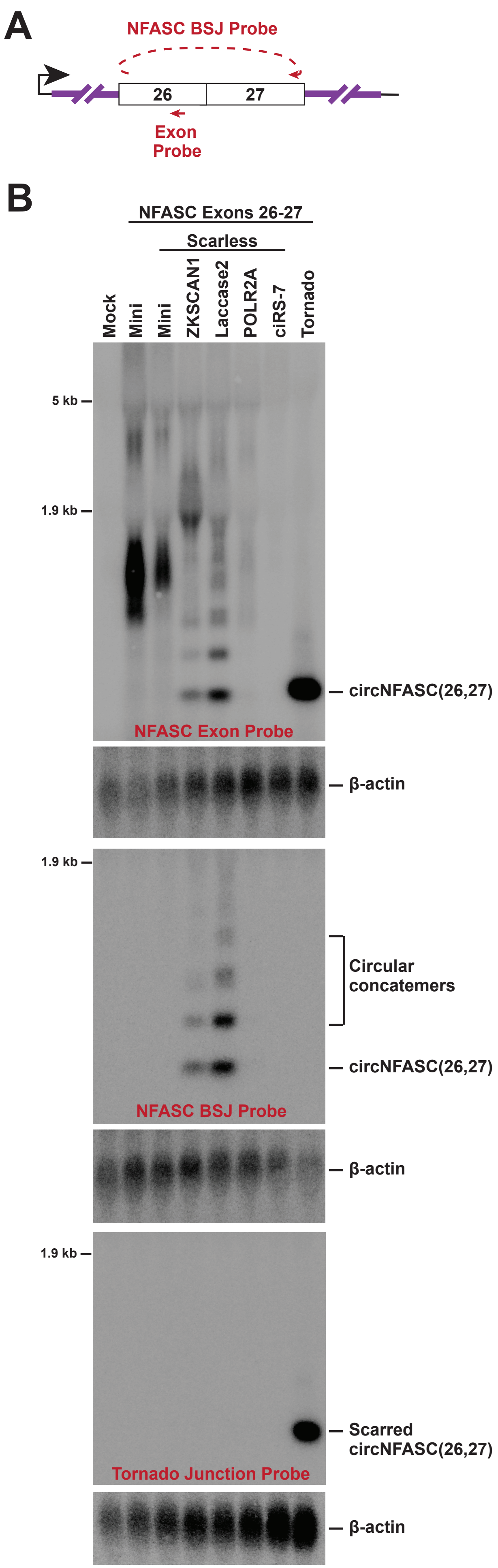

A
NFASC BSJ Probe
26
27
Exon
Probe
B
NFASC Exons 26-27
Scarless
Mock
Mini
Mini
ZKSCAN1
Laccase2
POLR2A
ciRS-7
Tornado
5 kb
1.9 kb
circNFASC(26,27)
NFASC Exon Probe
β-actin
1.9 kb
Circular
concatemers
circNFASC(26,27)
NFASC BSJ Probe
β-actin
1.9 kb
Scarred
circNFASC(26,27)
Tornado Junction Probe
β-actin

**Supplementary Figure S10. The Tornado system can generate a small circRNA more effectively than spliceosome-dependent strategies.**

**(A)** Backsplicing of exons 26 and 27 of human NFASC generates circNFASC(26,27), a 270-nt circRNA. The oligonucleotide probes used for Northern blotting are indicated in red.

**(B)** Plasmids expressing NFASC exons flanked by the indicated intronic sequences were transfected into HEK293FT cells, and total RNA was isolated 24 h post-transfection. 18 μg of total RNA was analyzed by Northern blot using the indicated probes. β-actin was used as a loading control.

### Figure S11

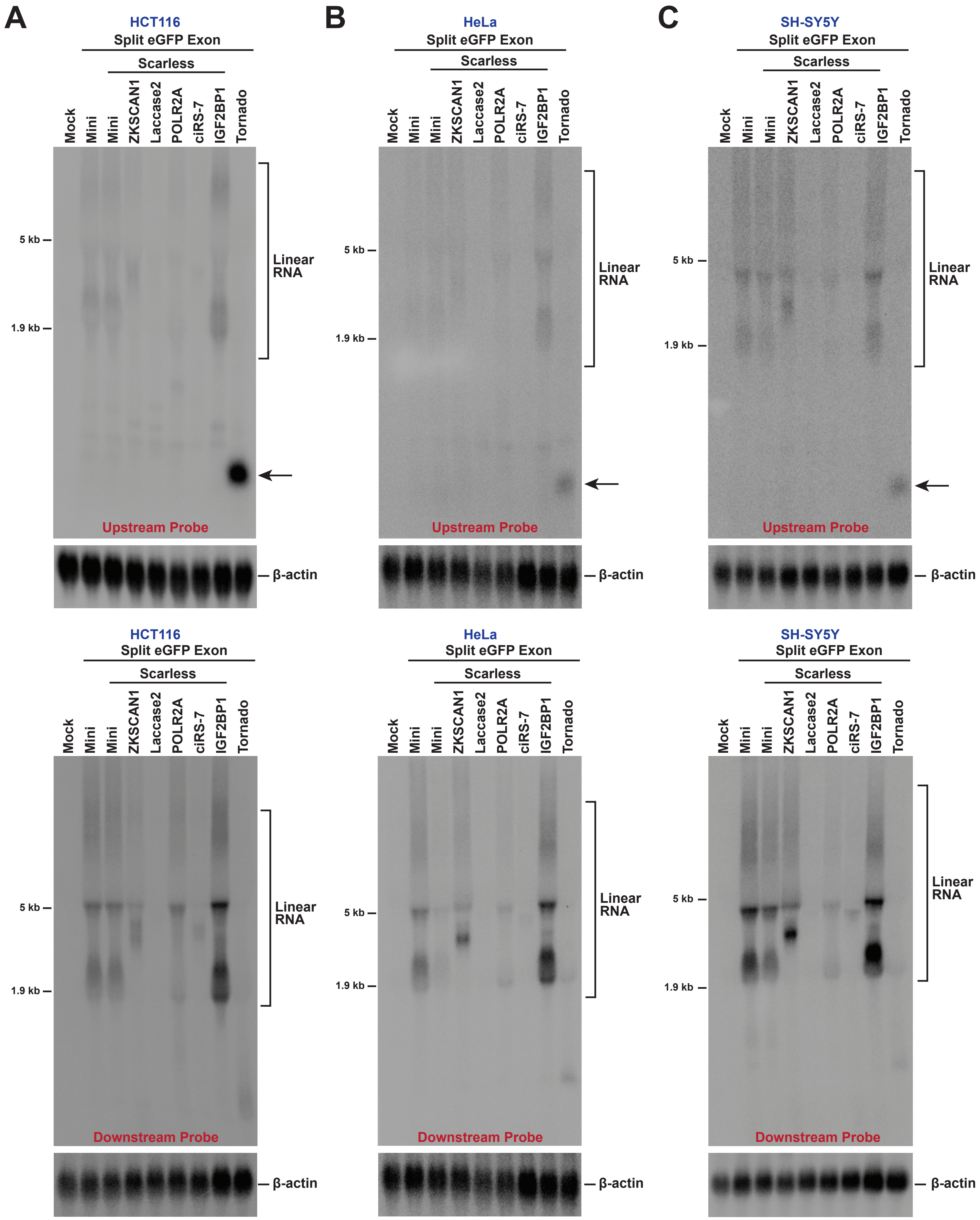

A
HCT116
Split eGFP Exon
Scarless
Mock
Mini
Mini
ZKSCAN1
Laccase2
POLR2A
ciRS-7
IGF2BP1
Tornado
5 kb
1.9 kb
Linear RNA
Upstream Probe
β-actin
B
HeLa
C
SH-SY5Y
Downstream Probe

**Supplementary Figure S11. Linear byproducts of circRNA overexpression strategies in HCT116, HeLa, and SH-SY5Y cells.**

**(A-C)** Plasmids expressing the split eGFP/EMCV IRES exon between the indicated intronic sequences were transfected into **(A)** HCT116, **(B)** HeLa, and **(C)** SH-SY5Y cells, and total RNA was isolated 24 h post-transfection. 18 $\mu$g of total RNA was analyzed by Northern blot using the indicated probes (locations diagrammed in **Figure 2A**). $\beta$-actin was used as a loading control.

### Figure S12

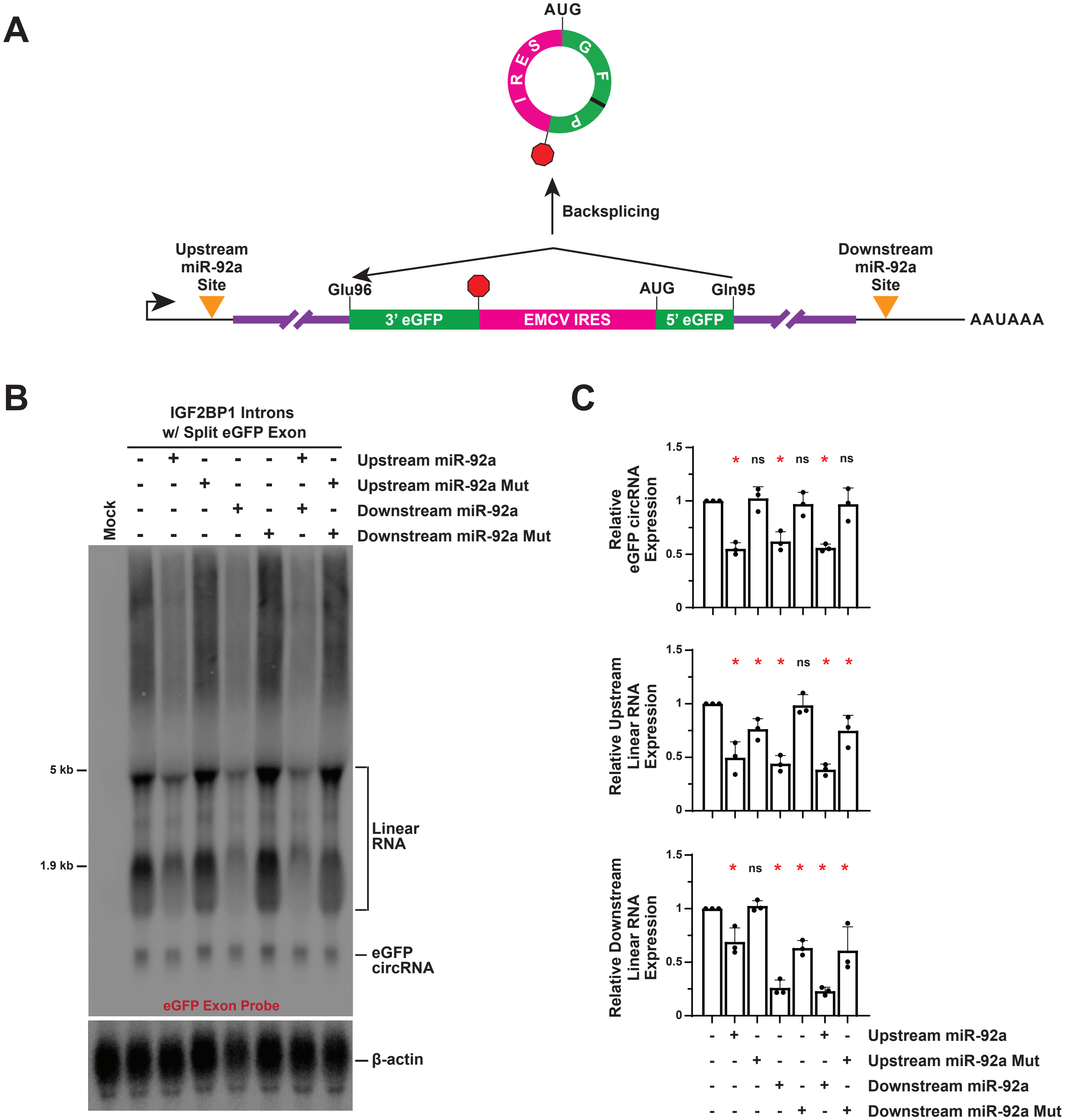
A
AUG
IRES
GFP
Backsplicing
Upstream miR-92a Site
Glu96
AUG
Gln95
Downstream miR-92a Site
3' eGFP
EMCV IRES
5' eGFP
AAUAAA
B
IGF2BP1 Introns w/ Split eGFP Exon
Mock
Upstream miR-92a
Upstream miR-92a Mut
Downstream miR-92a
Downstream miR-92a Mut
5 kb
1.9 kb
Linear RNA
eGFP circRNA
eGFP Exon Probe
β-actin
C
Relative eGFP circRNA Expression
Relative Upstream Linear RNA Expression
Relative Downstream Linear RNA Expression
ns
Upstream miR-92a
Upstream miR-92a Mut
Downstream miR-92a
Downstream miR-92a Mut

**Supplementary Figure S12. Introduction of microRNA target sites into flanking introns reduces bystander linear RNAs.**

**(A)** Perfectly complementary target sites for miR-92a were inserted near the 5' end, 3' end, or both ends of constructs containing the eGFP/EMCV IRES exon expression cassette to selectively trigger degradation of bystander linear RNAs, while sparing the mature circRNA.

**(B-C)** Plasmids expressing the split eGFP/EMCV IRES exon flanked by *IGF2BP1* introns (containing either wildtype or mutant [Mut] miR-92a target sites, as indicated) were transfected into HEK293FT cells, and total RNA was isolated 24 h post-transfection. **(B)** 18 μg of total RNA was analyzed by Northern blot using a probe complementary to the eGFP ORF. β-actin was used as a loading control. **(C)** RT-qPCR was used to quantify circRNA and linear RNA expression from each plasmid. Data are normalized to the plasmid lacking miR-92a target sites and are presented as mean ± SD (N = 3). p values were generated with unpaired t-tests, comparing each construct to the construct lacking miR-92a target sites. $*p < 0.05$; ns, not significant.

### Figure S13

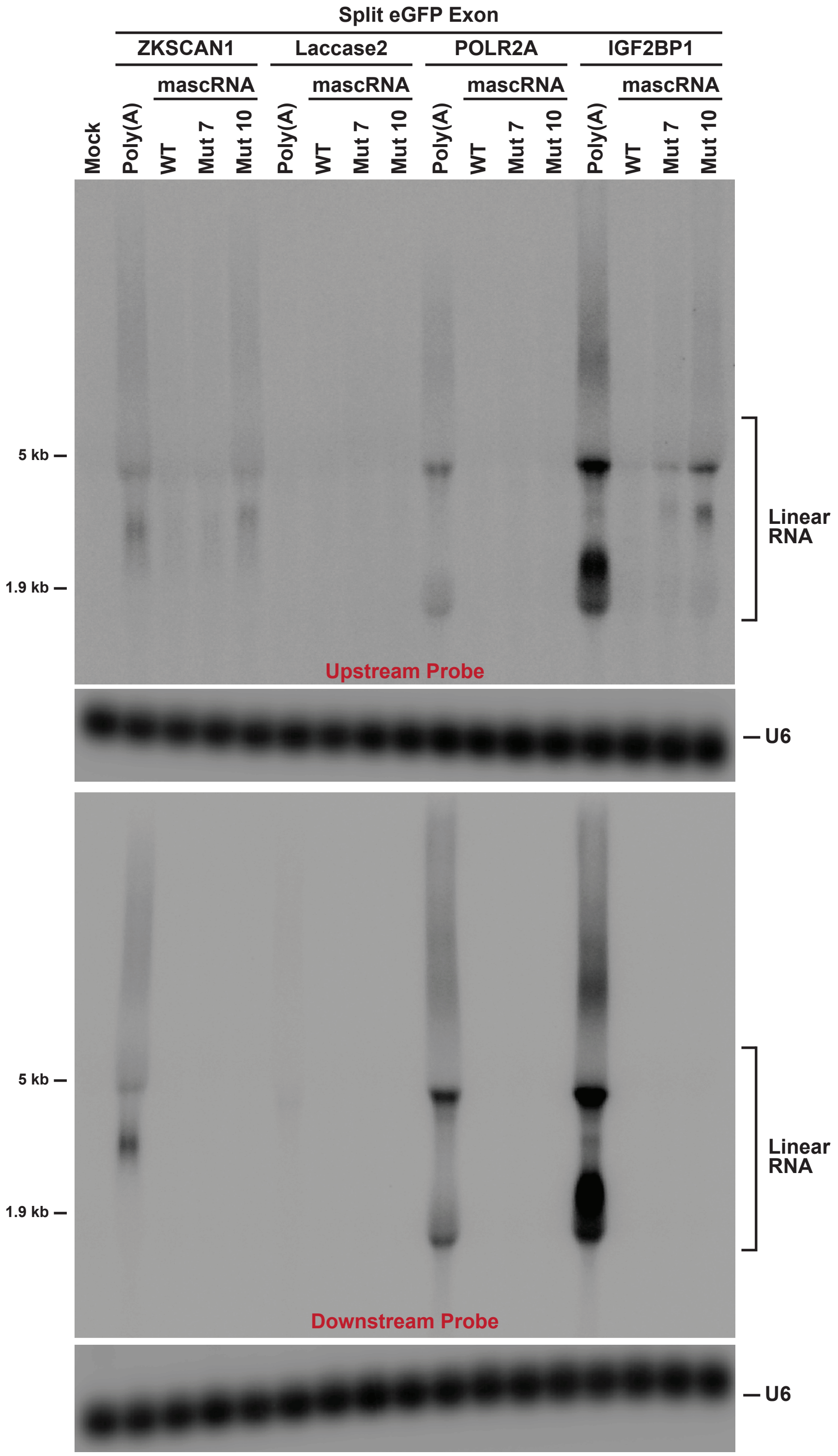

Split eGFP Exon
ZKSCAN1
Laccase2
POLR2A
IGF2BP1
mascRNA
mascRNA
mascRNA
mascRNA
Mock
Poly(A)
WT
Mut 7
Mut 10
Poly(A)
WT
Mut 7
Mut 10
Poly(A)
WT
Mut 7
Mut 10
Poly(A)
WT
Mut 7
Mut 10
5 kb
1.9 kb
Linear RNA
Upstream Probe
U6
5 kb
1.9 kb
Linear RNA
Downstream Probe
U6

**Supplementary Figure S13. Destabilization of the pre-mRNA 3' end reduces bystander linear RNAs.**

As in **Figure 5**, plasmids expressing the split eGFP/EMCV IRES exon between the indicated intronic sequences and terminating in a poly(A) signal or a mascRNA variant were transfected into HEK293FT cells, and total RNA was isolated after 24 h. 18 μg of total RNA was analyzed by Northern blot using probes complementary to the upstream or downstream common regions of the linear RNAs (probe locations indicated in **Figure 2A**). U6 snRNA was used as a loading control.

### Figure S14

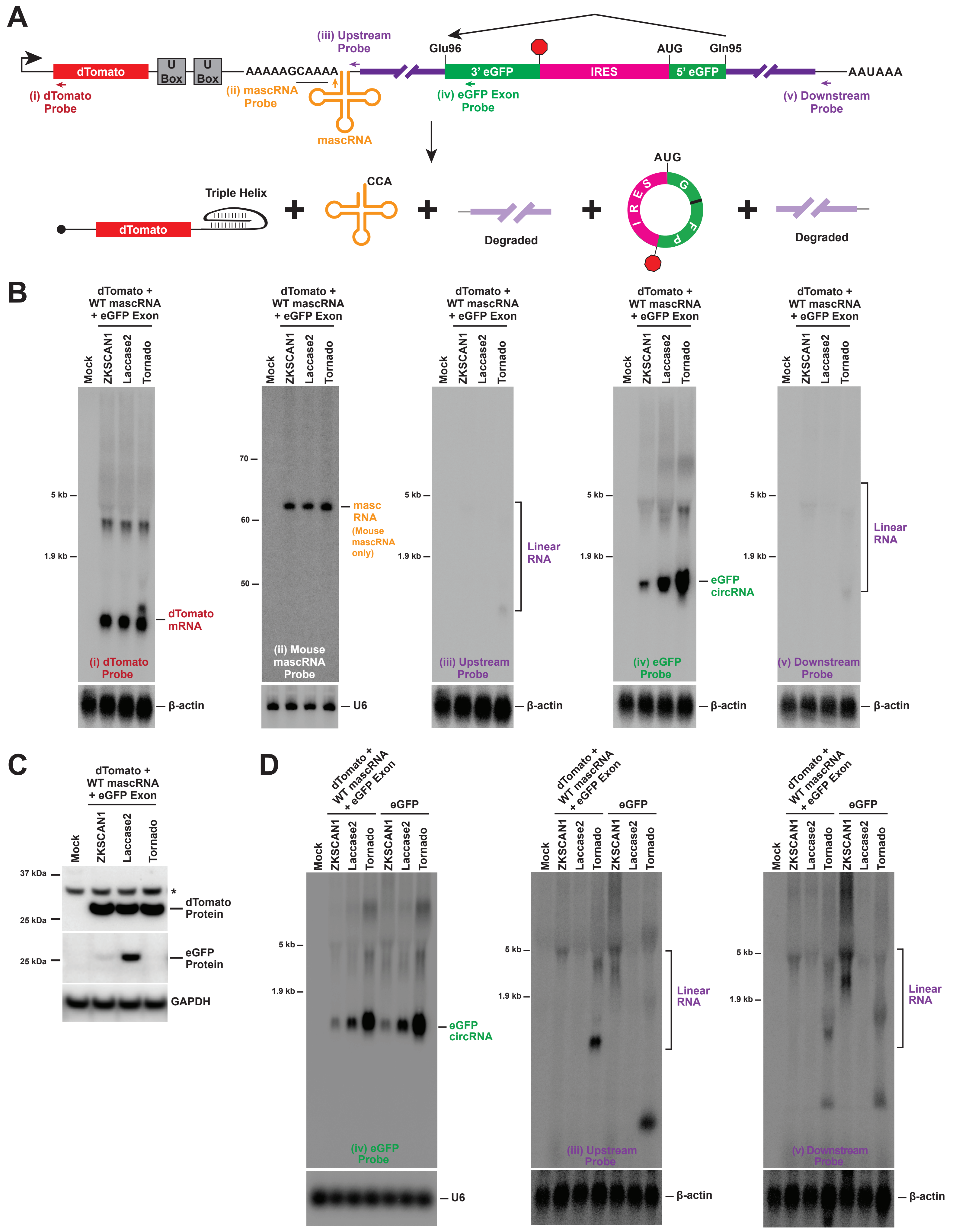

A
dTomato
U Box
U Box
AAAAAGCAAAA
(iii) Upstream Probe
Glu96
3' eGFP
IRES
AUG
Gln95
5' eGFP
AAUAAA
(i) dTomato Probe
(ii) mascRNA Probe
mascRNA
(iv) eGFP Exon Probe
(v) Downstream Probe
Triple Helix
CCA
Degraded
AUG
IRES
GFP
Degraded
B
dTomato + WT mascRNA + eGFP Exon
Mock
ZKSCAN1
Laccase2
Tornado
5 kb
1.9 kb
dTomato mRNA
(i) dTomato Probe
β-actin
70
60
50
masc RNA
(Mouse mascRNA only)
(ii) Mouse mascRNA Probe
U6
Linear RNA
(iii) Upstream Probe
eGFP circRNA
(iv) eGFP Probe
(v) Downstream Probe
C
37 kDa
25 kDa
dTomato Protein
eGFP Protein
GAPDH
*
D
eGFP
(iv) eGFP Probe
(iii) Upstream Probe
(v) Downstream Probe

**Supplementary Figure S14. CIRCUS platform built with wildtype mascRNA confirms proper processing of nascent transcripts.**

**(A)** Schematic of the CIRCUS platform enabling co-expression of a linear mRNA encoding dTomato with a circRNA encoding eGFP. This construct is processed analogously to **Figure 6**, except that wildtype mascRNA is employed and thus accumulates in cells. Oligonucleotide probes used for Northern blotting are indicated.

**(B)** The CIRCUS plasmids with wildtype mascRNA and the indicated sequences flanking the split eGFP/EMCV IRES exon were transfected into HEK293FT cells, and total RNA was isolated after 24 h. 18 $\mu$g of total RNA was analyzed by Northern blots with the indicated probes. $\beta$-actin and U6 snRNA were used as loading controls.

**(C)** Total protein was likewise isolated 24 h post-transfection. 20 $\mu$g of total protein was analyzed by immunoblotting using an anti-eGFP antibody. GAPDH was used as a loading control. *, non-specific band.

**(D)** The output of CIRCUS plasmids expressing linear dTomato and an eGFP circRNA **(Supplementary Figure S14A)** were benchmarked against plasmids that only express the eGFP circRNA **(Figure 2A)**. Plasmids with the indicated sequences flanking the split eGFP/EMCV IRES exon were transfected into HEK293FT cells, and total RNA was isolated after 24 h. 18 $\mu$g of total RNA was analyzed by Northern blots with the indicated probes. $\beta$-actin or U6 snRNA was used as a loading control. Addition of the dTomato ORF did not markedly affect eGFP circRNA yield, and the *Laccase2* introns resulted in the least amount of contaminating linear RNAs.

### Figure S15

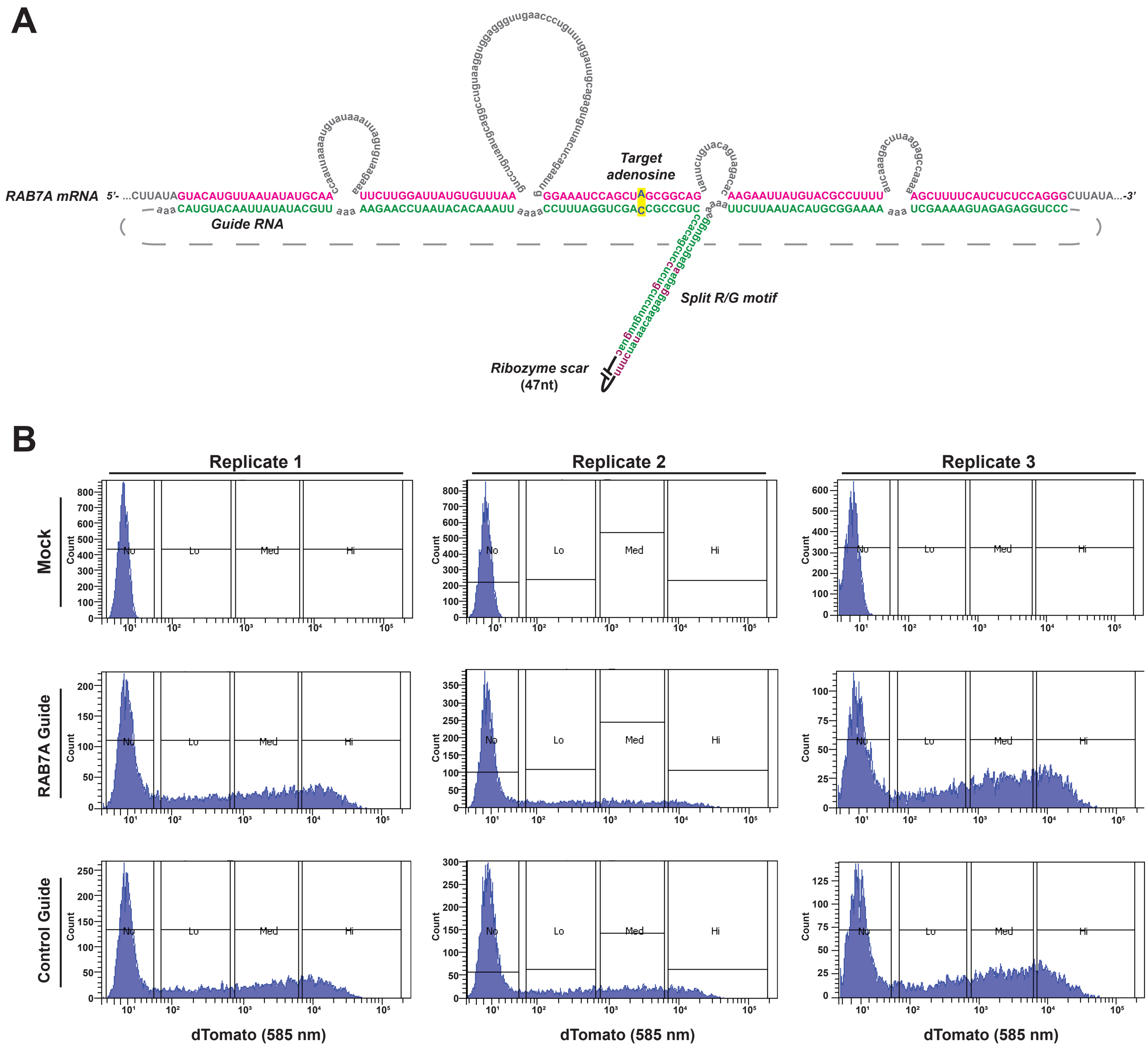

A
RAB7A mRNA
5'- ...CUUAUAGUACAUGUUAAUAUAUGCAA UUCUUGGAUUAUGUGUUUAA GGAAAUCCAGCUAGCGGCAG AAGAAUUAUGUACGCCUUUU AGCUUUUCAUCUCUCCAGGGCUUAUA...-3'
aaa CAUGUACAAUUAUAUACGUU aaa AAGAACCUAAUACACAAAUU aaaa CCUUUAGGUCGACCGCCGUC aaa UUCUUAAUACAUGCGGAAAA aaa UCGAAAAGUAGAGAGGUCCC
Guide RNA
Target
adenosine
Split R/G motif
Ribozyme scar
(47nt)
B
Replicate 1
Replicate 2
Replicate 3
Mock
RAB7A Guide
Control Guide
Count
No
Lo
Med
Hi
dTomato (585 nm)

**Supplementary Figure S15. RAB7A guide RNA design and FACS gating strategy.**

**(A)** Schematic depicting regions of complementarity between the RAB7A 3' UTR (pink) and the circular RAB7A guide RNA (green). The target adenosine is highlighted in yellow, with an A:C mismatch at this site to enhance editing efficiency [67]. The split-R/G motif in the guide RNA is modified from the GluR2 glutamate receptor mRNA and helps recruit ADAR [28]. For clarity, the 47-nt Tornado junction scar sequence is not shown, and the circRNA is depicted in a linear conformation. The dashed line indicates the two nucleotides that are adjacent to each other in the circRNA.

**(B)** FACS plots illustrating the gating strategy used to sort dTomato-expressing cells in each biological replicate. Cells were first gated on forward scatter (FSC) and side scatter (SSC) to exclude debris, followed by doublet discrimination using FSC-height versus FSC-area. dTomato fluorescence intensity was plotted on a log10 scale, and gates were set based on a mock-transfected control to define the dTomato-negative population. The remaining dTomato-positive cells were divided into three non-overlapping sequential gates (Low, Medium, and High) spanning the observed range of fluorescence intensities.

#### SUPPLEMENTARY TABLES

**Supplementary Table S1. Northern blot probes.**

**Supplementary Table S2. RT-qPCR Primers.**

**Supplementary Table S3. Nanopore sequencing results from RT-PCR to detect trans-spliced eGFP RNAs.**

**Supplementary Table S4. Nanopore sequencing results for detecting A-to-I editing of the RAB7A 3' UTR.**

### Supplementary Table S1

**Northern blot probes**

| **Probe** | **Probe Sequence** | **Figure Panels** | **Supplementary Figure Panels** |
|---|---|---|---|
| β-Actin | AGCACTGTGTTGGCGTACAG | 2B, 2D, 2F, 2G, 3B, 3E, 4A, 4D, 4G, 6B | S3, S4B, S6B, S9C, S9E, S10, S11A, S11B, S11C, S12B, S14B, S14D |
| eGFP Exon Probe | TTGCCGTCCTCCTTGAAGTC | 2B, 4A, 4D, 4G, 5B, 6B | S3, S4B, S6B, S12B, S14B, S14D |
| eGFP BSJ Probe | GTGCGCTCCTGGACGT | 2D | |
| Tornado Junction Probe | GGCATGGTTCTACAGTCC | 2D | S9C, S9E, S10 |
| Upstream Probe | GAATTCCACCACACTGGAC | 2F, 6B | S3, S9C, S9E, S11A, S11B, S11C, S13, S14B, S14D |
| Downstream Probe | GGGAGTGGCACCTTCCAG | 2G. 6B | S3, S9C, S9E, S11A, S11B, S11C, S13, S14B, S14D |
| Probe 1 (Tornado Cleavage Product) | AGGCGGTTTCCCGCCCCC | | S8B, S8C |
| Probe 2 (Tornado Cleavage Product) | GCCGCACTCGGCATGGTT | | S8B |
| ciRS-7 Exon Probe | TGGAAGATCTGTATTTGATGGAAGACCTT | 3B | |
| ciRS-7 BSJ Probe | TCGGAAACCCTGGATATT | | S9C |
| ZKSCAN1 Exon Probe | AGCAGTCATCATTCAGGCTCCA | 3E | |
| ZKSCAN1 BSJ Probe | TTTACTATTCCTCGTGAC | | S9E |
| NFASC Exon Probe | GGGCCCGAAATTGTGCTTCC | | S10 |
| NFASC BSJ Probe | ATCAGGGGCTGTACTG | | S10 |
| mascRNA (Mouse mascRNA Only Probe) | AGGAGTGCCAGCCACCAGCG | 5B | S14B |
| mascRNA (All mascRNA Probe) | GCAAAGACACCGCAGGGATTTGAACCCCGTCCTGGAAACCAGGAGTGCCA | 5B, 6B | |
| U6 | GCTAATCTTCTCTGTATCGTTCCAATTTTAGTATATGTGCTGCCG | 5B, 6B | S8B, S8C, S13, S14B, S14D |
| dTomato Probe | TTCTTGTAATCGGGGATGTCGG | 6B | S14B |

### Supplementary Table S2

**RT-qPCR Primers**

| Target RNA | Forward Primer | Reverse Primer | Figure Panels | Supplementary Figure Panels |
|---|---|---|---|---|
| GAPDH | GGTGGTCTCCTCTGACTTCAACA | GTTGCTGTAGCCAAATTCGTTGT | 2C, 3C, 3F, 4B, 4E, 4H, 5C, 7C | S6B, S7, S9A, S12C |
| eGFP circRNA | GCGTGCAGTGCTTCAGCCGC | CCTTGAAGAAGATGGTGCGCTC | 2C, 4B, 4E, 4H, 5C | S6B, S12C |
| Upstream Linear RNA | GCTTACTGGCTTATCGAAAT | ATCCGAGCTCGGTACCAAGC | 2C, 3C, 3F, 4B, 4E, 4H, 5C | S12C |
| Downstream Linear RNA | AGGGCCCGTTTAAACCCG | GGGAGTGGCACCTTCCAG | 2C, 3C, 3F, 4B, 4E, 4H, 5C | S12C |
| ciRS-7 | ACCCAGTCTTCCATCAACTGGC | TCTTGGAAGACATGGATTGTCCGG | 3C | S9A |
| circZKSCAN1(2,3) | CCAGTTCAGGAGTCCTCGAGC | CTTGATCCAAACAGGGTCTGTGC | 3F | S9A |
| RIG-I | TTCATGTCCACCTTCAGAAGTG | TCATAGCAGGCAAAGCAAGC | | S7 |
| MDA5 | GGCACCATGGGAAGTGATT | ATTTGGTAAGGCCTGAGCTG | | S7 |
| OAS | GCTCCTACCCTGTGTGTGTGT | TGGTGAGAGGAGTGAGGAAGA | | S7 |
| OASL | AGGGTACAGATGGGACATCG | AAGGGTTCACGATGAGGTTG | | S7 |
| PKR | TTCTTCCCCGGCTGCTT | CTCGCTCTCTTGCTTCCTCTCT | | S7 |
| dTomato | CAAGCTGAAGGTGACCAAGGG | TTCTTGTAATCGGGGATGTCGG | 7C | |
| RAB7A guide RNA | CACCCTGCCGCCAGCTGG | GCTTTTCATCTCTCCAGGGTTTG | 7C | |

### Supplementary Table S3

**Nanopore sequencing results from RT-PCR to detect trans-spliced eGFP RNAs**

| Sample | Consensus Sequence |
|---|---|
| ZKSCAN1 Upper Band | CTACGTCCAGGAGCGCACCATCTTCTTCAAGGACGACGGCAACTACAAGACCCGCGCCGAGGTGAAGTTCGAGGGCGACACCCTGGTGAACCGCATCGAGCT<br>GAAGGGCATCGACTTCAAGGAGGACGGCAACATCCTGGGGCACAAGCTGGAGTACAACCACAACAGCCACAACGTCTATATCATGGCCGACAAGCAGAAGAAC<br>GGCATCAAGGTGAACTTCAAGATCCGCCACAACATCGAGGACGGCAGCGTGCAGCTCGCCGACCACTACCAGCAGAACACCCCCATCGGCGACGGCCCCGTG<br>CTGCTGCCCGACAACCACTACCTGAGCACCCAGTCCGCCCTGAGCAAAGACCCCAACGAGAAGCGCGATCACATGGTCCTGCTGGAGTTCGTGACCGCCGCC<br>GGGATCACTCTCGGCATGGACGAGCTGTACAAGTAAGGATCTGAATTCTGCAGTCGACGATCCGCCCCTCTCCCTCCCCCCCCCCTAACGTTACTGGCCGAAG<br>CCGCTTGGAATAAGGCCGGTGTGCGTTTGTCTATATGTTATTTTCCACCATATTGCCGTCTTTTGGCAATGTGAGGGCCCGGAAACCTGGCCCTGTCTTCTTGA<br>CGAGCATTCCTAGGGGTCTTTCCCCTCTCGCCAAAGGAATGCAAGGTCTGTTGAATGTCGTGAAGGAAGCAGTTCCTCTGGAAGCTTCTTGAAGACAAACAACG<br>TCTGTAGCGACCCTTTGCAGGCAGCGGAACCCCCCACCTGGCGACAGGTGCCTCTGCGGCCAAAAGCCACGTGTATAAGATACACCTGCAAAGGCGGCACAA<br>CCCCAGTGCCACGTTGTGAGTTGGATAGTTGTGGAAAGAGTCAAATGGCTCTCCTCAAGCGTATTCAACAAGGGGCTGAAGGATGCCCAGAAGGTACCCCATT<br>GTATGGGATCTGATCTGGGGCCTCGGTACACATGCTTTACATGTGTTTAGTCGAGGTTAAAAAAACGTCTAGGCCCCCCGAACCACGGGGACGTGGTTTTCCT<br>TTGAAAAACACGATGATAATATGGCCACAACCATGGTGAGCAAGGGCGAGGAGCTGTTCACCGGGGTGGTGCCCATCCTGGTCGAGCTGGACGGCGACGTAA<br>ACGGCCACAAGTTCAGCGTGTCCGGCGAGGGCGAGGGCGATGCCACCTACGGCAAGCTGACCCTGAAGTTCATCTGCACCACCGGCAAGCTGCCCGTGCCC<br>TGGCCCACCCTCGTGACCACCCTGACCTACGGCGTGCAGTGCTTCAGCCGCTACCCCGACCACATGAAGCAGCACGACTTCTTCAAGTCCGCCATGCCCGAA<br>GGCTACGTCCAGGTAAGAAGCAAGGTTTCATTTAGGGGAAGGGAAATGATTCAGGACGAGAGTCTTTGTGCTGCTGAGTGCCTGTGATGAAGAAGCATGTTAG<br>TCCTGGGCAACGTAGCGAGACCCCATCTCTACAAAAAATAGAAAAATTAGCCAGGTATAGTGGCGCACACCTGTGATTCCAGCTACGCAGGAGGCTGAGGTGG<br>GAGGATTGCTTGAGCCCAGGAGGTTGAGGCTGCAGTGAGCTGGAATCATGCCACTACTCCAACCTGGGCAACACAGCAAGGACCCTGTCTCAAAAGCTACTTA<br>CAGAAAAGAATTAGGCTCGGCACGGTAGCTCACACCTGTAATCCCAGCACTTTGGGAGGCTGAGGCGGGCAGATCACTTGAGGTCAGGAGTTTGAGACCAGC<br>CTGGCCAACATGGTGAAACCTTGTCTCTACTAAAAATATGAAAATTAGCCAGGCATGGTGGCACATTCCTGTAATCCCAGCTACTCGGGAGGCTGAGGCAGGAG<br>AATCACTTGAACCCAGGAGGTGGAGGTTGCAGTAAGCCGAGATCGTACCACTGTGCTCTAGCCTTGGTGACAGAGCGAGACTGTCTTAAAAAAAAAAAAAAAA<br>GAATTAATTAAAAATTTAAAAAAAAATGAAAAAAAGCTGCATGCTTGTTTTTTGTTTTTAGTTATTCTACATTGTTGTCATTATTACCAAATATTGGGGAAAATACAAC<br>TTACAGACCAATCTCAGGAGTTAAATGTTACTACGAAGGCAAATGAACTATGCGTAATGAACCTGGTAGGCATTAGCGGCCGCTCGAGTCTAGAGGGCCCGTTT<br>AAACCCGCTGATCAGCCTCGACTGTGCCTTCTAGTTGCCAGCCATCTGTTGTTTGCCCCTCCCCCGTGCCTTCCTTGACCCTGGAAGGTG |
| ZKSCAN1 Lower Band | CTACGTCCAGGAGCGCACCATCTTCTTCAAGGACGACGGCAACTACAAGACCCGCGCCGAGGTGAAGTTCGAGGGCGACACCCTGGTGAACCGCATCGAGCT<br>GAAGGGCATCGACTTCAAGGAGGACGGCAACATCCTGGGGCACAAGCTGGAGTACAACCACAACAGCCACAACGTCTATATCATGGCCGACAAGCAGAAGAAC<br>GGCATCAAGGTGAACTTCAAGATCCGCCACAACATCGAGGACGGCAGCGTGCAGCTCGCCGACCACTACCAGCAGAACACCCCCATCGGCGACGGCCCCGTG<br>CTGCTGCCCGACAACCACTACCTGAGCACCCAGTCCGCCCTGAGCAAAGACCCCAACGAGAAGCGCGATCACATGGTCCTGCTGGAGTTCGTGACCGCCGCC<br>GGGATCACTCTCGGCATGGACGAGCTGTACAAGTAAGGATCTGAATTCTGCAGTCGACGATCCGCCCCTCTCCCTCCCCCCCCCCTAACGTTACTGGCCGAAG<br>CCGCTTGGAATAAGGCCGGTGTGCGTTTGTCTATATGTTATTTTCCACCATATTGCCGTCTTTTGGCAATGTGAGGGCCCGGAAACCTGGCCCTGTCTTCTTGA<br>CGAGCATTCCTAGGGGTCTTTCCCCTCTCGCCAAAGGAATGCAAGGTCTGTTGAATGTCGTGAAGGAAGCAGTTCCTCTGGAAGCTTCTTGAAGACAAACAACG<br>TCTGTAGCGACCCTTTGCAGGCAGCGGAACCCCCCACCTGGCGACAGGTGCCTCTGCGGCCAAAAGCCACGTGTATAAGATACACCTGCAAAGGCGGCACAA<br>CCCCAGTGCCACGTTGTGAGTTGGATAGTTGTGGAAAGAGTCAAATGGCTCTCCTCAAGCGTATTCAACAAGGGGCTGAAGGATGCCCAGAAGGTACCCCATT<br>GTATGGGATCTGATCTGGGGCCTCGGTACACATGCTTTACATGTGTTTAGTCGAGGTTAAAAAAACGTCTAGGCCCCCCGAACCACGGGGACGTGGTTTTCCT<br>TTGAAAAACACGATGATAATATGGCCACAACCATGGTGAGCAAGGGCGAGGAGCTGTTCACCGGGGTGGTGCCCATCCTGGTCGAGCTGGACGGCGACGTAA<br>ACGGCCACAAGTTCAGCGTGTCCGGCGAGGGCGAGGGCGATGCCACCTACGGCAAGCTGACCCTGAAGTTCATCTGCACCACCGGCAAGCTGCCCGTGCCC<br>TGGCCCACCCTCGTGACCACCCTGACCTACGGCGTGCAGTGCTTCAGCCGCTACCCCGACCACATGAAGCAGCACGACTTCTTCAAGTCCGCCATGCCCGAA<br>GGCTACGTCCAGTTATTCTACATTGTTGTCATTATTACCAAATATTGGGGAAAATACAACTTACAGACCAATCTCAGGAGTTAAATGTTACTACGAAGGCAAATGAA<br>CTATGCGTAATGAACCTGGTAGGCATTAGCGGCCGCTCGAGTCTAGAGGGCCCGTTTAAACCCGCTGATCAGCCTCGACTGTGCCTTCTAGTTGCCAGCCATC<br>TGTTGTTTGCCCCTCCCCCGTGCCTTCCTTGACCCTGGAAGGTGCCACTCCC |
| Laccase2 Upper Band | AGCGCACCATCTTCTTCAAGGACGACGGCAACTACAAGACCCGCGCCGAGGTGAAGTTCGAGGGCGACACCCTGGTGAACCGCATCGAGCTGAAGGGCATCG<br>ACTTCAAGGAGGACGGCAACATCCTGGGGCACAAGCTGGAGTACAACCACAACAGCCACAACGTCTATATCATGGCCGACAAGCAGAAGAACGGCATCAAGGT<br>GAACTTCAAGATCCGCCACAACATCGAGGACGGCAGCGTGCAGCTCGCCGACCACTACCAGCAGAACACCCCCATCGGCGACGGCCCCGTGCTGCTGCCCGA<br>CAACCACTACCTGAGCACCCAGTCCGCCCTGAGCAAAGACCCCAACGAGAAGCGCGATCACATGGTCCTGCTGGAGTTCGTGACCGCCGCCGGGATCACTCT<br>CGGCATGGACGAGCTGTACAAGTAAGGATCTGAATTCTGCAGTCGACGATCCGCCCCTCTCCCTCCCCCCCCCCTAACGTTACTGGCCGAAGCCGCTTGGAATA<br>AGGCCGGTGTGCGTTTGTCTATATGTTATTTTCCACCATATTGCCGTCTTTTGGCAATGTGAGGGCCCGGAAACCTGGCCCTGTCTTCTTGACGAGCATTCCTA<br>GGGGTCTTTCCCCTCTCGCCAAAGGAATGCAAGGTCTGTTGAATGTCGTGAAGGAAGCAGTTCCTCTGGAAGCTTCTTGAAGACAAACAACGTCTGTAGCGAC<br>CCTTTGCAGGCAGCGGAACCCCCCACCTGGCGACAGGTGCCTCTGCGGCCAAAAGCCACGTGTATAAGATACACCTGCAAAGGCGGCACAACCCCAGTGCCA<br>CGTTGTGAGTTGGATAGTTGTGGAAAGAGTCAAATGGCTCTCCTCAAGCGTATTCAACAAGGGGCTGAAGGATGCCCAGAAGGTACCCCATTGTATGGGATCT<br>GATCTGGGGCCTCGGTACACATGCTTTACATGTGTTTAGTCGAGGTTAAAAAAACGTCTAGGCCCCCCGAACCACGGGGACGTGGTTTTCCTTTGAAAAACACG<br>ATGATAATATGGCCACAACCATGGTGAGCAAGGGCGAGGAGCTGTTCACCGGGGTGGTGCCCATCCTGGTCGAGCTGGACGGCGACGTAAACGGCCACAAG<br>TTCAGCGTGTCCGGCGAGGGCGAGGGCGATGCCACCTACGGCAAGCTGACCCTGAAGTTCATCTGCACCACCGGCAAGCTGCCCGTGCCCTGGCCCACCCT<br>CGTGACCACCCTGACCTACGGCGTGCAGTGCTTCAGCCGCTACCCCGACCACATGAAGCAGCACGACTTCTTCAAGTCCGCCATGCCCGAAGGCTACGTCCA<br>GGTAAGTATTCAAAATTCCAAAATTTTTTACTAGAAATATTCGATTTTTTAATAGGCAGTTTCTATACTATTGTATACTATTGTAGATTCGTTGAAAAGTATGTAACAG<br>GAAGAATAAAGCATTTCCGACCATGTAAAGTATATATATTCTTAATAAGGATCAATAGCCGAGTCGATCTCGCCATGTCCGTCTGTCTTATTGTTTTATTACCGCC<br>GAGACATCAGGAACTATAAAAGCTAGAAGGATGAGTTTTAGCATACAGATTCTAGAGACAAGGACGCAGAGCAAGTTTGTTGATCCATGCTGCCACGCTTTAACT<br>TTCTCAAATTGCCCAAAACTGCCATGCCCACATTTTTGAACTATTTTCGAAATTTTTTCATAATTGTATTACTCGTGTAAATTTCCATCAATTTGCCAAAAAACTTTTT<br>GTCACGCGTTAACGCCCTAAAGCCGCCAATTTGGTCACGCCCACACTATTGAGCAATTATCAAATTTTTTCTCATTTTATTCCCCAATATCTATCGATATCCCCGAT<br>TATGAAATTATTAAATTTCGCGTTCGCATTCACACTAGCTGAGTAACGAGTATCTGATAGTTGGGGAAATCGACTTATTTTTTATATACAATGAAAATGAATTTAATC<br>ATATGAATATCGATTATAGCTTTTTATTTAATATGAATATTTATTTGGGCTTAAGGTGTAACCTCCTCGACATAAGACTCACATGGCGCAGGCACATTGAAGACAAA<br>AATACTCATTGTCGGGTCTCGCACCCTCCAGCAGCACCTAAAATTATGTCTTCAATTATTGCCAACATTGGAGACACAATTAGTCTGTGGCACCTCAGGCGGCCG<br>CTCGAGTCTAGAGGGCCCGTTTAAACCCGCTGATCAGCCTCGACTGTGCCTTCTAGTTGCCAGCCATCTGTTGTTTGCCCCTCCCCCGTGCCTTCCTTGACCC<br>TGG |
| Laccase2 Lower Band | CTACGTCCAGGAGCGCACCATCTTCTTCAAGGACGACGGCAACTACAAGACCCGCGCCGAGGTGAAGTTCGAGGGCGACACCCTGGTGAACCGCATCGAGCT<br>GAAGGGCATCGACTTCAAGGAGGACGGCAACATCCTGGGGCACAAGCTGGAGTACAACCACAACAGCCACAACGTCTATATCATGGCCGACAAGCAGAAGAAC<br>GGCATCAAGGTGAACTTCAAGATCCGCCACAACATCGAGGACGGCAGCGTGCAGCTCGCCGACCACTACCAGCAGAACACCCCCATCGGCGACGGCCCCGTG<br>CTGCTGCCCGACAACCACTACCTGAGCACCCAGTCCGCCCTGAGCAAAGACCCCAACGAGAAGCGCGATCACATGGTCCTGCTGGAGTTCGTGACCGCCGCC<br>GGGATCACTCTCGGCATGGACGAGCTGTACAAGTAAGGATCTGAATTCTGCAGTCGACGATCCGCCCCTCTCCCTCCCCCCCCCCTAACGTTACTGGCCGAAG<br>CCGCTTGGAATAAGGCCGGTGTGCGTTTGTCTATATGTTATTTTCCACCATATTGCCGTCTTTTGGCAATGTGAGGGCCCGGAAACCTGGCCCTGTCTTCTTGA<br>CGAGCATTCCTAGGGGTCTTTCCCCTCTCGCCAAAGGAATGCAAGGTCTGTTGAATGTCGTGAAGGAAGCAGTTCCTCTGGAAGCTTCTTGAAGACAAACAACG<br>TCTGTAGCGACCCTTTGCAGGCAGCGGAACCCCCCACCTGGCGACAGGTGCCTCTGCGGCCAAAAGCCACGTGTATAAGATACACCTGCAAAGGCGGCACAA<br>CCCCAGTGCCACGTTGTGAGTTGGATAGTTGTGGAAAGAGTCAAATGGCTCTCCTCAAGCGTATTCAACAAGGGGCTGAAGGATGCCCAGAAGGTACCCCATT<br>GTATGGGATCTGATCTGGGGCCTCGGTACACATGCTTTACATGTGTTTAGTCGAGGTTAAAAAAACGTCTAGGCCCCCCGAACCACGGGGACGTGGTTTTCCT<br>TTGAAAAACACGATGATAATATGGCCACAACCATGGTGAGCAAGGGCGAGGAGCTGTTCACCGGGGTGGTGCCCATCCTGGTCGAGCTGGACGGCGACGTAA<br>ACGGCCACAAGTTCAGCGTGTCCGGCGAGGGCGAGGGCGATGCCACCTACGGCAAGCTGACCCTGAAGTTCATCTGCACCACCGGCAAGCTGCCCGTGCCC<br>TGGCCCACCCTCGTGACCACCCTGACCTACGGCGTGCAGTGCTTCAGCCGCTACCCCGACCACATGAAGCAGCACGACTTCTTCAAGTCCGCCATGCCCGAA<br>GGCTACGTCCAGGTGTAACCTCCTCGACATAAGACTCACATGGCGCAGGCACATTGAAGACAAAAATACTCATTGTCGGGTCTCGCACCCTCCAGCAGCACCTA<br>AAATTATGTCTTCAATTATTGCCAACATTGGAGACACAATTAGTCTGTGGCACCTCAGGCGGCCGCTCGAGTCTAGAGGGCCCGTTTAAACCCGCTGATCAGCC<br>TCGACTGTGCCTTCTAGTTGCCAGCCATCTGTTGTTTGCCCCTCCCCCGTGCCTTCCTTGACCCTGGAAGGTGCCACTCCC |
| POLR2A | CTACGTCCAGGAGCGCACCATCTTCTTCAAGGACGACGGCAACTACAAGACCCGCGCCGAGGTGAAGTTCGAGGGCGACACCCTGGTGAACCGCATCGAGCT<br>GAAGGGCATCGACTTCAAGGAGGACGGCAACATCCTGGGGCACAAGCTGGAGTACAACCACAACAGCCACAACGTCTATATCATGGCCGACAAGCAGAAGAAC<br>GGCATCAAGGTGAACTTCAAGATCCGCCACAACATCGAGGACGGCAGCGTGCAGCTCGCCGACCACTACCAGCAGAACACCCCCATCGGCGACGGCCCCGTG<br>CTGCTGCCCGACAACCACTACCTGAGCACCCAGTCCGCCCTGAGCAAAGACCCCAACGAGAAGCGCGATCACATGGTCCTGCTGGAGTTCGTGACCGCCGCC<br>GGGATCACTCTCGGCATGGACGAGCTGTACAAGTAAGGATCTGAATTCTGCAGTCGACGATCCGCCCCTCTCCCTCCCCCCCCCCTAACGTTACTGGCCGAAG<br>CCGCTTGGAATAAGGCCGGTGTGCGTTTGTCTATATGTTATTTTCCACCATATTGCCGTCTTTTGGCAATGTGAGGGCCCGGAAACCTGGCCCTGTCTTCTTGA<br>CGAGCATTCCTAGGGGTCTTTCCCCTCTCGCCAAAGGAATGCAAGGTCTGTTGAATGTCGTGAAGGAAGCAGTTCCTCTGGAAGCTTCTTGAAGACAAACAACG<br>TCTGTAGCGACCCTTTGCAGGCAGCGGAACCCCCCACCTGGCGACAGGTGCCTCTGCGGCCAAAAGCCACGTGTATAAGATACACCTGCAAAGGCGGCACAA<br>CCCCAGTGCCACGTTGTGAGTTGGATAGTTGTGGAAAGAGTCAAATGGCTCTCCTCAAGCGTATTCAACAAGGGGCTGAAGGATGCCCAGAAGGTACCCCATT<br>GTATGGGATCTGATCTGGGGCCTCGGTACACATGCTTTACATGTGTTTAGTCGAGGTTAAAAAAACGTCTAGGCCCCCCGAACCACGGGGACGTGGTTTTCCT<br>TTGAAAAACACGATGATAATATGGCCACAACCATGGTGAGCAAGGGCGAGGAGCTGTTCACCGGGGTGGTGCCCATCCTGGTCGAGCTGGACGGCGACGTAA<br>ACGGCCACAAGTTCAGCGTGTCCGGCGAGGGCGAGGGCGATGCCACCTACGGCAAGCTGACCCTGAAGTTCATCTGCACCACCGGCAAGCTGCCCGTGCCC<br>TGGCCCACCCTCGTGACCACCCTGACCTACGGCGTGCAGTGCTTCAGCCGCTACCCCGACCACATGAAGCAGCACGACTTCTTCAAGTCCGCCATGCCCGAA<br>GGCTACGTCCAGGGCGGCCGCTCGAGTCTAGAGGGCCCGTTTAAACCCGCTGATCAGCCTCGACTGTGCCTTCTAGTTGCCAGCCATCTGTTGTTTGCCCCT<br>CCCCCGTGCCTTCCTTGACCCTGGAAGGTGCCACTCCC |
| IGF2BP1 | CGCACCATCTTCTTCAAGGACGACGGCAACTACAAGACCCGCGCCGAGGTGAAGTTCGAGGGCGACACCCTGGTGAACCGCATCGAGCTGAAGGGCATCGAC<br>TTCAAGGAGGACGGCAACATCCTGGGGCACAAGCTGGAGTACAACTACAACAGCCACAACGTCTATATCATGGCCGACAAGCAGAAGAACGGCATCAAGGTGA<br>ACTTCAAGATCCGCCACAACATCGAGGACGGCAGCGTGCAGCTCGCCGACCACTACCAGCAGAACACCCCCATCGGCGACGGCCCCGTGCTGCTGCCCGACA<br>ACCACTACCTGAGCACCCAGTCCGCCCTGAGCAAAGACCCCAACGAGAAGCGCGATCACATGGTCCTGCTGGAGTTCGTGACCGCCGCCGGGATCACTCTCG<br>GCATGGACGAGCTGTACAAGTAAGGATCTGAATTCTGCAGTCGACGATCCGCCCCTCTCCCTCCCCCCCCCCCTAACGTTACTGGCCGAAGCCGCTTGGAATA<br>AGGCCGGTGTGCGTTTGTCTATATGTTATTTTCCACCATATTGCCGTCTTTTGGCAATGTGAGGGCCCGGAAACCTGGCCCTGTCTTCTTGACGAGCATTCCTA<br>GGGGTCTTTCCCCTCTCGCCAAAGGAATGCAAGGTCTGTTGAATGTCGTGAAGGAAGCAGTTCCTCTGGAAGCTTCTTGAAGACAAACAACGTCTGTAGCGAC<br>CCTTTGCAGGCAGCGGAACCCCCCACCTGGCGACAGGTGCCTCTGCGGCCAAAAGCCACGTGTATAAGATACACCTGCAAAGGCGGCACAACCCCAGTGCCA<br>CGTTGTGAGTTGGATAGTTGTGGAAAGAGTCAAATGGCTCTCCTCAAGCGTATTCAACAAGGGGCTGAAGGATGCCCAGAAGGTACCCCATTGTATGGGATCT<br>GATCTGGGGCCTCGGTACACATGCTTTACATGTGTTTAGTCGAGGTTAAAAAAACGTCTAGGCCCCCCGAACCACGGGGACGTGGTTTTCCTTTGAAAAACACG<br>ATGATAATATGGCCACAACCATGGTGAGCAAGGGCGAGGAGCTGTTCACCGGGGTGGTGCCCATCCTGGTCGAGCTGGACGGCGACGTAAACGGCCACAAG<br>TTCAGCGTGTCCGGCGAGGGCGAGGGCGATGCCACCTACGGCAAGCTGACCCTGAAGTTCATCTGCACCACCGGCAAGCTGCCCGTGCCCTGGCCCACCCT<br>CGTGACCACCCTGACCTACGGCGTGCAGTGCTTCAGCCGCTACCCCGACCACATGAAGCAGCACGACTTCTTCAAGTCCGCCATGCCCGAAGGCTACGTCCA<br>GGTAAGTCTCGACGGTACCGCGGGCCCGGGATCCCAAATAAGATGCCCTCAGACTTCTTTCCTTCCCATCTTCTTCTGTGGACCTTAAAAAGGGAAGACGAAGT<br>CAACAGAAATCCAGTTTGATGTCAATTTGAAAATAGCTCTGCAGCCCTTTATCCTGCATGGAAGGCTTAAAGATGTGATGAGGACTTGACGGCTAACGGAAACT<br>GACCCAGCCTTTTTCCTACCTCAGGGGAAATACCCAACCTCAATTCCTCCATGTTCAGCTTGGAAAACAATAAAGACCAAATCCTTGAATTGGGCCTCTTCTAGAG<br>CGGCCGCTCGAGTCTAGAGGGCCCGTTTAAACCCGCTGATCAGCCTCGACTGTGCCTTCTAGTTGCCAGCCATCTGTTGTTTGCCCCTCCCCCGTGCCTTCC<br>TTGACCCTGGA |

#### SUPPLEMENTARY METHODS

##### Expression plasmid construction

To generate circRNA expression plasmids, the following sequences were cloned between the EcoRI and NotI restriction sites in **pcDNA3.1(+) CircRNA Mini Vector** (Addgene #60648). Lowercase letters denote exon sequences. The sequence that is removed following digestion with PaqCI is denoted in blue.

**pcDNA3.1(+) circRNA Mini Scarless MCS**

AAAGTGCTGAGATTACAGGCGTGAGCCACCACCCCCGGCCCACTTTTTGTAAAGGTACGTACTAATGACTTTTTTTTTATACTTCA
GacgggcaggtgttgtgtacgtcacctgcctcgGTAAGAAGCAAGGAAAAGAATTAGGCTCGGCACGGTAGCTCACACCTGTAATC
CCAGCA

**pcDNA3.1(+) ZKSCAN1 Scarless MCS**

CAGTGACAGTGGAGATTGTACAGTTTTTTCCTCGATTTGTCAGGATTTTTTTTTTTTTGACGGAGTTTAACTTCTTGTCTCCCAGG
TAGGAAGTGCAGTGGCGTAATCTCGGCTCACTACAACCTCCACCTCCTGGGTTCAAGCGTTTCTCCTGCCTCAGCTTTCCGAGTAG
CTGGGATTACAGGCGCCTGCCACCATGCCCTGCTGACTTTTGTATTTTTAGTAGAGACGGGGTTTCACCATGTTGGCCAGGCTGGT
CTTGAACTCCTGACCGCAGGCGATTGGCCTGCCTCGGCCTCCCAAAGTGCTGAGATTACAGGCGTGAGCCACCACCCCCGGCCTCA
GGAGCGTTCTGATAGTGCCTCGATGTGCTGCCTCCTATAAAGTGTTAGCAGCACAGATCACTTTTTGTAAAGGTACGTACTAATGA
CTTTTTTTTTATACTTCAGacgggcaggtgttgtgtacgtcacctgcctcgGTAAGAAGCAAGGTTTCATTTAGGGGAAGGGAAAT
GATTCAGGACGAGAGTCTTTGTGCTGCTGAGTGCCTGTGATGAAGAAGCATGTTAGTCCTGGGCAACGTAGCGAGACCCCATCTCT
ACAAAAAATAGAAAAATTAGCCAGGTATAGTGGCGCACACCTGTGATTCCAGCTACGCAGGAGGCTGAGGTGGGAGGATTGCTTGA
GCCCAGGAGGTTGAGGCTGCAGTGAGCTGGAATCATGCCACTACTCCAACCTGGGCAACACAGCAAGGACCCTGTCTCAAAAGCTA
CTTACAGAAAAGAATTAGGCTCGGCACGGTAGCTCACACCTGTAATCCCAGCACTTTGGGAGGCTGAGGCGGGCAGATCACTTGAG
GTCAGGAGTTTGAGACCAGCCTGGCCAACATGGTGAAACCTTGTCTCTACTAAAAATATGAAAATTAGCCAGGCATGGTGGCACAT
TCCTGTAATCCCAGCTACTCGGGAGGCTGAGGCAGGAGAATCACTTGAACCCAGGAGGTGGAGGTTGCAGTAAGCCGAGATCGTAC
CACTGTGCTCTAGCCTTGGTGACAGAGCGAGACTGTCTTAAAAAAAAAAAAAAAAAAAAGAATTAATTAAAAATTTAAAAAAAAATG
AAAAAAAGCTGCATGCTTGTTTTTTGTTTTTAGTTATTCTACATTGTTGTCATTATTACCAAATATTGGGGAAAATACAACTTACA
GACCAATCTCAGGAGTTAAATGTTACTACGAAGGCAAATGAACTATGCGTAATGAACCTGGTAGGCATTA

**pcDNA3.1(+) Laccase2 Scarless MCS**

CATTGAGAAATGACTGAGTTCCGGTGCTCTCAAGTCATTGATCTTTGTCGACTTTTATTTGGTCTCTGTAATAACGACTTCAAAAA
CATTAAATTCTGTTGCGAAGCCAGTAAGCTACAAAAAGAAAAAACAAGAGAGAATGCTATAGTCGTATAGTATAGTTTCCCGACTA
TCTGATACCCATTACTTATCTAGGGGGAATGCGAACCCAAAATTTTATCAGTTTTCTCGGATATCGATAGATATTGGGGAATAAAT
TTAAATAAATAAATTTTGGGCGGGTTTAGGGCGTGGCAAAAAGTTTTTTGGCAAATCGCTAGAAATTTACAAGACTTATAAAATTA
TGAAAAAATACAACAAAATTTTAAACACGTGGGCGTGACAGTTTTGGGCGGTTTTAGGGCGTTAGAGTAGGCGAGGACAGGGTTAC
ATCGACTAGGCTTTGATCCTGATCAAGAATATATATACTTTATACCGCTTCCTTCTACATGTTACCTATTTTTCAACGAATCTAGT
ATACCTTTTTACTGTACGATTTATGGGTATAATAATAAGCTAAATCGAGACTAAGTTTTATTGTTATATATATTTTTTTTATTTAT
GCAGacgggcaggtgttgtgtacgtcacctgcctcgGTAAGTATTCAAAATTCCAAAATTTTTTACTAGAAATATTCGATTTTTTA
ATAGGCAGTTTCTATACTATTGTATACTATTGTAGATTCGTTGAAAAGTATGTAACAGGAAGAATAAAGCATTTCCGACCATGTAA
AGTATATATATATTCTTAATAAGGATCAATAGCCGAGTCGATCTCGCCATGTCCGTCTGTCTTATTGTTTTATTACCGCCGAGACATC
AGGAACTATAAAAGCTAGAAGGATGAGTTTTAGCATACAGATTCTAGAGACAAGGACGCAGAGCAAGTTTGTTGATCCATGCTGCC
ACGCTTTAACTTTCTCAAATTGCCCAAAACTGCCATGCCCACATTTTTGAACTATTTTCGAAATTTTTTCATAATTGTATTACTCG
TGTAAATTTCCATCAATTTGCCAAAAAACTTTTTGTCACGCGTTAACGCCCTAAAGCCGCCAATTTGGTCACGCCCACACTATTGA
GCAATTATCAAATTTTTTCTCATTTTATTCCCCAATATCTATCGATATCCCCGATTATGAAATTATTAAATTTCGCGTTCGCATTC
ACACTAGCTGAGTAACGAGTATCTGATAGTTGGGGAAATCGACTTATTTTTTATATACAATGAAAATGAATTTAATCATATGAATA
TCGATTATAGCTTTTTATTTAATATGAATATTTATTTGGGCTTAAGGTGTAACCTCCTCGACATAAGACTCACATGGCGCAGGCAC
ATTGAAGACAAAAATACTCATTGTCGGGTCTCGCACCCTCCAGCAGCACCTAAAATTATGTCTTCAATTATTGCCAACATTGGAGA
CACAATTAGTCTGTGGCACCTCAG

**pcDNA3.1(+) POLR2A Scarless MCS**

AGGCACGTAGCAGGCCAGGGCCTCTCTCAGCCACCTGAGCAGAAAGCTTTCCAAGATAGGGCAGGCTGGGTTAGGCCATCTGAGTC
TGTCTCGTTCATTGGGATCCAGACTTGACTGTCTTGTTAAAGGCTGTTGCTGCCCAGGTGTGCAGGGAGCTGTTGGTCTCTGGCAT
TCAGGGTGGGGGTGGTATAAACCCGGGGCAGCTTGCATATGGCAGGGAAGAGGGATCCGTGGAGGAACAGTGCAGAAGGCTTTATG
TTCAGAATCTCTCTTGCTTTTCTTCTAGACTGAGTTCCTTGAGATTGGTGAATGCTGTGTATTATTCATCCCTGATAACCTGGTGT
TTGGCCCAGGGCCTTGTCCAGAGGAGTGTTTGATAAGTGTTTCAAGTGAATTAGCACCACGATGTCATCTCTTTTCAGTTTACAAA
GGACGGACACCCCCTGACCCGGTCTCAGAAAGCCTGAAAGCAGAATTAGTCATTAGAAGGGTGGTTGGCTTGGTCGGCATAGACTT

TGAGCAGAAAGAGGTTGAAAATGTTGAGCCTGATTTCTCTTAGGCCCCTCTGCAGTGTCTGTTGTGGAGGCCAGATACGTAACTGC
TTCCGCTTTTTTTGGTGTCATTCAAGGTGAGCAAATCCCCTTCATGTTTCTCACCAGACAATGCAGCTGATGAGGTTCCAGCTTTG
CAAATGTAGTCATCCATGAGGACTGTCTTCCTGAGATTTCATCAGGCTCGAGTGGACTTGCAAAGGACTTTAGGTCCATTGTCCTT
TTATTCTTAGATACCTCTTTCACTGAGACCTTTTCCTTACCTCACCTCTCTAGacgggcaggtgttgtgtacgtcacctgcctcgT
GTGTGGTCCAAATGGAAACCTGGCTTAAGTGGGCAGTGGGGCTCTGGGGTGCAAGGTGGAGGCTAGAGAGGAAGAGCTGTGTTTTT
TTTCCTGACTTACCCAGCAGTGGTCTGTGAGATTGTCTTTTCTGGTGGGCGAACAAAAAGGGGGTTAGGAAAACTCAGGCCAAAAA
AGTGTAAGGCGTTAATTCCCCATTTAATTCCTTAAAATTTCATGTAATACCAGGTATTGCCTGTAAAGGAAAGATAAAGGGAAAAA
TAAGTAAGACCTTGTTAAAATTTTATTTTTCTATTTTAACCTTCACTTATTTCCTAATTATTAAAAGAAATTTATGCTTATTGTTA
AGAACAAAAAAAATTTCAGTATTACAATGAATTTTTAATTAAAAGTTTTTGGCCTGATGAAATCTCAGGAAGACAGTCCTCATGGAT
GACTACATTTGCAAAGCTGGAACCTCATCAGCTGCATTGTCTGGTGAGAAACATGAAGGGGATTTGCTCACCTTGAATGAGACCAA
AAAAAGCGGAAGCAGTTACGTATCTGGCCTCCACAACAGACACTGCAGAGGGGCCTAAGAGAAATCAGGCTCAACATTTTCAACCT
CTTTCTGCTCAAAGTCTATGCCGACCAAGCCAACCACCCTTCTAATGACTAATTCTGCTTTCAGGCTTTCTGAGACCGGGTCAGGG
GGTGTCCTTAAAGGTTGGAAAAAACTTTTCCTGTCATCTTTGCCTCCAAAATCTGGCTTTCTCCCTTGGGCAGGGAAACCTCCCCA
ACATTTCTCTATCATCCCTGAGATGTGGGGCCTGCACTCTGACTTCTGTCTGCCTTACTCTTTGTCTTACAGG

#### pcDNA3.1(+) ciRS-7 Scarless MCS

CTGTAAGAGTAGTCTCATGATGTCTTTTAATTTAATATCTAGGCCCTAATACACAACTCTTTGAATATTGCTACTATCTTTCAGCA
TGACTTTCCAATACCATTAAGATTAAGCCTTTCTGTTCTTCAGTTTCTCTAACTCAAGTCTTCTAAGCTCCACAGCTTCCAACATC
TTTAATTACCAGATCTTCCCAATATTATAACATCCGTGCATTCCAGTCCCTGTCCTGCAGCATCTCCAATGCCCTTGTTTTCTAAA
CTTCCCAGTATTCAAGTCATTAAAAATTTAGGTCTTCTAATATCTCCAACAACCCATCTTCAAAGTCCAGGTCTTGTCTTTCAGAA
TCCAGAGTTCTAAATTACAAGTCTTCCATGGTCCTAATCCTCCAATGTCTCCTATATACAGGTCTTCTAGAGGCTATGTATTTCAT
AGTGTTTAAGTCTAGATTCTTCAATGTCTTCCAGCACTTACATCATTCAGTTTTTACATTTTTAAATGTCTCCTACTTTCTAGTTT
TACAGTGTATTTTTATCGCCAATTCCTCCAATGCCCAAGTCTTTAAATACCTGCAAGAACTTTCAGGTACATCAATATCCTCAATG
TCTGAATGAATAGACCTCTTAATATCCTGGTCTCATATTGTCCAGGCTGCCAGTATTCTTGACATCCAGCATCTTTAAAATTTAAT
TCTTCCAAACTGTCTTTTTGACATAAATATGTCAATATCCATGCTTACCAATGTCTCCCAATATCCAGGGCTTCTCATTTTAGGAT
TTATAAATGTCTACATCTGCTGTTGTTTATATCTGCCAGTATTTAGGAGTTCCAAGGTACAGGACTTACAATATCTCTAATTTCCA
GAGATTTCAGTGTCTATGACTTCCAATATTCAATATCCCAATGTTCAGGGCCTCCAGAAATATCAATGTCCAGAGTAACCAATGTC
TTCAAAATCCAAGGCTTGCAATGTCTCCAATATCCTGTGCTTACAATGTTTCCAATGTCCAGacgggcaggtgttgtgtacgtcac
ctgcctcgGTATTCTAACATGTTCTATAGTCCAGATCTTCCAGCCTTTCCCCAGGTCCAGGCCTTTTCAAATCTAGGCTTCCCCTG
TCTGAGCCTTCCCATGTTAAAATCACTCTACAGAAAAATCTTCCAACTCAACTATTTCTGCTTCTAGTTGAATTAGCTTCAAAAAC
ATTTAATGGGACATTCCTCCTTATAGACCAGATCTTATGAACCAACAGTACTGCAGTTTGGCGGCCGCTCGACTGGTTACTCTGGA
CATTGATATTTCTGGAGGCCCTGAACATTGGGATATTGAATATTGGAAGTCATAGACACTGAAATCTCTGGAAATTAGAGATATTG
TAAGTCCTGTACCTTGGAACTCCTAAATACTGGCAGATATAAACAACAGCAGATGTAGACATTTATAAATCCTAAAATGAGAAGCC
CTGGATATTGGGAGACATTGGTAAGCATGGATATTGACATATTTATGTCAAAAAGACAGTTTGGAAGAATTAAATTTTAAAGATGC
TGGATGTCAAGAATACTGGCAGCCTGGACAATATGAGACCAGGATATTAAGAGGTCTATTCATTCAGACATTGAGGATATTGATGT
ACCTGAAAGTTCTTGCAGGTATTTAAAGACTTGGGCATTGGAGGAATTGGCGATAAAAATACACTGTAAAACTAGAAAGTAGGAGA
CATTTAAAAATGTAAAAACTGAATGATGTAAGTGCTGGAAGACATTGAAGAATCTAGACTTAAACACTATGAAATACATAGCCTCT
AGAAGACCTGTATATAGGAGACATTGGAGGATTAGGACCATGGAAGACTTGTAATTTAGAACTCTGGATTCTGAAAGACAAGACCT
GGACTTTGAAGATGGGTTGTTGGAGATATTAGAAGACCTAAATTTTTAATGACTTGAATACTGGGAAGTTTAGAAAACAAGGGCAT
TGGAGATGCTGCAGGACAGGGACTGGAATGCACGGATGTTATAATATTGGGAAGATCTGGTAATTAAAGATGTTGGAAGCTGTGGA
GCTTAGAAGACTTGAGTTAGAGAAACTGAAGAACAGAAAGGCTTAATCTTAATGGTATTGGAAAGTCATGCTGAAAGATAGTAGCA
ATATTCAAAGAGTTGTGTATTAGGGCCTAGATATTAAATTAAAAGACATCATGAGACTACTCTTACAGGTCGAGCATGCATCTAGA

#### pcDNA3.1(+) IGF2BP1 Scarless MCS

GACTGAACATGGAGGAATTGAGGTTGGGTATTTCCCCTGAGGTAGGAAAAAGGCTGGGTCAGTTTCCCGTTAGCCGTCAAGTCCTC
ATCACATCTTTAAGCCTTCCATGCAGGATAAAGGGCTGCAGAGCTATTTTCAAATTGACATCAAACTGGATTTCTGTTGACTTCGT
CTTCCCTTTTTAAGGTCCACAGAAGAAGATGGGAAGGAAAGAAGTCTGAGGGCATCTTATTTGCACTCCGCTGTCATTTCTAAGGA
AGGCCTTTAATGCCAAATTCTCATCTTTTATGTCCCCACTAAATCCTAAGGTTCTTGAACTTCTGATCAGACAGCCAAAAAATGAA
CCATCAACTAGCTTAACCTAACATATGTGAGGATAGAGGACTGGGACAGCTCTCTGGGCCACTGGAGAGTCAGACAGGCCTGCCCT
CTGTGTGACTTGACCGCGGTCTCTTTCTTCCAGacgggcaggtgttgtgtacgtcacctgcctcgGTAAGTCTCGACGGTACCGCG
GGCCCGGGATCCCAAATAAGATGCCCTCAGACTTCTTTCCTTCCCATCTTCTTCTGTGGACCTTAAAAAGGGAAGACGAAGTCAAC
AGAAATCCAGTTTGATGTCAATTTGAAAATAGCTCTGCAGCCCTTTATCCTGCATGGAAGGCTTAAAGATGTGATGAGGACTTGAC
GGCTAACGGGAAACTGACCCAGCCTTTTTCCTACCTCAGGGGAAATACCCAACCTCAATTCCTCCATGTTCAGCTTGGAAAACAAT
AAAGACCAAATCCTTGAATTGGGCCTCTTCTAGA

To generate **pcDNA3.1(+) Tornado PaqCI MCS**, a sequence containing the Tornado ribozymes (based on pAV-U6+27-Tornado-Broccoli (Addgene #124360)) and PaqCI restriction sites was synthesized and cloned between the EcoRI and NotI restriction sites in pcDNA3.1(+) CircRNA

Mini Vector (Addgene #60648). The sequence that is removed following digestion with PaqCI is denoted in blue.

GGCCGCACTCGCCGGTCCCAAGCCCGGATAAAATGGGAGGGGGCGGGAAACCGCCTAACCATGCCGAGTGCGGCCGCacgggcagg
tgttgtgtacgtcacctgcctcgGTGGCCGCGGTCGGCGTGGACTGTAGAACACTGCCAATGCCGGTCCCAAGCCCGGATAAAAGT
GGAGGGTACAGTCCACGC

To generate the following plasmids:

**pcDNA3.1(+) circRNA Mini-Split eGFP**
**pcDNA3.1(+) circRNA Mini Scarless-Split eGFP**
**pcDNA3.1(+) ZKSCAN1 Scarless-Split eGFP**
**pcDNA3.1(+) Laccase2 Scarless-Split eGFP**
**pcDNA3.1(+) POLR2A Scarless-Split eGFP**
**pcDNA3.1(+) ciRS-7 Scarless-Split eGFP**
**pcDNA3.1(+) IGF2BP1 Scarless-Split eGFP**
**pcDNA3.1(+) Tornado PaqCI-Split eGFP**

the following exonic sequence was cloned into each of the MCS plasmids listed above. PaqCI restriction sites were used, except to generate **pcDNA3.1(+) circRNA Mini-Split eGFP**, where EcoRV and SacII restriction sites were used to clone into pcDNA3.1(+) CircRNA Mini Vector (Addgene #60648). eGFP start and stop codons are denoted

GAGCGCACCATCTTCTTCAAGGACGACGGCAACTACAAGACCCGCGCCGAGGTGAAGTTCGAGGGCGACACCCTGGTGAACCGCAT
CGAGCTGAAGGGCATCGACTTCAAGGAGGACGGCAACATCCTGGGGCACAAGCTGGAGTACAACCACAACAGCCACAACGTCTATA
TCATGGCCGACAAGCAGAAGAACGGCATCAAGGTGAACTTCAAGATCCGCCACAACATCGAGGACGGCAGCGTGCAGCTCGCCGAC
CACTACCAGCAGAACACCCCCATCGGCGACGGCCCCGTGCTGCTGCCCGACAACCACTACCTGAGCACCCAGTCCGCCCTGAGCAA
AGACCCCAACGAGAAGCGCGATCACATGGTCCTGCTGGAGTTCGTGACCGCCGCCGGGATCACTCTCGGCATGGACGAGCTGTACA
AGTAAGGATCTGAATTCTGCAGTCGACGATCCGCCCCTCTCCCTCCCCCCCCCCCTAACGTTACTGGCCGAAGCCGCTTGGAATAA
GGCCGGTGTGCGTTTGTCTATATGTTATTTTCCACCATATTGCCGTCTTTTGGCAATGTGAGGGCCCGGAAACCTGGCCCTGTCTT
CTTGACGAGCATTCCTAGGGGTCTTTCCCCTCTCGCCAAAGGAATGCAAGGTCTGTTGAATGTCGTGAAGGAAGCAGTTCCTCTGG
AAGCTTCTTGAAGACAAACAACGTCTGTAGCGACCCTTTGCAGGCAGCGGAACCCCCCACCTGGCGACAGGTGCCTCTGCGGCCAA
AAGCCACGTGTATAAGATACACCTGCAAAGGCGGCACAACCCCAGTGCCACGTTGTGAGTTGGATAGTTGTGGAAAGAGTCAAATG
GCTCTCCTCAAGCGTATTCAACAAGGGGCTGAAGGATGCCCAGAAGGTACCCCATTGTATGGGATCTGATCTGGGGCCTCGGTACA
CATGCTTTACATGTGTTTAGTCGAGGTTAAAAAAACGTCTAGGCCCCCCGAACCACGGGGACGTGGTTTTCCTTTGAAAAACACGA
TGATAATATGGCCACAACCATGGTGAGCAAGGGCGAGGAGCTGTTCACCGGGGTGGTGCCCATCCTGGTCGAGCTGGACGGCGACG
TAAACGGCCACAAGTTCAGCGTGTCCGGCGAGGGCGAGGGCGATGCCACCTACGGCAAGCTGACCCTGAAGTTCATCTGCACCACC
GGCAAGCTGCCCGTGCCCTGGCCCACCCTCGTGACCACCCTGACCTACGGCGTGCAGTGCTTCAGCCGCTACCCCGACCACATGAA
GCAGCACGACTTCTTCAAGTCCGCCATGCCCGAAGGCTACGTCCAG

To generate the following plasmids:

**pcDNA3.1(+) circRNA Mini-ZKSCAN1 Exons 2-3**
**pcDNA3.1(+) circRNA Mini Scarless-ZKSCAN1 Exons 2-3**
**pcDNA3.1(+) ZKSCAN1 Scarless-ZKSCAN1 Exons 2-3**
**pcDNA3.1(+) Laccase2 Scarless-ZKSCAN1 Exons 2-3**
**pcDNA3.1(+) POLR2A Scarless-ZKSCAN1 Exons 2-3**
**pcDNA3.1(+) ciRS-7 Scarless-ZKSCAN1 Exons 2-3**
**pcDNA3.1(+) Tornado PaqCI-ZKSCAN1 Exons 2-3**

the following sequence was cloned into each of the MCS plasmids listed above. PaqCI restriction sites were used, except to generate **pcDNA3.1(+) circRNA Mini-ZKSCAN1 Exons 2-3**, where EcoRV and SacII restriction sites were used to clone into pcDNA3.1(+) CircRNA Mini Vector (Addgene #60648). The intervening 221-nt intron sequence is shown in lowercase.

```
GAATAGTAAAGAAACACATCATAAAACCTCCCAGGACATAAAGGTGAGCACAGACCCTGTTTGGATCAAGTCAGTTCCTGGAGCCT
GAATGATGACTGCTGAATCACGGGAAGCCACGGGTCTGTCCCCACAGGCTGCACAGGAGAAGGATGGTATCGTAATAGTGAAGGTG
GAAGAGGAAGATGAGGAAGACCACATGTGGGGGCAGGATTCCACCCTACAGGACACGCCTCCTCCAGACCCAGAGATATTCCGCCA
ACGCTTCAGGCGCTTCTGTTACCAGAACACTTTTGGGCCCCGAGAGGCTCTCAGTCGGCTGAAGGAACTTTGTCATCAGTGGCTGC
GGCCAGAAATAAACACCAAGGAACAGATCCTGGAGCTTCTGGTGCTAGAGCAGTTTCTTTCCATCCTGCCCAAGGAGCTCCAGGTC
TGGCTGCAGGAATACCGCCCCGATAGTGGAGAGGAGGCCGTGACCCTTCTAGAAGACTTGGAGCTTGATTTATCAGGACAACAGgt
aaaaagaggtgaaacctattatgtgtgagcagggcacagacgttgaaactggagccaggagaagtattggcaggctttaggttatt
aggtggttactctgtcttaaaaatgttctggctttcttcctgcatccactggcatactcatggtctgtttttaaatattttaattc
ccatttacaaagtgatttacccacaagcccaacctgtctgtcttcagGTCCCAGGTCAAGTTCATGGACCTGAGATGCTCGCAAGG
GGGATGGTGCCTCTGGATCCAGTTCAGGAGTCCTCGAGCTTTGACCTTCATCACGAGGCCACCCAGTCCCACTTCAAACATTCGTC
TCGGAAACCCCGCCTCTTACAGTCACGAG
```

To generate the following plasmids:

**pcDNA3.1(+) circRNA Mini-ciRS-7**
**pcDNA3.1(+) circRNA Mini Scarless-ciRS-7**
**pcDNA3.1(+) ZKSCAN1 Scarless-ciRS-7**
**pcDNA3.1(+) Laccase2 Scarless-ciRS-7**
**pcDNA3.1(+) POLR2A Scarless-ciRS-7**
**pcDNA3.1(+) ciRS-7 Scarless-ciRS-7**
**pcDNA3.1(+) Tornado PaqCI-ciRS-7**

the following sequence was cloned into each of the MCS plasmids listed above. PaqCI restriction sites were used, except to generate **pcDNA3.1(+) circRNA Mini-ciRS-7**, where EcoRV and SacII restriction sites were used to clone into pcDNA3.1(+) CircRNA Mini Vector (Addgene #60648).

```
GGTTTCCGATGGCACCTGTGTCAAGGTCTTCCAACAACTCCGGGTCTTCCAGCGACTTCAAGTCTTCCAATAATCTCAAGGTCTTC
CAGATAATCCTGAGCTTCCAGAAAATCCACATCTTCCAGACAATCCATGTCTTCCGGACAATCCATGTCTTCCAAGAAGCTCCAAG
TCTTCCAGTAAATCAAGTCTTCCAGCAAATCCAGTCTTCCAGCAATTACTGGTCTTCCACCAAATCCAGATCTTCCAGGAAAATCC
ACGTCTTCCAGGAAATCCATGTCTTCCAATAATTTCAAGGTCTTCCATCAAATACAGATCTTCCAGCTAATCCATGTCTTCCAGAA
AAATCTGTGTCTTCCACCAAATCCAAGTCTTCCAGTAAATCTAGTTCTTCCAGAAAAATCTAGATCTTCCAGTCAATCAGTGTCTT
CCAGAAAGAAATCCAGGTCTTCCAGTCAATCAGTGTCTTCCAGAAAGAAATCCAGGTCTTCCAGTCAGTCAGTGTCTTCCAGAAAA
ATCTACGTCTTCCACCAAATCCAGGTCTTCCAGTCAATCCACATCTTCCGGAAAAAATCCAGGTCTTCCAGCCAATATATGTCTTC
CTGAAGATCCACGTCTTCCAGAAAATCCATGTCTTCCAGAAAATCCATGTCTTCCAGTAACCTCCCAGTCTTCCAGAAAATCCACG
TCTTCCCAACAATCCAAGTCTTCCGGATAATTTGGGTCTTCCTGAAAATCTACGTCTTCCAAAAAAGCCATGTCTTCCAGAAAATC
CACATCTTCCAATGGCCTCCAGGTCTTCCAGACTATCCATGTCTTCCAGAAAATCCTTGTCTTCCCTTAAATCTATAGCTTCCAAA
AAATCCGGGTCTTCCAGGAAATCCGTGTCTTCCAGCAAGTCCACGTCTTCCAACAAAGCCATGTCTTCCAGACTATCCATGTCTTC
CAGAAAATCCTTGTCTTCCCTCAAATCCATAGCTTCCGAAAAATCCAGGTCTTCCAGGAAATCCGTGTCTTCCAGCAAATCCACGT
CTTCCAACAAAGCCATGTCTTCCATCAAATTAATGTCTTCCAGCCTACTTGTGTCTTCCAACAAAGGTACGTCTTCCAACAAAGGT
ACGTCTTCCAACAAAGGTATGTCTTCCAACAAAGGTACGTCTTCCAGAAAATCCACGTCTTCCAACCAAGCCATGTCTTCCAGAAA
ATCCACGTCTTCCAGAAAATATATGTCTTCCAACTAAGCTACGTCTTCCAACAAATCCATGTCTTCCTATATCTCCAGGTCTTCCA
GCATCTCCAGGGCTTCCAGCATCTGCTCGTCTTCCAACATCTCCACGTCTTCCAGCATCTCTGTGTCTTCCAGCATCTTCATGTCT
TCCAACAACTACCCAGTCTTCCATCAACTGGCTCAATATCCATGTCTTCCAACGTCTCCAGTGTGCTGATCTTCTGACATTCAGGT
CTTCCAGTGTCTGCAATATCCAG
```

To generate the following plasmids:

**pcDNA3.1(+) circRNA Mini-NFASC Exons 26-27**
**pcDNA3.1(+) circRNA Mini Scarless-NFASC Exons 26-27**
**pcDNA3.1(+) ZKSCAN1 Scarless-NFASC Exons 26-27**
**pcDNA3.1(+) Laccase2 Scarless-NFASC Exons 26-27**
**pcDNA3.1(+) POLR2A Scarless-NFASC Exons 26-27**
**pcDNA3.1(+) ciRS-7 Scarless-NFASC Exons 26-27**
**pcDNA3.1(+) Tornado PaqCI-NFASC Exons 26-27**

the following sequence was cloned into each of the MCS plasmids listed above. PaqCI restriction sites were used, except to generate **pcDNA3.1(+) circRNA Mini-NFASC Exons 26-27**, where EcoRV and SacII restriction sites were used to clone into pcDNA3.1(+) CircRNA Mini Vector (Addgene #60648).

CCCCTGATGAGCAGTCCATATGGAACGTCACGGTGCTCCCCAACAGTAAATGGGCCAACATCACCTGGAAGCACAATTTCGGGCCC
GGAACTGACTTTGTGGTTGAGTACATCGACAGCAACCATACGAAAAAAACTGTCCCAGTTAAGGCCCAGGCTCAGCCTATACAGCT
GACAGACCTCTATCCCGGGATGACATACACGTTGCGGGTTTATTCCCGGGACAACGAGGGCATCAGCAGTACCGTCATCACCTTTA
TGACCAGTACAG

To generate the following expression plasmids with deletions of intronic repeat sequences, the following sequences were cloned between the EcoRI and NotI restriction sites in **pcDNA3.1(+) CircRNA Mini Vector** (Addgene #60648). The split eGFP ORF and BSJ scars (when applicable) are highlighted.

**pcDNA3.1(+) circRNA Mini-Split eGFP Delta Repeat**
AAAGTGCTGAGATTACAGGCGTGAGCCACCACCCCCGGCCCACTTTTTGTAAAGGTACGTACTAATGACTTTTTTTTTATACTTCA
GGATATCGAGCGCACCATCTTCTTCAAGGACGACGGCAACTACAAGACCCGCGCCGAGGTGAAGTTCGAGGGCGACACCCTGGTGA
ACCGCATCGAGCTGAAGGGCATCGACTTCAAGGAGGACGGCAACATCCTGGGGCACAAGCTGGAGTACAACCACAACAGCCACAAC
GTCTATATCATGGCCGACAAGCAGAAGAACGGCATCAAGGTGAACTTCAAGATCCGCCACAACATCGAGGACGGCAGCGTGCAGCT
CGCCGACCACTACCAGCAGAACACCCCCATCGGCGACGGCCCCGTGCTGCTGCCCGACAACCACTACCTGAGCACCCAGTCCGCCC
TGAGCAAAGACCCCAACGAGAAGCGCGATCACATGGTCCTGCTGGAGTTCGTGACCGCCGCCGGGATCACTCTCGGCATGGACGAG
CTGTACAAGTAAGGATCTGAATTCTGCAGTCGACGATCCGCCCCTCTCCCTCCCCCCCCCCTAACGTTACTGGCCGAAGCCGCTTG
GAATAAGGCCGGTGTGCGTTTGTCTATATGTTATTTTCCACCATATTGCCGTCTTTTGGCAATGTGAGGGCCCGGAAACCTGGCCC
TGTCTTCTTGACGAGCATTCCTAGGGGTCTTTCCCCTCTCGCCAAAGGAATGCAAGGTCTGTTGAATGTCGTGAAGGAAGCAGTTC
CTCTGGAAGCTTCTTGAAGACAAACAACGTCTGTAGCGACCCTTTGCAGGCAGCGGAACCCCCCACCTGGCGACAGGTGCCTCTGC
GGCCAAAAGCCACGTGTATAAGATACACCTGCAAAGGCGGCACAACCCCAGTGCCACGTTGTGAGTTGGATAGTTGTGGAAAGAGT
CAAATGGCTCTCCTCAAGCGTATTCAACAAGGGGCTGAAGGATGCCCAGAAGGTACCCCATTGTATGGGATCTGATCTGGGGCCTC
GGTACACATGCTTTACATGTGTTTAGTCGAGGTTAAAAAAACGTCTAGGCCCCCCGAACCACGGGGACGTGGTTTTCCTTTGAAAA
ACACGATGATAATATGGCCACAACCATGGTGAGCAAGGGCGAGGAGCTGTTCACCGGGGTGGTGCCCATCCTGGTCGAGCTGGACG
GCGACGTAAACGGCCACAAGTTCAGCGTGTCCGGCGAGGGCGAGGGCGATGCCACCTACGGCAAGCTGACCCTGAAGTTCATCTGC
ACCACCGGCAAGCTGCCCGTGCCCTGGCCCACCCTCGTGACCACCCTGACCTACGGCGTGCAGTGCTTCAGCCGCTACCCCGACCA
CATGAAGCAGCACGACTTCTTCAAGTCCGCCATGCCCGAAGGCTACGTCCAGCCGCGGAG

**pcDNA3.1(+) circRNA Mini Scarless-Split eGFP Delta Repeat**
AAAGTGCTGAGATTACAGGCGTGAGCCACCACCCCCGGCCCACTTTTTGTAAAGGTACGTACTAATGACTTTTTTTTTATACTTCA
GGAGCGCACCATCTTCTTCAAGGACGACGGCAACTACAAGACCCGCGCCGAGGTGAAGTTCGAGGGCGACACCCTGGTGAACCGCA
TCGAGCTGAAGGGCATCGACTTCAAGGAGGACGGCAACATCCTGGGGCACAAGCTGGAGTACAACCACAACAGCCACAACGTCTAT
ATCATGGCCGACAAGCAGAAGAACGGCATCAAGGTGAACTTCAAGATCCGCCACAACATCGAGGACGGCAGCGTGCAGCTCGCCGA
CCACTACCAGCAGAACACCCCCATCGGCGACGGCCCCGTGCTGCTGCCCGACAACCACTACCTGAGCACCCAGTCCGCCCTGAGCA
AAGACCCCAACGAGAAGCGCGATCACATGGTCCTGCTGGAGTTCGTGACCGCCGCCGGGATCACTCTCGGCATGGACGAGCTGTAC
AAGTAAGGATCTGAATTCTGCAGTCGACGATCCGCCCCTCTCCCTCCCCCCCCCCTAACGTTACTGGCCGAAGCCGCTTGGAATAA
GGCCGGTGTGCGTTTGTCTATATGTTATTTTCCACCATATTGCCGTCTTTTGGCAATGTGAGGGCCCGGAAACCTGGCCCTGTCTT
CTTGACGAGCATTCCTAGGGGTCTTTCCCCTCTCGCCAAAGGAATGCAAGGTCTGTTGAATGTCGTGAAGGAAGCAGTTCCTCTGG

AAGCTTCTTGAAGACAAACAACGTCTGTAGCGACCCTTTGCAGGCAGCGGAACCCCCCACCTGGCGACAGGTGCCTCTGCGGCCAA
AAGCCACGTGTATAAGATACACCTGCAAAGGCGGCACAACCCCAGTGCCACGTTGTGAGTTGGATAGTTGTGGAAAGAGTCAAATG
GCTCTCCTCAAGCGTATTCAACAAGGGGCTGAAGGATGCCCAGAAGGTACCCCATTGTATGGGATCTGATCTGGGGCCTCGGTACA
CATGCTTTACATGTGTTTAGTCGAGGTTAAAAAAACGTCTAGGCCCCCCGAACCACGGGGACGTGGTTTTCCTTTGAAAAACACGA
TGATAATATGGCCACAACCATGGTGAGCAAGGGCGAGGAGCTGTTCACCGGGGTGGTGCCCATCCTGGTCGAGCTGGACGGCGACG
TAAACGGCCACAAGTTCAGCGTGTCCGGCGAGGGCGAGGGCGATGCCACCTACGGCAAGCTGACCCTGAAGTTCATCTGCACCACC
GGCAAGCTGCCCGTGCCCTGGCCCACCCTCGTGACCACCCTGACCTACGGCGTGCAGTGCTTCAGCCGCTACCCCGACCACATGAA
GCAGCACGACTTCTTCAAGTCCGCCATGCCCGAAGGCTACGTCCAG

#### pcDNA3.1(+) ZKSCAN1 Scarless-Split eGFP Delta Repeat

CAGTGACAGTGGAGATTGTACAGTTTTTTCCTCGATTTGTCAGGATTTTTTTTTTTTGACGGAGTTTAACTTCTTGTCTCCCAGGT
AGGAAGTGCAGTGGCGTAATCTCGGCTCACTACAACCTCCACCTCCTGGGTTCAAGCGTTTCTCCTGCCTCAGCTTTCCGAGTAGC
TGGGATTACAGGCGCCTGCCACCATGCCCTGCTGACTTTTGTATTTTTAGTAGAGACGGGGTTTCACCATGTTGGCCAGGCTGGTC
TTGAACTCCTGACCGCAGGCGATTGGCCTGCCTCGGCCTCCCAAAGTGCTGAGATTACAGGCGTGAGCCACCACCCCCGGCCTCAG
GAGCGTTCTGATAGTGCCTCGATGTGCTGCCTCCTATAAAGTGTTAGCAGCACAGATCACTTTTTGTAAAGGTACGTACTAATGAC
TTTTTTTTTATACTTCAGGAGCGCACCATCTTCTTCAAGGACGACGGCAACTACAAGACCCGCGCCGAGGTGAAGTTCGAGGGCGA
CACCCTGGTGAACCGCATCGAGCTGAAGGGCATCGACTTCAAGGAGGACGGCAACATCCTGGGGCACAAGCTGGAGTACAACCACA
ACAGCCACAACGTCTATATCATGGCCGACAAGCAGAAGAACGGCATCAAGGTGAACTTCAAGATCCGCCACAACATCGAGGACGGC
AGCGTGCAGCTCGCCGACCACTACCAGCAGAACACCCCCATCGGCGACGGCCCCGTGCTGCTGCCCGACAACCACTACCTGAGCAC
CCAGTCCGCCCTGAGCAAAGACCCCAACGAGAAGCGCGATCACATGGTCCTGCTGGAGTTCGTGACCGCCGCCGGGATCACTCTCG
GCATGGACGAGCTGTACAAGTAAGGATCTGAATTCTGCAGTCGACGATCCGCCCCTCTCCCTCCCCCCCCCCTAACGTTACTGGCC
GAAGCCGCTTGGAATAAGGCCGGTGTGCGTTTGTCTATATGTTATTTTCCACCATATTGCCGTCTTTTGGCAATGTGAGGGCCCGG
AAACCTGGCCCTGTCTTCTTGACGAGCATTCCTAGGGGTCTTTCCCCTCTCGCCAAAGGAATGCAAGGTCTGTTGAATGTCGTGAA
GGAAGCAGTTCCTCTGGAAGCTTCTTGAAGACAAACAACGTCTGTAGCGACCCTTTGCAGGCAGCGGAACCCCCCACCTGGCGACA
GGTGCCTCTGCGGCCAAAAGCCACGTGTATAAGATACACCTGCAAAGGCGGCACAACCCCAGTGCCACGTTGTGAGTTGGATAGTT
GTGGAAAGAGTCAAATGGCTCTCCTCAAGCGTATTCAACAAGGGGCTGAAGGATGCCCAGAAGGTACCCCATTGTATGGGATCTGA
TCTGGGGCCTCGGTACACATGCTTTACATGTGTTTAGTCGAGGTTAAAAAAACGTCTAGGCCCCCCGAACCACGGGGACGTGGTTT
TCCTTTGAAAAACACGATGATAATATGGCCACAACCATGGTGAGCAAGGGCGAGGAGCTGTTCACCGGGGTGGTGCCCATCCTGGT
CGAGCTGGACGGCGACGTAAACGGCCACAAGTTCAGCGTGTCCGGCGAGGGCGAGGGCGATGCCACCTACGGCAAGCTGACCCTGA
AGTTCATCTGCACCACCGGCAAGCTGCCCGTGCCCTGGCCCACCCTCGTGACCACCCTGACCTACGGCGTGCAGTGCTTCAGCCGC
TACCCCGACCACATGAAGCAGCACGACTTCTTCAAGTCCGCCATGCCCGAAGGCTACGTCCAGGTAAGAAGCAAGGTTTCATTTAG
GGGAAGGGAAATGATTCAGGACGAGAGTCTTTGTGCTGCTGAGTGCCTGTGATGAAGAAGCATGTTAGTCCTGGGCAACGTAGCGA
GACCCCATCTCTACAAAAAATAGAAAAATTAGCCAGGTATAGTGGCGCACACCTGTGATTCCAGCTACGCAGGAGGCTGAGGTGGG
AGGATTGCTTGAGCCCAGGAGGTTGAGGCTGCAGTGAGCTGGAATCATGCCACTACTCCAACCTGGGCAACACAGCAAGGACCCTG
TCTCAAAAGCTACTTACAGAAAAGAATTAG

#### pcDNA3.1(+) Laccase2 Scarless-Split eGFP Delta Repeat

CATTGAGAAATGACTGAGTTCCGGTGCTCTCAAGTCATTGATCTTTGTCGACTTTTATTTGGTCTCTGTAATAACGACTTCAAAAA
CATTAAATTCTGTTGCGAAGCCAGTAAGCTACAAAAAGAAAAAACAAGAGAGAATGCTATAGTCGTATAGTATAGTTTCCCGACTA
TCTGATACCCATTACTTATCTAGGGGGAATGCGAACCCAAAATTTTATCAGTTTTCTCGGATATCGATAGATATTGGGGAATAAAT
TTAAATAAATAAATTTTGGGCGGGTTTAGGGCGTGGCAAAAAGTTTTTTGGCAAATCGCTAGAAATTTACAAGACTTATAAAATTA
TGAAAAAATACAACAAAATTTTAAACACGTGGGCGTGACAGTTTTGGGCGGTTTTAGGGCGTTAGAGTAGGCGAGGACAGGGTTAC
ATCGACTAGGCTTTGATCCTGATCAAGAATATATATACTTTATACCGCTTCCTTCTACATGTTACCTATTTTTCAACGAATCTAGT
ATACCTTTTTACTGTACGATTTATGGGTATAATAATAAGCTAAATCGAGACTAAGTTTTATTGTTATATATATTTTTTTTTATTTTA
TGCAGGAGCGCACCATCTTCTTCAAGGACGACGGCAACTACAAGACCCGCGCCGAGGTGAAGTTCGAGGGCGACACCCTGGTGAAC
CGCATCGAGCTGAAGGGCATCGACTTCAAGGAGGACGGCAACATCCTGGGGCACAAGCTGGAGTACAACCACAACAGCCACAACGT
CTATATCATGGCCGACAAGCAGAAGAACGGCATCAAGGTGAACTTCAAGATCCGCCACAACATCGAGGACGGCAGCGTGCAGCTCG
CCGACCACTACCAGCAGAACACCCCCATCGGCGACGGCCCCGTGCTGCTGCCCGACAACCACTACCTGAGCACCCAGTCCGCCCTG
AGCAAAGACCCCAACGAGAAGCGCGATCACATGGTCCTGCTGGAGTTCGTGACCGCCGCCGGGATCACTCTCGGCATGGACGAGCT
GTACAAGTAAGGATCTGAATTCTGCAGTCGACGATCCGCCCCTCTCCCTCCCCCCCCCCTAACGTTACTGGCCGAAGCCGCTTGGA
ATAAGGCCGGTGTGCGTTTGTCTATATGTTATTTTCCACCATATTGCCGTCTTTTGGCAATGTGAGGGCCCGGAAACCTGGCCCTG
TCTTCTTGACGAGCATTCCTAGGGGTCTTTCCCCTCTCGCCAAAGGAATGCAAGGTCTGTTGAATGTCGTGAAGGAAGCAGTTCCT
CTGGAAGCTTCTTGAAGACAAACAACGTCTGTAGCGACCCTTTGCAGGCAGCGGAACCCCCCACCTGGCGACAGGTGCCTCTGCGG
CCAAAAGCCACGTGTATAAGATACACCTGCAAAGGCGGCACAACCCCAGTGCCACGTTGTGAGTTGGATAGTTGTGGAAAGAGTCA
AATGGCTCTCCTCAAGCGTATTCAACAAGGGGCTGAAGGATGCCCAGAAGGTACCCCATTGTATGGGATCTGATCTGGGGCCTCGG
TACACATGCTTTACATGTGTTTAGTCGAGGTTAAAAAAACGTCTAGGCCCCCCGAACCACGGGGACGTGGTTTTCCTTTGAAAAAC
ACGATGATAATATGGCCACAACCATGGTGAGCAAGGGCGAGGAGCTGTTCACCGGGGTGGTGCCCATCCTGGTCGAGCTGGACGGC
GACGTAAACGGCCACAAGTTCAGCGTGTCCGGCGAGGGCGAGGGCGATGCCACCTACGGCAAGCTGACCCTGAAGTTCATCTGCAC
CACCGGCAAGCTGCCCGTGCCCTGGCCCACCCTCGTGACCACCCTGACCTACGGCGTGCAGTGCTTCAGCCGCTACCCCGACCACA
TGAAGCAGCACGACTTCTTCAAGTCCGCCATGCCCGAAGGCTACGTCCAGGTAAGTATTCAAAATTCCAAAATTTTTTACTAGAAA
TATTCGATTTTTTAATAGGCAGTTTCTATACTATTG

#### pcDNA3.1(+) POLR2A Scarless-Split eGFP Delta Repeat

AGGCACGTAGCAGGCCAGGGCCTCTCTCAGCCACCTGAGCAGAAAGCTTTCCAAGATAGGGCAGGCTGGGTTAGGCCATCTGAGTC
TGTCTCGTTCATTGGGATCCAGACTTGACTGTCTTGTTAAAGGCTGTTGCTGCCCAGGTGTGCAGGGAGCTGTTGGTCTCTGGCAT
TCAGGGTGGGGGTGGTATAAACCCGGGGCAGCTTGCATATGGCAGGGAAGAGGGATCCGTGGAGGAACAGTGCAGAAGGCTTTATG
TTCAGAATCTCTCTTGCTTTTCTTCTAGACTGAGTTCCTTGAGATTGGTGAATGCTGTGTATTATTCATCCCTGATAACCTGGTGT
TTGGCCCAGGGCCTTGTCCAGAGGAGTGTTTGATAAGTGTTTCAAGTGAATTAGCACCACGATGTCATCTCTTTTCAGTTTACAAA
GGACGGACACCCCCTGACCCGGTCTCAGAAAGCCTGAAAGCAGAATTAGTCATTAGAAGGGTGGTTGGCTTGGTCGGCATAGACTT
TGAGCAGAAAGAGGTTGAAAATGTTGAGCCTGATTTCTCTTAGGCCCCTCTGCAGTGTCTGTTGTGGAGGCCAGATACGTAACTGC
TTCCGCTTTTTTTGGTGTCATTCAAGGTGAGCAAATCCCCTTCATGTTTCTCACCAGACAATGCAGCTGATGAGGTTCCAGCTTTG
CAAATGTAGTCATCCATGAGGACTGTCTTCCTGAGATTTCATCAGGCTCGAGTGGACTTGCAAAGGACTTTAGGTCCATTGTCCTT
TTATTCTTAGATACCTCTTTCACTGAGACCTTTTCCTTACCTCACCTCTCTAGGAGCGCACCATCTTCTTCAAGGACGACGGCAAC
TACAAGACCCGCGCCGAGGTGAAGTTCGAGGGCGACACCCTGGTGAACCGCATCGAGCTGAAGGGCATCGACTTCAAGGAGGACGG
CAACATCCTGGGGCACAAGCTGGAGTACAACCACAACAGCCACAACGTCTATATCATGGCCGACAAGCAGAAGAACGGCATCAAGG
TGAACTTCAAGATCCGCCACAACATCGAGGACGGCAGCGTGCAGCTCGCCGACCACTACCAGCAGAACACCCCCATCGGCGACGGC
CCCGTGCTGCTGCCCGACAACCACTACCTGAGCACCCAGTCCGCCCTGAGCAAAGACCCCAACGAGAAGCGCGATCACATGGTCCT
GCTGGAGTTCGTGACCGCCGCCGGGATCACTCTCGGCATGGACGAGCTGTACAAGTAAGGATCTGAATTCTGCAGTCGACGATCCG
CCCCTCTCCCTCCCCCCCCCCTAACGTTACTGGCCGAAGCCGCTTGGAATAAGGCCGGTGTGCGTTTGTCTATATGTTATTTTCCA
CCATATTGCCGTCTTTTGGCAATGTGAGGGCCCGGAAACCTGGCCCTGTCTTCTTGACGAGCATTCCTAGGGGTCTTTCCCCTCTC
GCCAAAGGAATGCAAGGTCTGTTGAATGTCGTGAAGGAAGCAGTTCCTCTGGAAGCTTCTTGAAGACAAACAACGTCTGTAGCGAC
CCTTTGCAGGCAGCGGAACCCCCCACCTGGCGACAGGTGCCTCTGCGGCCAAAAGCCACGTGTATAAGATACACCTGCAAAGGCGG
CACAACCCCAGTGCCACGTTGTGAGTTGGATAGTTGTGGAAAGAGTCAAATGGCTCTCCTCAAGCGTATTCAACAAGGGGCTGAAG
GATGCCCAGAAGGTACCCCATTGTATGGGATCTGATCTGGGGCCTCGGTACACATGCTTTACATGTGTTTAGTCGAGGTTAAAAAA
ACGTCTAGGCCCCCCGAACCACGGGGACGTGGTTTTCCTTTGAAAAACACGATGATAATATGGCCACAACCATGGTGAGCAAGGGC
GAGGAGCTGTTCACCGGGGTGGTGCCCATCCTGGTCGAGCTGGACGGCGACGTAAACGGCCACAAGTTCAGCGTGTCCGGCGAGGG
CGAGGGCGATGCCACCTACGGCAAGCTGACCCTGAAGTTCATCTGCACCACCGGCAAGCTGCCCGTGCCCTGGCCCACCCTCGTGA
CCACCCTGACCTACGGCGTGCAGTGCTTCAGCCGCTACCCCGACCACATGAAGCAGCACGACTTCTTCAAGTCCGCCATGCCCGAA
GGCTACGTCCAGGTGTGTGGTCCAAATGGAAACCTGGCTTAAGTGGGCAGTGGGGCTCTGGGGTGCAAGGTGGAGGCTAGAGAGGA
AGAGCTGTGTTTTTTTTCCTGACTTACCCAGCAGTGGTCTGTGAGATTGTCTTTTCTGGTGGGCGAACAAAAAGGGGGTTAGGAAA
ACTCAGGCCAAAAAAGTGTAAGGCGTTAATTCCCCATTTAATTCCTTAAAATTTCATGTAATACCAGGTATTGCCTGTAAAGGAAA
GATAAAGGGAAAAATAAGTAAGACCTTGTTAAAATTTTATTTTTCTATTTTAACCTTCACTTATTTCCTAATTATTAAAAGAAATT
TATGCTTATTGTTAAGAACAAAAAAATTTCAGTATTACAATGAATTTTTAATTAAAAGTTTTTG

#### pcDNA3.1(+) ciRS-7 Scarless-Split eGFP Delta Repeat

CTGTAAGAGTAGTCTCATGATGTCTTTTAATTTAATATCTAGGCCCTAATACACAACTCTTTGAATATTGCTACTATCTTTCAGCA
TGACTTTCCAATACCATTAAGATTAAGCCTTTCTGTTCTTCAGTTTCTCTAACTCAAGTCTTCTAAGCTCCACAGCTTCCAACATC
TTTAATTACCAGATCTTCCCAATATTATAACATCCGTGCATTCCAGTCCCTGTCCTGCAGCATCTCCAATGCCCTTGTTTTCTAAA
CTTCCCAGTATTCAAGTCATTAAAAATTTAGGTCTTCTAATATCTCCAACAACCCATCTTCAAAGTCCAGGTCTTGTCTTTCAGAA
TCCAGAGTTCTAAATTACAAGTCTTCCATGGTCCTAATCCTCCAATGTCTCCTATATACAGGTCTTCTAGAGGCTATGTATTTCAT
AGTGTTTAAGTCTAGATTCTTCAATGTCTTCCAGCACTTACATCATTCAGTTTTTACATTTTTAAATGTCTCCTACTTTCTAGTTT
TACAGTGTATTTTTATCGCCAATTCCTCCAATGCCCAAGTCTTTAAATACCTGCAAGAACTTTCAGGTACATCAATATCCTCAATG
TCTGAATGAATAGACCTCTTAATATCCTGGTCTCATATTGTCCAGGCTGCCAGTATTCTTGACATCCAGCATCTTTAAAATTTAAT
TCTTCCAAACTGTCTTTTTGACATAAATATGTCAATATCCATGCTTACCAATGTCTCCCAATATCCAGGGCTTCTCATTTTAGGAT
TTATAAATGTCTACATCTGCTGTTGTTTATATCTGCCAGTATTTAGGAGTTCCAAGGTACAGGACTTACAATATCTCTAATTTCCA
GAGATTTCAGTGTCTATGACTTCCAATATTCAATATCCCAATGTTCAGGGCCTCCAGAAATATCAATGTCCAGAGTAACCAATGTC
TTCAAAATCCAAGGCTTGCAATGTCTCCAATATCCTGTGCTTACAATGTTTCCAATGTCCAGGAGCGCACCATCTTCTTCAAGGAC
GACGGCAACTACAAGACCCGCGCCGAGGTGAAGTTCGAGGGCGACACCCTGGTGAACCGCATCGAGCTGAAGGGCATCGACTTCAA
GGAGGACGGCAACATCCTGGGGCACAAGCTGGAGTACAACCACAACAGCCACAACGTCTATATCATGGCCGACAAGCAGAAGAACG
GCATCAAGGTGAACTTCAAGATCCGCCACAACATCGAGGACGGCAGCGTGCAGCTCGCCGACCACTACCAGCAGAACACCCCCATC
GGCGACGGCCCCGTGCTGCTGCCCGACAACCACTACCTGAGCACCCAGTCCGCCCTGAGCAAAGACCCCAACGAGAAGCGCGATCA
CATGGTCCTGCTGGAGTTCGTGACCGCCGCCGGGATCACTCTCGGCATGGACGAGCTGTACAAGTAAGGATCTGAATTCTGCAGTC
GACGATCCGCCCCTCTCCCTCCCCCCCCCCTAACGTTACTGGCCGAAGCCGCTTGGAATAAGGCCGGTGTGCGTTTGTCTATATGT
TATTTTCCACCATATTGCCGTCTTTTGGCAATGTGAGGGCCCGGAAACCTGGCCCTGTCTTCTTGACGAGCATTCCTAGGGGTCTT
TCCCCTCTCGCCAAAGGAATGCAAGGTCTGTTGAATGTCGTGAAGGAAGCAGTTCCTCTGGAAGCTTCTTGAAGACAAACAACGTC
TGTAGCGACCCTTTGCAGGCAGCGGAACCCCCCACCTGGCGACAGGTGCCTCTGCGGCCAAAAGCCACGTGTATAAGATACACCTG
CAAAGGCGGCACAACCCCAGTGCCACGTTGTGAGTTGGATAGTTGTGGAAAGAGTCAAATGGCTCTCCTCAAGCGTATTCAACAAG
GGGCTGAAGGATGCCCAGAAGGTACCCCATTGTATGGGATCTGATCTGGGGCCTCGGTACACATGCTTTACATGTGTTTAGTCGAG
GTTAAAAAAACGTCTAGGCCCCCCGAACCACGGGGACGTGGTTTTCCTTTGAAAAACACGATGATAATATGGCCACAACCATGGTG
AGCAAGGGCGAGGAGCTGTTCACCGGGGTGGTGCCCATCCTGGTCGAGCTGGACGGCGACGTAAACGGCCACAAGTTCAGCGTGTC
CGGCGAGGGCGAGGGCGATGCCACCTACGGCAAGCTGACCCTGAAGTTCATCTGCACCACCGGCAAGCTGCCCGTGCCCTGGCCCA
CCCTCGTGACCACCCTGACCTACGGCGTGCAGTGCTTCAGCCGCTACCCCGACCACATGAAGCAGCACGACTTCTTCAAGTCCGCC
ATGCCCGAAGGCTACGTCCAGGTATTCTAACATGTTCTATAGTCCAGATCTTCCAGCCTTTCCCCAGGTCCAGGCCTTTTCAAATC
TAGGCTTCCCCTGTCTGAGCCTTCCCATGTTAAAATCACTCTACAGAAAAATCTTCCAACTCAACTATTTCTGCTTCTAGTTGAAT

TAGCTTCAAAAACATTTAATGGGACATTCCTCCTTATAGACCAGATCTTATGAACCAACAGTACTGCAGTTTGGCGGCCGCTCGAC
TGGTTACTCTGGA

**pcDNA3.1(+) IGF2BP1 Scarless-Split eGFP Delta Repeat**

GACTGAACATGGAGGAATTGAGGTTGGGTATTTCCCCTGAGGTAGGAAAAAGGCTGGGTCAGTTTCCCGTTAGCCGTCAAGTCCTC
ATCACATCTTTAAGCCTTCCATGCAGGATAAAGGGCTGCAGAGCTATTTTCAAATTGACATCAAACTGGATTTCTGTTGACTTCGT
CTTCCCTTTTTAAGGTCCACAGAAGAAGATGGGAAGGAAAGAAGTCTGAGGGCATCTTATTTGCACTCCGCTGTCATTTCTAAGGA
AGGCCTTTAATGCCAAATTCTCATCTTTTATGTCCCCACTAAATCCTAAGGTTCTTGAACTTCTGATCAGACAGCCAAAAAATGAA
CCATCAACTAGCTTAACCTAACATATGTGAGGATAGAGGACTGGGACAGCTCTCTGGGCCACTGGAGAGTCAGACAGGCCTGCCCT
CTGTGTGACTTGACCGCGGTCTCTTTCTTCCAGGAGCGCACCATCTTCTTCAAGGACGACGGCAACTACAAGACCCGCGCCGAGGT
GAAGTTCGAGGGCGACACCCTGGTGAACCGCATCGAGCTGAAGGGCATCGACTTCAAGGAGGACGGCAACATCCTGGGGCACAAGC
TGGAGTACAACCACAACAGCCACAACGTCTATATCATGGCCGACAAGCAGAAGAACGGCATCAAGGTGAACTTCAAGATCCGCCAC
AACATCGAGGACGGCAGCGTGCAGCTCGCCGACCACTACCAGCAGAACACCCCCATCGGCGACGGCCCCGTGCTGCTGCCCGACAA
CCACTACCTGAGCACCCAGTCCGCCCTGAGCAAAGACCCCAACGAGAAGCGCGATCACATGGTCCTGCTGGAGTTCGTGACCGCCG
CCGGGATCACTCTCGGCATGGACGAGCTGTACAAGTAAGGATCTGAATTCTGCAGTCGACGATCCGCCCCTCTCCCTCCCCCCCCC
CTAACGTTACTGGCCGAAGCCGCTTGGAATAAGGCCGGTGTGCGTTTGTCTATATGTTATTTTCCACCATATTGCCGTCTTTTGGC
AATGTGAGGGCCCGGAAACCTGGCCCTGTCTTCTTGACGAGCATTCCTAGGGGTCTTTCCCCTCTCGCCAAAGGAATGCAAGGTCT
GTTGAATGTCGTGAAGGAAGCAGTTCCTCTGGAAGCTTCTTGAAGACAAACAACGTCTGTAGCGACCCTTTGCAGGCAGCGGAACC
CCCCACCTGGCGACAGGTGCCTCTGCGGCCAAAAGCCACGTGTATAAGATACACCTGCAAAGGCGGCACAACCCCAGTGCCACGTT
GTGAGTTGGATAGTTGTGGAAAGAGTCAAATGGCTCTCCTCAAGCGTATTCAACAAGGGGCTGAAGGATGCCCAGAAGGTACCCCA
TTGTATGGGATCTGATCTGGGGCCTCGGTACACATGCTTTACATGTGTTTAGTCGAGGTTAAAAAAACGTCTAGGCCCCCCGAACC
ACGGGGACGTGGTTTTCCTTTGAAAAACACGATGATAATATGGCCACAACCATGGTGAGCAAGGGCGAGGAGCTGTTCACCGGGGT
GGTGCCCATCCTGGTCGAGCTGGACGGCGACGTAAACGGCCACAAGTTCAGCGTGTCCGGCGAGGGCGAGGGCGATGCCACCTACG
GCAAGCTGACCCTGAAGTTCATCTGCACCACCGGCAAGCTGCCCGTGCCCTGGCCCACCCTCGTGACCACCCTGACCTACGGCGTG
CAGTGCTTCAGCCGCTACCCCGACCACATGAAGCAGCACGACTTCTTCAAGTCCGCCATGCCCGAAGGCTACGTCCAGGTAAGTCT
CGACGGTACCGCGGGCCCGGGATCC

To generate the following expression plasmids with altered splice site sequences, the following sequences were cloned between the EcoRI and NotI restriction sites in **pcDNA3.1(+) CircRNA Mini Vector** (Addgene #60648). The split eGFP ORF is highlighted. Altered splice site sequences are underlined.

**pcDNA3.1(+) ciRS-7 Scarless-Split eGFP (Laccase2 ss)**

CTGTAAGAGTAGTCTCATGATGTCTTTTAATTTAATATCTAGGCCCTAATACACAACTCTTTGAATATTGCTACTATCTTTCAGCA
TGACTTTCCAATACCATTAAGATTAAGCCTTTCTGTTCTTCAGTTTCTCTAACTCAAGTCTTCTAAGCTCCACAGCTTCCAACATC
TTTAATTACCAGATCTTCCCAATATTATAACATCCGTGCATTCCAGTCCCTGTCCTGCAGCATCTCCAATGCCCTTGTTTTCTAAA
CTTCCCAGTATTCAAGTCATTAAAAATTTAGGTCTTCTAATATCTCCAACAACCCATCTTCAAAGTCCAGGTCTTGTCTTTCAGAA
TCCAGAGTTCTAAATTACAAGTCTTCCATGGTCCTAATCCTCCAATGTCTCCTATATACAGGTCTTCTAGAGGCTATGTATTTCAT
AGTGTTTAAGTCTAGATTCTTCAATGTCTTCCAGCACTTACATCATTCAGTTTTTACATTTTTAAATGTCTCCTACTTTCTAGTTT
TACAGTGTATTTTTATCGCCAATTCCTCCAATGCCCAAGTCTTTAAATACCTGCAAGAACTTTCAGGTACATCAATATCCTCAATG
TCTGAATGAATAGACCTCTTAATATCCTGGTCTCATATTGTCCAGGCTGCCAGTATTCTTGACATCCAGCATCTTTAAAATTTAAT
TCTTCCAAACTGTCTTTTTGACATAAATATGTCAATATCCATGCTTACCAATGTCTCCCAATATCCAGGGCTTCTCATTTTAGGAT
TTATAAATGTCTACATCTGCTGTTGTTTATATCTGCCAGTATTTAGGAGTTCCAAGGTACAGGACTTACAATATCTCTAATTTCCA
GAGATTTCAGTGTCTATGACTTCCAATATTCAATATCCCAATGTTCAGGGCCTCCAGAAATATCAATGTCCAGAGTAACCAATGTC
TTCAAAATCCAAGGCTTGCAATGTCTCCAATATCCTGTGCTTATTTTTTTTTATTTTATGCAGGAGCGCACCATCTTCTTCAAGGAC
GACGGCAACTACAAGACCCGCGCCGAGGTGAAGTTCGAGGGCGACACCCTGGTGAACCGCATCGAGCTGAAGGGCATCGACTTCAA
GGAGGACGGCAACATCCTGGGGCACAAGCTGGAGTACAACCACAACAGCCACAACGTCTATATCATGGCCGACAAGCAGAAGAACG
GCATCAAGGTGAACTTCAAGATCCGCCACAACATCGAGGACGGCAGCGTGCAGCTCGCCGACCACTACCAGCAGAACACCCCCATC
GGCGACGGCCCCGTGCTGCTGCCCGACAACCACTACCTGAGCACCCAGTCCGCCCTGAGCAAAGACCCCAACGAGAAGCGCGATCA
CATGGTCCTGCTGGAGTTCGTGACCGCCGCCGGGATCACTCTCGGCATGGACGAGCTGTACAAGTAAGGATCTGAATTCTGCAGTC
GACGATCCGCCCCTCTCCCTCCCCCCCCCCTAACGTTACTGGCCGAAGCCGCTTGGAATAAGGCCGGTGTGCGTTTGTCTATATGT
TATTTTCCACCATATTGCCGTCTTTTGGCAATGTGAGGGCCCGGAAACCTGGCCCTGTCTTCTTGACGAGCATTCCTAGGGGTCTT
TCCCCTCTCGCCAAAGGAATGCAAGGTCTGTTGAATGTCGTGAAGGAAGCAGTTCCTCTGGAAGCTTCTTGAAGACAAACAACGTC
TGTAGCGACCCTTTGCAGGCAGCGGAACCCCCCACCTGGCGACAGGTGCCTCTGCGGCCAAAAGCCACGTGTATAAGATACACCTG
CAAAGGCGGCACAACCCCAGTGCCACGTTGTGAGTTGGATAGTTGTGGAAAGAGTCAAATGGCTCTCCTCAAGCGTATTCAACAAG
GGGCTGAAGGATGCCCAGAAGGTACCCCATTGTATGGGATCTGATCTGGGGCCTCGGTACACATGCTTTACATGTGTTTAGTCGAG
GTTAAAAAAACGTCTAGGCCCCCCGAACCACGGGGACGTGGTTTTCCTTTGAAAAACACGATGATAATATGGCCACAACCATGGTG
AGCAAGGGCGAGGAGCTGTTCACCGGGGTGGTGCCCATCCTGGTCGAGCTGGACGGCGACGTAAACGGCCACAAGTTCAGCGTGTC

CGGCGAGGGCGAGGGCGATGCCACCTACGGCAAGCTGACCCTGAAGTTCATCTGCACCACCGGCAAGCTGCCCGTGCCCTGGCCCA
CCCTCGTGACCACCCTGACCTACGGCGTGCAGTGCTTCAGCCGCTACCCCGACCACATGAAGCAGCACGACTTCTTCAAGTCCGCC
ATGCCCGAAGGCTACGTCCAGGTAAGTTAACATGTTCTATAGTCCAGATCTTCCAGCCTTTCCCCAGGTCCAGGCCTTTTCAAATC
TAGGCTTCCCCTGTCTGAGCCTTCCCATGTTAAAATCACTCTACAGAAAAATCTTCCAACTCAACTATTTCTGCTTCTAGTTGAAT
TAGCTTCAAAAACATTTAATGGGACATTCCTCCTTATAGACCAGATCTTATGAACCAACAGTACTGCAGTTTGGCGGCCGCTCGAC
TGGTTACTCTGGACATTGATATTTCTGGAGGCCCTGAACATTGGGATATTGAATATTGGAAGTCATAGACACTGAAATCTCTGGAA
ATTAGAGATATTGTAAGTCCTGTACCTTGGAACTCCTAAATACTGGCAGATATAAACAACAGCAGATGTAGACATTTATAAATCCT
AAAATGAGAAGCCCTGGATATTGGGAGACATTGGTAAGCATGGATATTGACATATTTATGTCAAAAAGACAGTTTGGAAGAATTAA
ATTTTAAAGATGCTGGATGTCAAGAATACTGGCAGCCTGGACAATATGAGACCAGGATATTAAGAGGTCTATTCATTCAGACATTG
AGGATATTGATGTACCTGAAAGTTCTTGCAGGTATTTAAAGACTTGGGCATTGGAGGAATTGGCGATAAAAATACACTGTAAAACT
AGAAAGTAGGAGACATTTAAAAATGTAAAAACTGAATGATGTAAGTGCTGGAAGACATTGAAGAATCTAGACTTAAACACTATGAA
ATACATAGCCTCTAGAAGACCTGTATATAGGAGACATTGGAGGATTAGGACCATGGAAGACTTGTAATTTAGAACTCTGGATTCTG
AAAGACAAGACCTGGACTTTGAAGATGGGTTGTTGGAGATATTAGAAGACCTAAATTTTTAATGACTTGAATACTGGGAAGTTTAG
AAAACAAGGGCATTGGAGATGCTGCAGGACAGGGACTGGAATGCACGGATGTTATAATATTGGGAAGATCTGGTAATTAAAGATGT
TGGAAGCTGTGGAGCTTAGAAGACTTGAGTTAGAGAAACTGAAGAACAGAAAGGCTTAATCTTAATGGTATTGGAAAGTCATGCTG
AAAGATAGTAGCAATATTCAAAGAGTTGTGTATTAGGGCCTAGATATTAAATTAAAAGACATCATGAGACTACTCTTACAGGTCGA
GCATGCATCTAGA

##### pcDNA3.1(+) ciRS-7 Scarless-Split eGFP (ZKSCAN1 ss)

CTGTAAGAGTAGTCTCATGATGTCTTTTAATTTAATATCTAGGCCCTAATACACAACTCTTTGAATATTGCTACTATCTTTCAGCA
TGACTTTCCAATACCATTAAGATTAAGCCTTTCTGTTCTTCAGTTTCTCTAACTCAAGTCTTCTAAGCTCCACAGCTTCCAACATC
TTTAATTACCAGATCTTCCCAATATTATAACATCCGTGCATTCCAGTCCCTGTCCTGCAGCATCTCCAATGCCCTTGTTTTCTAAA
CTTCCCAGTATTCAAGTCATTAAAAATTTAGGTCTTCTAATATCTCCAACAACCCATCTTCAAAGTCCAGGTCTTGTCTTTCAGAA
TCCAGAGTTCTAAATTACAAGTCTTCCATGGTCCTAATCCTCCAATGTCTCCTATATACAGGTCTTCTAGAGGCTATGTATTTCAT
AGTGTTTAAGTCTAGATTCTTCAATGTCTTCCAGCACTTACATCATTCAGTTTTTACATTTTTAAATGTCTCCTACTTTCTAGTTT
TACAGTGTATTTTTATCGCCAATTCCTCCAATGCCCAAGTCTTTAAATACCTGCAAGAACTTTCAGGTACATCAATATCCTCAATG
TCTGAATGAATAGACCTCTTAATATCCTGGTCTCATATTGTCCAGGCTGCCAGTATTCTTGACATCCAGCATCTTTAAAATTTAAT
TCTTCCAAACTGTCTTTTTGACATAAATATGTCAATATCCATGCTTACCAATGTCTCCCAATATCCAGGGCTTCTCATTTTAGGAT
TTATAAATGTCTACATCTGCTGTTGTTTATATCTGCCAGTATTTAGGAGTTCCAAGGTACAGGACTTACAATATCTCTAATTTCCA
GAGATTTCAGTGTCTATGACTTCCAATATTCAATATCCCAATGTTCAGGGCCTCCAGAAATATCAATGTCCAGAGTAACCAATGTC
TTCAAAATCCAAGGCTTGCAATGTCTCCAATATCCTGTGCTTACTTTTTTTTTATACTTCAGGAGCGCACCATCTTCTTCAAGGAC
GACGGCAACTACAAGACCCGCGCCGAGGTGAAGTTCGAGGGCGACACCCTGGTGAACCGCATCGAGCTGAAGGGCATCGACTTCAA
GGAGGACGGCAACATCCTGGGGCACAAGCTGGAGTACAACCACAACAGCCACAACGTCTATATCATGGCCGACAAGCAGAAGAACG
GCATCAAGGTGAACTTCAAGATCCGCCACAACATCGAGGACGGCAGCGTGCAGCTCGCCGACCACTACCAGCAGAACACCCCCATC
GGCGACGGCCCCGTGCTGCTGCCCGACAACCACTACCTGAGCACCCAGTCCGCCCTGAGCAAAGACCCCAACGAGAAGCGCGATCA
CATGGTCCTGCTGGAGTTCGTGACCGCCGCCGGGATCACTCTCGGCATGGACGAGCTGTACAAGTAAGGATCTGAATTCTGCAGTC
GACGATCCGCCCCTCTCCCTCCCCCCCCCCTAACGTTACTGGCCGAAGCCGCTTGGAATAAGGCCGGTGTGCGTTTGTCTATATGT
TATTTTCCACCATATTGCCGTCTTTTGGCAATGTGAGGGCCCGGAAACCTGGCCCTGTCTTCTTGACGAGCATTCCTAGGGGTCTT
TCCCCTCTCGCCAAAGGAATGCAAGGTCTGTTGAATGTCGTGAAGGAAGCAGTTCCTCTGGAAGCTTCTTGAAGACAAACAACGTC
TGTAGCGACCCTTTGCAGGCAGCGGAACCCCCCACCTGGCGACAGGTGCCTCTGCGGCCAAAAGCCACGTGTATAAGATACACCTG
CAAAGGCGGCACAACCCCAGTGCCACGTTGTGAGTTGGATAGTTGTGGAAAGAGTCAAATGGCTCTCCTCAAGCGTATTCAACAAG
GGGCTGAAGGATGCCCAGAAGGTACCCCATTGTATGGGATCTGATCTGGGGCCTCGGTACACATGCTTTACATGTGTTTAGTCGAG
GTTAAAAAAACGTCTAGGCCCCCCGAACCACGGGGACGTGGTTTTCCTTTGAAAAACACGATGATAATATGGCCACAACCATGGTG
AGCAAGGGCGAGGAGCTGTTCACCGGGGTGGTGCCCATCCTGGTCGAGCTGGACGGCGACGTAAACGGCCACAAGTTCAGCGTGTC
CGGCGAGGGCGAGGGCGATGCCACCTACGGCAAGCTGACCCTGAAGTTCATCTGCACCACCGGCAAGCTGCCCGTGCCCTGGCCCA
CCCTCGTGACCACCCTGACCTACGGCGTGCAGTGCTTCAGCCGCTACCCCGACCACATGAAGCAGCACGACTTCTTCAAGTCCGCC
ATGCCCGAAGGCTACGTCCAGGTAAGATAACATGTTCTATAGTCCAGATCTTCCAGCCTTTCCCCAGGTCCAGGCCTTTTCAAATC
TAGGCTTCCCCTGTCTGAGCCTTCCCATGTTAAAATCACTCTACAGAAAAATCTTCCAACTCAACTATTTCTGCTTCTAGTTGAAT
TAGCTTCAAAAACATTTAATGGGACATTCCTCCTTATAGACCAGATCTTATGAACCAACAGTACTGCAGTTTGGCGGCCGCTCGAC
TGGTTACTCTGGACATTGATATTTCTGGAGGCCCTGAACATTGGGATATTGAATATTGGAAGTCATAGACACTGAAATCTCTGGAA
ATTAGAGATATTGTAAGTCCTGTACCTTGGAACTCCTAAATACTGGCAGATATAAACAACAGCAGATGTAGACATTTATAAATCCT
AAAATGAGAAGCCCTGGATATTGGGAGACATTGGTAAGCATGGATATTGACATATTTATGTCAAAAAGACAGTTTGGAAGAATTAA
ATTTTAAAGATGCTGGATGTCAAGAATACTGGCAGCCTGGACAATATGAGACCAGGATATTAAGAGGTCTATTCATTCAGACATTG
AGGATATTGATGTACCTGAAAGTTCTTGCAGGTATTTAAAGACTTGGGCATTGGAGGAATTGGCGATAAAAATACACTGTAAAACT
AGAAAGTAGGAGACATTTAAAAATGTAAAAACTGAATGATGTAAGTGCTGGAAGACATTGAAGAATCTAGACTTAAACACTATGAA
ATACATAGCCTCTAGAAGACCTGTATATAGGAGACATTGGAGGATTAGGACCATGGAAGACTTGTAATTTAGAACTCTGGATTCTG
AAAGACAAGACCTGGACTTTGAAGATGGGTTGTTGGAGATATTAGAAGACCTAAATTTTTAATGACTTGAATACTGGGAAGTTTAG
AAAACAAGGGCATTGGAGATGCTGCAGGACAGGGACTGGAATGCACGGATGTTATAATATTGGGAAGATCTGGTAATTAAAGATGT
TGGAAGCTGTGGAGCTTAGAAGACTTGAGTTAGAGAAACTGAAGAACAGAAAGGCTTAATCTTAATGGTATTGGAAAGTCATGCTG
AAAGATAGTAGCAATATTCAAAGAGTTGTGTATTAGGGCCTAGATATTAAATTAAAAGACATCATGAGACTACTCTTACAGGTCGA
GCATGCATCTAGA

To generate **pcDNA3.1(+) IGF2BP1 Scarless-Split eGFP (Upstream miR-92a)**, a miR-92a target sequence (ACAGGCCGGGACAAGTGCAATA) was inserted immediately upstream of the NheI restriction site in pcDNA3.1(+) IGF2BP1 Scarless-Split eGFP.

To generate **pcDNA3.1(+) IGF2BP1 Scarless-Split eGFP (Upstream miR-92a Mut)**, a miR-92a seed mutant target sequence (ACAGGCCGGGATGGTAATCTA) was inserted immediately upstream of the NheI restriction site in pcDNA3.1(+) IGF2BP1 Scarless-Split eGFP.

To generate **pcDNA3.1(+) IGF2BP1 Scarless-Split eGFP (Downstream miR-92a)**, the following sequence with a miR-92a target sequence (ACAGGCCGGGACAAGTGCAATA) was inserted between NotI and BssHII in pcDNA3.1(+) IGF2BP1 Scarless-Split eGFP:

```
TCGAGTCTAGAGGGCCCGTTTAAACCCGCTGATCAGCCTCGAGGCGCGACAGGCCGGGACAAGTGCAATACGCGCCctgtgccttc
tagttgccagccatctgttgtttgcccctcccccgtgccttccttgaccctggaaggtgccactcccactgtcctttcctaataaa
atgaggaaattgcatcgcattgtctgagtaggtgtcattctattctggggggtggggtggggcaggacagcaagggggaggattgg
gaagacaatagcaggcatgctggggatgcggtgggctctatggcttctgaggcggaaagaaccagctggggctctagggggtatcc
ccacgcgccctgtagcggcgcattaagcgcggcgggtgtggtggttacgcgcagcgtgaccgctacacttgccagcgccctagcgc
ccgctcctttcgctttcttcccttcctttctcgccacgttcgccggctttccccgtcaagctctaaatcgggggctccctttaggg
ttccgatttagtgctttacggcacctcgaccccaaaaaacttgattagggtgatggttcacgtagtgggccatcgccctgatagac
ggtttttcgccctttgacgttggagtccacgttctttaatagtggactcttgttccaaactggaacaacactcaaccctatctcgg
tctattcttttgatttataagggattttgccgatttcggcctattggttaaaaaatgagctgatttaacaaaaatttaacgcgaat
taattctgtggaatgtgtgtcagttagggtgtggaaagtccccaggctccccagcaggcagaagtatgcaaagcatgcatctcaat
tagtcagcaaccaggtgtggaaagtccccaggctccccagcaggcagaagtatgcaaagcatgcatctcaattagtcagcaaccat
agtcccgcccctaactccgcccatcccgcccctaactccgcccagttccgcccattctccgccccatggctgactaatttttttta
tttatgcagaggccgaggccgcctctgcctctgagctattccagaagtagtgaggaggcttttttggaggcctaggcttttgcaaa
aagctcccgggagcttgtatatccattttcggatctgatcaagagacaggatgaggatcgtttcgcatgattgaacaagatggatt
gcacgcaggttctccggccgcttgggtggagaggctattcggctatgactgggcacaacagacaatcggctgctctgatgccgccg
tgttccggctgtcagcgcaggggcgcccggttctttttgtcaagaccgacctgtccggtgccctgaatgaactgcaggacgaggca
gcgcggctatcgtggctggccacgacgggcgttccttgcgcagctgtgctcgacgttgtcactgaagcgggaagggactggctgct
attgggcgaagtgccggggcaggatctcctgtcatctcaccttgctcctgccgagaaagtatccatcatggctgatgcaatgcggc
ggctgcatacgcttgatccggctacctgcccattcgaccaccaagcgaaacatcgcatcgagcgagcacgtactcggatggaagcc
ggtcttgtcgatcaggatgatctggacgaagagcatcaggggctcgcgccagccgaactgttcgccaggctcaag
```

To generate **pcDNA3.1(+) IGF2BP1 Scarless-Split eGFP (Downstream miR-92a Mut)**, the following sequence with a miR-92a seed mutant target sequence (ACAGGCCGGGATGGTAATCTA) was inserted between NotI and BssHII in **pcDNA3.1(+) IGF2BP1 Scarless-Split eGFP**:

```
TCGAGTCTAGAGGGCCCGTTTAAACCCGCTGATCAGCCTCGAGGCGCGACAGGCCGGGATGGTAATCTACGCGCCctgtgccttct
agttgccagccatctgttgtttgcccctcccccgtgccttccttgaccctggaaggtgccactcccactgtcctttcctaataaaa
tgaggaaattgcatcgcattgtctgagtaggtgtcattctattctggggggtggggtggggcaggacagcaagggggaggattggg
aagacaatagcaggcatgctggggatgcggtgggctctatggcttctgaggcggaaagaaccagctggggctctagggggtatccc
cacgcgccctgtagcggcgcattaagcgcggcgggtgtggtggttacgcgcagcgtgaccgctacacttgccagcgccctagcgcc
cgctcctttcgctttcttcccttcctttctcgccacgttcgccggctttccccgtcaagctctaaatcgggggctccctttagggt
tccgatttagtgctttacggcacctcgaccccaaaaaacttgattagggtgatggttcacgtagtgggccatcgccctgatagacg
gtttttcgccctttgacgttggagtccacgttctttaatagtggactcttgttccaaactggaacaacactcaaccctatctcggt
ctattcttttgatttataagggattttgccgatttcggcctattggttaaaaaatgagctgatttaacaaaaatttaacgcgaatt
aattctgtggaatgtgtgtcagttagggtgtggaaagtccccaggctccccagcaggcagaagtatgcaaagcatgcatctcaatt
agtcagcaaccaggtgtggaaagtccccaggctccccagcaggcagaagtatgcaaagcatgcatctcaattagtcagcaaccata
gtcccgcccctaactccgcccatcccgcccctaactccgcccagttccgcccattctccgccccatggctgactaattttttttat
ttatgcagaggccgaggccgcctctgcctctgagctattccagaagtagtgaggaggcttttttggaggcctaggcttttgcaaaa
agctcccgggagcttgtatatccattttcggatctgatcaagagacaggatgaggatcgtttcgcatgattgaacaagatggattg
cacgcaggttctccggccgcttgggtggagaggctattcggctatgactgggcacaacagacaatcggctgctctgatgccgccgt
gttccggctgtcagcgcaggggcgcccggttctttttgtcaagaccgacctgtccggtgccctgaatgaactgcaggacgaggcag
```

cgcggctatcgtggctggccacgacgggcgttccttgcgcagctgtgctcgacgttgtcactgaagcgggaagggactggctgcta
ttgggcgaagtgccggggcaggatctcctgtcatctcaccttgctcctgccgagaaagtatccatcatggctgatgcaatgcggcg
gctgcatacgcttgatccggctacctgcccattcgaccaccaagcgaaacatcgcatcgagcgagcacgtactcggatggaagccg
gtcttgtcgatcaggatgatctggacgaagagcatcaggggctcgcgccagccgaactgttcgccaggctcaag

To generate **pcDNA3.1(+) IGF2BP1 Scarless-Split eGFP (Up & Downstream miR-92a)**, a miR-92a target sequence (ACAGGCCGGGACAAGTGCAATA) was inserted immediately upstream of the NheI restriction site in pcDNA3.1(+) IGF2BP1 Scarless-Split eGFP (Downstream miR-92a).

To generate **pcDNA3.1(+) IGF2BP1 Scarless-Split eGFP (Up & Downstream miR-92a Mut)**, a miR-92a seed mutant target sequence (ACAGGCCGGGATGGTAATCTA) was inserted immediately upstream of the NheI restriction site in pcDNA3.1(+) IGF2BP1 Scarless-Split eGFP (Downstream miR-92a Mut).

To generate the following plasmids with the bGH poly(A) signal replaced with wildtype mascRNA:

**pcDNA3.1(+) ZKSCAN1 Scarless-Split eGFP (mascRNA WT)**
**pcDNA3.1(+) Laccase2 Scarless-Split eGFP (mascRNA WT)**
**pcDNA3.1(+) POLR2A Scarless-Split eGFP (mascRNA WT)**
**pcDNA3.1(+) IGF2BP1 Scarless-Split eGFP (mascRNA WT)**

the following sequence was cloned between the XbaI and XmaI restriction sites in pcDNA3.1(+) ZKSCAN1 Scarless-Split eGFP, pcDNA3.1(+) Laccase2 Scarless-Split eGFP, pcDNA3.1(+) POLR2A Scarless-Split eGFP, and pcDNA3.1(+) IGF2BP1 Scarless-Split eGFP, respectively. Mouse mascRNA is indicated.

GGGCCCGTTTAAACCCGCTGATCAGCCTCGAGACGCTGGTGGCTGGCACTCCTGGTTTCCAGGACGGGGTTCAAGTCCCTGCGGTG
TCTCTTCTGAGGCGGAAAGAACCAGCTGGGGCTCTAGGGGGTATCCCCACGCGCCCTGTAGCGGCGCATTAAGCGCGGCGGGTGTG
GTGGTTACGCGCAGCGTGACCGCTACACTTGCCAGCGCCCTAGCGCCCGCTCCTTTCGCTTTCTTCCCTTCCTTTCTCGCCACGTT
CGCCGGCTTTCCCCGTCAAGCTCTAAATCGGGGGCTCCCTTTAGGGTTCCGATTTAGTGCTTTACGGCACCTCGACCCCAAAAAAC
TTGATTAGGGTGATGGTTCACGTAGTGGGCCATCGCCCTGATAGACGGTTTTTCGCCCTTTGACGTTGGAGTCCACGTTCTTTAAT
AGTGGACTCTTGTTCCAAACTGGAACAACACTCAACCCTATCTCGGTCTATTCTTTTGATTTATAAGGGATTTTGCCGATTTCGGC
CTATTGGTTAAAAAATGAGCTGATTTAACAAAAATTTAACGCGAATTAATTCTGTGGAATGTGTGTCAGTTAGGGTGTGGAAAGTC
CCCAGGCTCCCCAGCAGGCAGAAGTATGCAAAGCATGCATCTCAATTAGTCAGCAACCAGGTGTGGAAAGTCCCCAGGCTCCCCAG
CAGGCAGAAGTATGCAAAGCATGCATCTCAATTAGTCAGCAACCATAGTCCCGCCCCTAACTCCGCCCATCCCGCCCCTAACTCCG
CCCAGTTCCGCCCATTCTCCGCCCCATGGCTGACTAATTTTTTTTATTTATGCAGAGGCCGAGGCCGCCTCTGCCTCTGAGCTATT
CCAGAAGTAGTGAGGAGGCTTTTTTGGAGGCCTAGGCTTTTGCAAAAAGCT

To generate the following plasmids with the bGH poly(A) signal replaced with a mutated version of mascRNA (Mut 7):

**pcDNA3.1(+) ZKSCAN1 Scarless-Split eGFP (mascRNA Mut 7)**
**pcDNA3.1(+) Laccase2 Scarless-Split eGFP (mascRNA Mut 7)**
**pcDNA3.1(+) POLR2A Scarless-Split eGFP (mascRNA Mut 7)**

**pcDNA3.1(+) IGF2BP1 Scarless-Split eGFP (mascRNA Mut 7)**

the following sequence was cloned between the XbaI and XmaI restriction sites in pcDNA3.1(+) ZKSCAN1 Scarless-Split eGFP, pcDNA3.1(+) Laccase2 Scarless-Split eGFP, pcDNA3.1(+) POLR2A Scarless-Split eGFP, and pcDNA3.1(+) IGF2BP1 Scarless-Split eGFP, respectively. mascRNA Mut 7 is indicated.

GGGCCCGTTTAAACCCGCTGATCAGCCTCGAGACGCTGGTGGCTGGCACTCCTGGTTTCCAGGACGGGGTTCAAGTCCCTGCGGTA
TCTCTTCTGAGGCGGAAAGAACCAGCTGGGGCTCTAGGGGGTATCCCCACGCGCCCTGTAGCGGCGCATTAAGCGCGGCGGGTGTG
GTGGTTACGCGCAGCGTGACCGCTACACTTGCCAGCGCCCTAGCGCCCGCTCCTTTCGCTTTCTTCCCTTCCTTTCTCGCCACGTT
CGCCGGCTTTCCCCGTCAAGCTCTAAATCGGGGGCTCCCTTTAGGGTTCCGATTTAGTGCTTTACGGCACCTCGACCCCAAAAAAC
TTGATTAGGGTGATGGTTCACGTAGTGGGCCATCGCCCTGATAGACGGTTTTTCGCCCTTTGACGTTGGAGTCCACGTTCTTTAAT
AGTGGACTCTTGTTCCAAACTGGAACAACACTCAACCCTATCTCGGTCTATTCTTTTGATTTATAAGGGATTTTGCCGATTTCGGC
CTATTGGTTAAAAAATGAGCTGATTTAACAAAAATTTAACGCGAATTAATTCTGTGGAATGTGTGTCAGTTAGGGTGTGGAAAGTC
CCCAGGCTCCCCAGCAGGCAGAAGTATGCAAAGCATGCATCTCAATTAGTCAGCAACCAGGTGTGGAAAGTCCCCAGGCTCCCCAG
CAGGCAGAAGTATGCAAAGCATGCATCTCAATTAGTCAGCAACCATAGTCCCGCCCCTAACTCCGCCCATCCCGCCCCTAACTCCG
CCCAGTTCCGCCCATTCTCCGCCCCATGGCTGACTAATTTTTTTTATTTATGCAGAGGCCGAGGCCGCCTCTGCCTCTGAGCTATT
CCAGAAGTAGTGAGGAGGCTTTTTTGGAGGCCTAGGCTTTTGCAAAAAGCT

To generate the following plasmids with the bGH poly(A) signal replaced with a mutated version of mascRNA (Mut 10):

**pcDNA3.1(+) ZKSCAN1 Scarless-Split eGFP (mascRNA Mut 10)**
**pcDNA3.1(+) Laccase2 Scarless-Split eGFP (mascRNA Mut 10)**
**pcDNA3.1(+) POLR2A Scarless-Split eGFP (mascRNA Mut 10)**
**pcDNA3.1(+) IGF2BP1 Scarless-Split eGFP (mascRNA Mut 10)**

the following sequence was cloned between the XbaI and XmaI restriction sites in pcDNA3.1(+) ZKSCAN1 Scarless-Split eGFP, pcDNA3.1(+) Laccase2 Scarless-Split eGFP, pcDNA3.1(+) POLR2A Scarless-Split eGFP, and pcDNA3.1(+) IGF2BP1 Scarless-Split eGFP, respectively. mascRNA Mut 10 is indicated.

GGGCCCGTTTAAACCCGCTGATCAGCCTCGAGGCGCTGGTGGCTGGCACTCCTGGTTTCCAGGACGGGGTTCAAGTCCCTGCGGTA
CCGCTTCTGAGGCGGAAAGAACCAGCTGGGGCTCTAGGGGGTATCCCCACGCGCCCTGTAGCGGCGCATTAAGCGCGGCGGGTGTG
GTGGTTACGCGCAGCGTGACCGCTACACTTGCCAGCGCCCTAGCGCCCGCTCCTTTCGCTTTCTTCCCTTCCTTTCTCGCCACGTT
CGCCGGCTTTCCCCGTCAAGCTCTAAATCGGGGGCTCCCTTTAGGGTTCCGATTTAGTGCTTTACGGCACCTCGACCCCAAAAAAC
TTGATTAGGGTGATGGTTCACGTAGTGGGCCATCGCCCTGATAGACGGTTTTTCGCCCTTTGACGTTGGAGTCCACGTTCTTTAAT
AGTGGACTCTTGTTCCAAACTGGAACAACACTCAACCCTATCTCGGTCTATTCTTTTGATTTATAAGGGATTTTGCCGATTTCGGC
CTATTGGTTAAAAAATGAGCTGATTTAACAAAAATTTAACGCGAATTAATTCTGTGGAATGTGTGTCAGTTAGGGTGTGGAAAGTC
CCCAGGCTCCCCAGCAGGCAGAAGTATGCAAAGCATGCATCTCAATTAGTCAGCAACCAGGTGTGGAAAGTCCCCAGGCTCCCCAG
CAGGCAGAAGTATGCAAAGCATGCATCTCAATTAGTCAGCAACCATAGTCCCGCCCCTAACTCCGCCCATCCCGCCCCTAACTCCG
CCCAGTTCCGCCCATTCTCCGCCCCATGGCTGACTAATTTTTTTTATTTATGCAGAGGCCGAGGCCGCCTCTGCCTCTGAGCTATT
CCAGAAGTAGTGAGGAGGCTTTTTTGGAGGCCTAGGCTTTTGCAAAAAGCT

To generate the pCIRCUS plasmids, the following sequences were cloned between the AflII and NotI restriction sites in **pcDNA3.1(+) CircRNA Mini Vector** (Addgene #60648). dTomato ORF, mouse MALAT1 triple helix, mascRNA (WT or mutant), split eGFP ORF or guide RNA sequence are indicated. Flanking sequences that drive circRNA production are in lowercase.

#### pCIRCUS dTomato + WT mascRNA + ZKSCAN1 Scarless-Split eGFP

GGCCACCATGGTGAGCAAGGGCGAGGAGGTCATCAAAGAGTTCATGCGCTTCAAGGTGCGCATGGAGGGCTCCATGAACGGCCACG
AGTTCGAGATCGAGGGCGAGGGCGAGGGCCGCCCCTACGAGGGCACCCAGACCGCCAAGCTGAAGGTGACCAAGGGCGGCCCCCTG
CCCTTCGCCTGGGACATCCTGTCCCCCCAGTTCATGTACGGCTCCAAGGCGTACGTGAAGCACCCCGCCGACATCCCCGATTACAA
GAAGCTGTCCTTCCCCGAGGGCTTCAAGTGGGAGCGCGTGATGAACTTCGAGGACGGCGGTCTGGTGACCGTGACCCAGGACTCCT
CCCTGCAGGACGGCACGCTGATCTACAAGGTGAAGATGCGCGGCACCAACTTCCCCCCCGACGGCCCCGTAATGCAGAAGAAGACC
ATGGGCTGGGAGGCCTCCACCGAGCGCCTGTACCCCCGCGACGGCGTGCTGAAGGGCGAGATCCACCAGGCCCTGAAGCTGAAGGA
CGGCGGCCACTACCTGGTGGAGTTCAAGACCATCTACATGGCCAAGAAGCCCGTGCAACTGCCCGGCTACTACTACGTGGACACCA
AGCTGGACATCACCTCCCACAACGAGGACTACACCATCGTGGAACAGTACGAGCGCTCCGAGGGCCGCCACCACCTGTTCCTGTAC
GGCATGGACGAGCTGTACAAGGATTCGTCAGTAGGGTTGTAAAGGTTTTTCTTTTCCTGAGAAAACAACCTTTTGTTTTCTCAGGT
TTTGCTTTTTGGCCTTTCCCTAGCTTTAAAAAAAAAAAAAAGCAAAAGACGCTGGTGGCTGGCACTCCTGGTTTCCAGGACGGGGTT
CAAGTCCCTGCGGTGTCTTTGCTTACCGAGCTCGGATCCACTAGTCCAGTGTGGTGGAATTcagtgacagtggagattgtacagtt
ttttcctcgatttgtcaggatttttttttttgacggagtttaacttcttgtctcccaggtaggaagtgcagtggcgtaatctcgg
ctcactacaacctccacctcctgggttcaagcgtttctcctgcctcagctttccgagtagctgggattacaggcgcctgccaccat
gccctgctgacttttgtatttttagtagagacggggtttcaccatgttggccaggctggtcttgaactcctgaccgcaggcgattg
gcctgcctcggcctcccaaagtgctgagattacaggcgtgagccaccacccccggcctcaggagcgttctgatagtgcctcgatgt
gctgcctcctataaagtgttagcagcacagatcacttttgtaaaggtacgtactaatgactttttttttatacttcagGAGCGCA
CCATCTTCTTCAAGGACGACGGCAACTACAAGACCCGCGCCGAGGTGAAGTTCGAGGGCGACACCCTGGTGAACCGCATCGAGCTG
AAGGGCATCGACTTCAAGGAGGACGGCAACATCCTGGGGCACAAGCTGGAGTACAACCACAACAGCCACAACGTCTATATCATGGC
CGACAAGCAGAAGAACGGCATCAAGGTGAACTTCAAGATCCGCCACAACATCGAGGACGGCAGCGTGCAGCTCGCCGACCACTACC
AGCAGAACACCCCCATCGGCGACGGCCCCGTGCTGCTGCCCGACAACCACTACCTGAGCACCCAGTCCGCCCTGAGCAAAGACCCC
AACGAGAAGCGCGATCACATGGTCCTGCTGGAGTTCGTGACCGCCGCCGGGATCACTCTCGGCATGGACGAGCTGTACAAGTAAGG
ATCTGAATTCTGCAGTCGACGATCCGCCCCTCTCCCTCCCCCCCCCCTAACGTTACTGGCCGAAGCCGCTTGGAATAAGGCCGGTG
TGCGTTTGTCTATATGTTATTTTCCACCATATTGCCGTCTTTTGGCAATGTGAGGGCCCGGAAACCTGGCCCTGTCTTCTTGACGA
GCATTCCTAGGGGTCTTTCCCCTCTCGCCAAAGGAATGCAAGGTCTGTTGAATGTCGTGAAGGAAGCAGTTCCTCTGGAAGCTTCT
TGAAGACAAACAACGTCTGTAGCGACCCTTTGCAGGCAGCGGAACCCCCCACCTGGCGACAGGTGCCTCTGCGGCCAAAAGCCACG
TGTATAAGATACACCTGCAAAGGCGGCACAACCCCAGTGCCACGTTGTGAGTTGGATAGTTGTGGAAAGAGTCAAATGGCTCTCCT
CAAGCGTATTCAACAAGGGGCTGAAGGATGCCCAGAAGGTACCCCATTGTATGGGATCTGATCTGGGGCCTCGGTACACATGCTTT
ACATGTGTTTAGTCGAGGTTAAAAAAACGTCTAGGCCCCCCGAACCACGGGGACGTGGTTTTCCTTTGAAAAACACGATGATAATA
TGGCCACAACCATGGTGAGCAAGGGCGAGGAGCTGTTCACCGGGGTGGTGCCCATCCTGGTCGAGCTGGACGGCGACGTAAACGGC
CACAAGTTCAGCGTGTCCGGCGAGGGCGAGGGCGATGCCACCTACGGCAAGCTGACCCTGAAGTTCATCTGCACCACCGGCAAGCT
GCCCGTGCCCTGGCCCACCCTCGTGACCACCCTGACCTACGGCGTGCAGTGCTTCAGCCGCTACCCCGACCACATGAAGCAGCACG
ACTTCTTCAAGTCCGCCATGCCCGAAGGCTACGTCCAGgtaagaagcaaggtttcatttaggggaagggaaatgattcaggacgag
agtctttgtgctgctgagtgcctgtgatgaagaagcatgttagtcctgggcaacgtagcgagaccccatctctacaaaaaatagaa
aaattagccaggtatagtggcgcacacctgtgattccagctacgcaggaggctgaggtgggaggattgcttgagcccaggaggttg
aggctgcagtgagctggaatcatgccactactccaacctgggcaacacagcaaggaccctgtctcaaaagctacttacagaaaaga
attaggctcggcacggtagctcacacctgtaatcccagcactttgggaggctgaggcgggcagatcacttgaggtcaggagtttga
gaccagcctggccaacatggtgaaaccttgtctctactaaaaatatgaaaattagccaggcatggtggcacattcctgtaatccca
gctactcgggaggctgaggcaggagaatcacttgaacccaggaggtggaggttgcagtaagccgagatcgtaccactgtgctctag
ccttggtgacagagcgagactgtcttaaaaaaaaaaaaaaaaaagaattaattaaaaatttaaaaaaaaatgaaaaaaagctgcat
gcttgtttttttgtttttagttattctacattgttgtcattattaccaaatattggggaaaatacaacttacagaccaatctcagga
gttaaatgttactacgaaggcaaatgaactatgcgtaatgaacctggtaggcatta

#### pCIRCUS dTomato + WT mascRNA + Laccase2 Scarless-Split eGFP

GGCCACCATGGTGAGCAAGGGCGAGGAGGTCATCAAAGAGTTCATGCGCTTCAAGGTGCGCATGGAGGGCTCCATGAACGGCCACG
AGTTCGAGATCGAGGGCGAGGGCGAGGGCCGCCCCTACGAGGGCACCCAGACCGCCAAGCTGAAGGTGACCAAGGGCGGCCCCCTG
CCCTTCGCCTGGGACATCCTGTCCCCCCAGTTCATGTACGGCTCCAAGGCGTACGTGAAGCACCCCGCCGACATCCCCGATTACAA
GAAGCTGTCCTTCCCCGAGGGCTTCAAGTGGGAGCGCGTGATGAACTTCGAGGACGGCGGTCTGGTGACCGTGACCCAGGACTCCT
CCCTGCAGGACGGCACGCTGATCTACAAGGTGAAGATGCGCGGCACCAACTTCCCCCCCGACGGCCCCGTAATGCAGAAGAAGACC
ATGGGCTGGGAGGCCTCCACCGAGCGCCTGTACCCCCGCGACGGCGTGCTGAAGGGCGAGATCCACCAGGCCCTGAAGCTGAAGGA
CGGCGGCCACTACCTGGTGGAGTTCAAGACCATCTACATGGCCAAGAAGCCCGTGCAACTGCCCGGCTACTACTACGTGGACACCA
AGCTGGACATCACCTCCCACAACGAGGACTACACCATCGTGGAACAGTACGAGCGCTCCGAGGGCCGCCACCACCTGTTCCTGTAC
GGCATGGACGAGCTGTACAAGGATTCGTCAGTAGGGTTGTAAAGGTTTTTCTTTTCCTGAGAAAACAACCTTTTGTTTTCTCAGGT
TTTGCTTTTTGGCCTTTCCCTAGCTTTAAAAAAAAAAAAAAGCAAAAGACGCTGGTGGCTGGCACTCCTGGTTTCCAGGACGGGGTTC
AAGTCCCTGCGGTGTCTTTGCTTACCGAGCTCGGATCCACTAGTCCAGTGTGGTGGAATTCcattgagaaatgactgagttccggt
gctctcaagtcattgatctttgtcgacttttatttggtctctgtaataacgacttcaaaaacattaaattctgttgcgaagccagt
aagctacaaaaagaaaaaacaagagagaatgctatagtcgtatagtatagtttcccgactatctgatacccattacttatctaggg
ggaatgcgaacccaaaattttatcagttttctcggatatcgatagatattggggaataaatttaaataaataaattttgggcgggt
ttagggcgtggcaaaaagtttttttggcaaatcgctagaaatttacaagacttataaaattatgaaaaaatacaacaaaattttaaa
cacgtgggcgtgacagttttgggcggttttagggcgttagagtaggcgaggacagggttacatcgactaggctttgatcctgatca
agaatatatatactttataccgcttccttctacatgttacctattttcaacgaatctagtatacctttttactgtacgatttatg
ggtataataataagctaaatcgagactaagttttattgttatatatatttttttttattttatgcagGAGCGCACCATCTTCTTCAA
GGACGACGGCAACTACAAGACCCGCGCCGAGGTGAAGTTCGAGGGCGACACCCTGGTGAACCGCATCGAGCTGAAGGGCATCGACT
TCAAGGAGGACGGCAACATCCTGGGGCACAAGCTGGAGTACAACCACAACAGCCACAACGTCTATATCATGGCCGACAAGCAGAAG

AACGGCATCAAGGTGAACTTCAAGATCCGCCACAACATCGAGGACGGCAGCGTGCAGCTCGCCGACCACTACCAGCAGAACACCCC
CATCGGCGACGGCCCCGTGCTGCTGCCCGACAACCACTACCTGAGCACCCAGTCCGCCCTGAGCAAAGACCCCAACGAGAAGCGCG
ATCACATGGTCCTGCTGGAGTTCGTGACCGCCGCCGGGATCACTCTCGGCATGGACGAGCTGTACAAGTAAGGATCTGAATTCTGC
AGTCGACGATCCGCCCCTCTCCCTCCCCCCCCCCTAACGTTACTGGCCGAAGCCGCTTGGAATAAGGCCGGTGTGCGTTTGTCTATA
TGTTATTTTCCACCATATTGCCGTCTTTTGGCAATGTGAGGGCCCGGAAACCTGGCCCTGTCTTCTTGACGAGCATTCCTAGGGGT
CTTTCCCCTCTCGCCAAAGGAATGCAAGGTCTGTTGAATGTCGTGAAGGAAGCAGTTCCTCTGGAAGCTTCTTGAAGACAAACAAC
GTCTGTAGCGACCCTTTGCAGGCAGCGGAACCCCCCACCTGGCGACAGGTGCCTCTGCGGCCAAAAGCCACGTGTATAAGATACAC
CTGCAAAGGCGGCACAACCCCAGTGCCACGTTGTGAGTTGGATAGTTGTGGAAAGAGTCAAATGGCTCTCCTCAAGCGTATTCAAC
AAGGGGCTGAAGGATGCCCAGAAGGTACCCCATTGTATGGGATCTGATCTGGGGCCTCGGTACACATGCTTTACATGTGTTTAGTC
GAGGTTAAAAAAACGTCTAGGCCCCCCGAACCACGGGGACGTGGTTTTCCTTTGAAAAACACGATGATAATATGGCCACAACCATG
GTGAGCAAGGGCGAGGAGCTGTTCACCGGGGTGGTGCCCATCCTGGTCGAGCTGGACGGCGACGTAAACGGCCACAAGTTCAGCGT
GTCCGGCGAGGGCGAGGGCGATGCCACCTACGGCAAGCTGACCCTGAAGTTCATCTGCACCACCGGCAAGCTGCCCGTGCCCTGGC
CCACCCTCGTGACCACCCTGACCTACGGCGTGCAGTGCTTCAGCCGCTACCCCGACCACATGAAGCAGCACGACTTCTTCAAGTCC
GCCATGCCCGAAGGCTACGTCCAGgtaagtattcaaaattccaaaatttttactagaaatattcgatttttaataggcagtttc
tatactattgtatactattgtagattcgttgaaaagtatgtaacaggaagaataaagcatttccgaccatgtaaagtatatatatt
cttaataaggatcaatagccgagtcgatctcgccatgtccgtctgtcttattgttttattaccgccgagacatcaggaactataaa
agctagaaggatgagttttagcatacagattctagagacaaggacgcagagcaagtttgttgatccatgctgccacgctttaactt
tctcaaattgcccaaaactgccatgcccacattttgaactattttcgaaatttttcataattgtattactcgtgtaaatttcca
tcaatttgccaaaaaactttttgtcacgcgttaacgccctaaagccgccaatttggtcacgcccacactattgagcaattatcaaa
ttttttctcattttattccccaatatctatcgatatccccgattatgaaattattaaatttcgcgttcgcattcacactagctgag
taacgagtatctgatagttggggaaatcgacttatttttatatacaatgaaaatgaatttaatcatatgaatatcgattatagct
ttttatttaatatgaatatttatttgggcttaaggtgtaacctcctcgacataagactcacatggcgcaggcacattgaagacaaa
aatactcattgtcgggtctcgcaccctccagcagcacctaaaattatgtcttcaattattgccaacattggagacacaattagtct
gtggcacctcag

#### pCIRCUS dTomato + WT mascRNA + Tornado-Split eGFP

GGCCACCATGGTGAGCAAGGGCGAGGAGGTCATCAAAGAGTTCATGCGCTTCAAGGTGCGCATGGAGGGCTCCATGAACGGCCACG
AGTTCGAGATCGAGGGCGAGGGCGAGGGCCGCCCCTACGAGGGCACCCAGACCGCCAAGCTGAAGGTGACCAAGGGCGGCCCCCTG
CCCTTCGCCTGGGACATCCTGTCCCCCCAGTTCATGTACGGCTCCAAGGCGTACGTGAAGCACCCCGCCGACATCCCCGATTACAA
GAAGCTGTCCTTCCCCGAGGGCTTCAAGTGGGAGCGCGTGATGAACTTCGAGGACGGCGGTCTGGTGACCGTGACCCAGGACTCCT
CCCTGCAGGACGGCACGCTGATCTACAAGGTGAAGATGCGCGGCACCAACTTCCCCCCCGACGGCCCCGTAATGCAGAAGAAGACC
ATGGGCTGGGAGGCCTCCACCGAGCGCCTGTACCCCCGCGACGGCGTGCTGAAGGGCGAGATCCACCAGGCCCTGAAGCTGAAGGA
CGGCGGCCACTACCTGGTGGAGTTCAAGACCATCTACATGGCCAAGAAGCCCGTGCAACTGCCCGGCTACTACTACGTGGACACCA
AGCTGGACATCACCTCCCACAACGAGGACTACACCATCGTGGAACAGTACGAGCGCTCCGAGGGCCGCCACCACCTGTTCCTGTAC
GGCATGGACGAGCTGTACAAGGATTCGTCAGTAGGGTTGTAAAGGTTTTTCTTTTCCTGAGAAAACAACCTTTTGTTTTCTCAGGT
TTTGCTTTTTGGCCTTTCCCTAGCTTTAAAAAAAAAAAAAAGCAAAAGACGCTGGTGGCTGGCACTCCTGGTTTCCAGGACGGGGTT
CAAGTCCCTGCGGTGTCTTTGCTTACCGAGCTCGGATCCACTAGTCCAGTGTGGTGGAATTCggccgcactcgccggtcccaagcc
cggataaaatgggagggggcgggaaaccgcctaaccatgccgagtgcggccgcGAGCGCACCATCTTCTTCAAGGACGACGGCAAC
TACAAGACCCGCGCCGAGGTGAAGTTCGAGGGCGACACCCTGGTGAACCGCATCGAGCTGAAGGGCATCGACTTCAAGGAGGACGG
CAACATCCTGGGGCACAAGCTGGAGTACAACCACAACAGCCACAACGTCTATATCATGGCCGACAAGCAGAAGAACGGCATCAAGG
TGAACTTCAAGATCCGCCACAACATCGAGGACGGCAGCGTGCAGCTCGCCGACCACTACCAGCAGAACACCCCCATCGGCGACGGC
CCCGTGCTGCTGCCCGACAACCACTACCTGAGCACCCAGTCCGCCCTGAGCAAAGACCCCAACGAGAAGCGCGATCACATGGTCCT
GCTGGAGTTCGTGACCGCCGCCGGGATCACTCTCGGCATGGACGAGCTGTACAAGTAAGGATCTGAATTCTGCAGTCGACGATCCG
CCCCTCTCCCTCCCCCCCCCCTAACGTTACTGGCCGAAGCCGCTTGGAATAAGGCCGGTGTGCGTTTGTCTATATGTTATTTTCCA
CCATATTGCCGTCTTTTGGCAATGTGAGGGCCCGGAAACCTGGCCCTGTCTTCTTGACGAGCATTCCTAGGGGTCTTTCCCCTCTC
GCCAAAGGAATGCAAGGTCTGTTGAATGTCGTGAAGGAAGCAGTTCCTCTGGAAGCTTCTTGAAGACAAACAACGTCTGTAGCGAC
CCTTTGCAGGCAGCGGAACCCCCCACCTGGCGACAGGTGCCTCTGCGGCCAAAAGCCACGTGTATAAGATACACCTGCAAAGGCGG
CACAACCCCAGTGCCACGTTGTGAGTTGGATAGTTGTGGAAAGAGTCAAATGGCTCTCCTCAAGCGTATTCAACAAGGGGCTGAAG
GATGCCCAGAAGGTACCCCATTGTATGGGATCTGATCTGGGGCCTCGGTACACATGCTTTACATGTGTTTAGTCGAGGTTAAAAAA
ACGTCTAGGCCCCCCGAACCACGGGGACGTGGTTTTCCTTTGAAAAACACGATGATAATATGGCCACAACCATGGTGAGCAAGGGC
GAGGAGCTGTTCACCGGGGTGGTGCCCATCCTGGTCGAGCTGGACGGCGACGTAAACGGCCACAAGTTCAGCGTGTCCGGCGAGGG
CGAGGGCGATGCCACCTACGGCAAGCTGACCCTGAAGTTCATCTGCACCACCGGCAAGCTGCCCGTGCCCTGGCCCACCCTCGTGA
CCACCCTGACCTACGGCGTGCAGTGCTTCAGCCGCTACCCCGACCACATGAAGCAGCACGACTTCTTCAAGTCCGCCATGCCCGAA
GGCTACGTCCAGgtggccgcggtcggcgtggactgtagaacactgccaatgccggtcccaagcccggataaaagtggagggtacag
tccacgc

#### pCIRCUS dTomato + mascRNA Mut 7 + ZKSCAN1 Scarless-Split eGFP

GGCCACCATGGTGAGCAAGGGCGAGGAGGTCATCAAAGAGTTCATGCGCTTCAAGGTGCGCATGGAGGGCTCCATGAACGGCCACG
AGTTCGAGATCGAGGGCGAGGGCGAGGGCCGCCCCTACGAGGGCACCCAGACCGCCAAGCTGAAGGTGACCAAGGGCGGCCCCCTG
CCCTTCGCCTGGGACATCCTGTCCCCCCAGTTCATGTACGGCTCCAAGGCGTACGTGAAGCACCCCGCCGACATCCCCGATTACAA
GAAGCTGTCCTTCCCCGAGGGCTTCAAGTGGGAGCGCGTGATGAACTTCGAGGACGGCGGTCTGGTGACCGTGACCCAGGACTCCT
CCCTGCAGGACGGCACGCTGATCTACAAGGTGAAGATGCGCGGCACCAACTTCCCCCCCGACGGCCCCGTAATGCAGAAGAAGACC

ATGGGCTGGGAGGCCTCCACCGAGCGCCTGTACCCCCGCGACGGCGTGCTGAAGGGCGAGATCCACCAGGCCCTGAAGCTGAAGGA
CGGCGGCCACTACCTGGTGGAGTTCAAGACCATCTACATGGCCAAGAAGCCCGTGCAACTGCCCGGCTACTACTACGTGGACACCA
AGCTGGACATCACCTCCCACAACGAGGACTACACCATCGTGGAACAGTACGAGCGCTCCGAGGGCCGCCACCACCTGTTCCTGTAC
GGCATGGACGAGCTGTACAAGGATTCGTCAGTAGGGTTGTAAAGGTTTTTCTTTTCCTGAGAAAACAACCTTTTGTTTTCTCAGGT
TTTGCTTTTTGGCCTTTCCCTAGCTTTAAAAAAAAAAAAAAGCAAAAGACGCTGGTGGCTGGCACTCCTGGTTTCCAGGACGGGGTT
CAAGTCCCTGCGGTATCTTTGCTTACCGAGCTCGGATCCACTAGTCCAGTGTGGTGGAATTcagtgacagtggagattgtacagtt
ttttcctcgatttgtcaggatttttttttttgacggagtttaacttcttgtctcccaggtaggaagtgcagtggcgtaatctcgg
ctcactacaacctccacctcctgggttcaagcgtttctcctgcctcagctttccgagtagctgggattacaggcgcctgccaccat
gccctgctgacttttgtatttttagtagagacggggtttcaccatgttggccaggctggtcttgaactcctgaccgcaggcgattg
gcctgcctcggcctcccaaagtgctgagattacaggcgtgagccaccacccccggcctcaggagcgttctgatagtgcctcgatgt
gctgcctcctataaagtgttagcagcacagatcacttttgtaaaggtacgtactaatgactttttttttatacttcagGAGCGCA
CCATCTTCTTCAAGGACGACGGCAACTACAAGACCCGCGCCGAGGTGAAGTTCGAGGGCGACACCCTGGTGAACCGCATCGAGCTG
AAGGGCATCGACTTCAAGGAGGACGGCAACATCCTGGGGCACAAGCTGGAGTACAACCACAACAGCCACAACGTCTATATCATGGC
CGACAAGCAGAAGAACGGCATCAAGGTGAACTTCAAGATCCGCCACAACATCGAGGACGGCAGCGTGCAGCTCGCCGACCACTACC
AGCAGAACACCCCCATCGGCGACGGCCCCGTGCTGCTGCCCGACAACCACTACCTGAGCACCCAGTCCGCCCTGAGCAAAGACCCC
AACGAGAAGCGCGATCACATGGTCCTGCTGGAGTTCGTGACCGCCGCCGGGATCACTCTCGGCATGGACGAGCTGTACAAGTAAGG
ATCTGAATTCTGCAGTCGACGATCCGCCCCTCTCCCTCCCCCCCCCCTAACGTTACTGGCCGAAGCCGCTTGGAATAAGGCCGGTG
TGCGTTTGTCTATATGTTATTTTCCACCATATTGCCGTCTTTTGGCAATGTGAGGGCCCGGAAACCTGGCCCTGTCTTCTTGACGA
GCATTCCTAGGGGTCTTTCCCCTCTCGCCAAAGGAATGCAAGGTCTGTTGAATGTCGTGAAGGAAGCAGTTCCTCTGGAAGCTTCT
TGAAGACAAACAACGTCTGTAGCGACCCTTTGCAGGCAGCGGAACCCCCCACCTGGCGACAGGTGCCTCTGCGGCCAAAAGCCACG
TGTATAAGATACACCTGCAAAGGCGGCACAACCCCAGTGCCACGTTGTGAGTTGGATAGTTGTGGAAAGAGTCAAATGGCTCTCCT
CAAGCGTATTCAACAAGGGGCTGAAGGATGCCCAGAAGGTACCCCATTGTATGGGATCTGATCTGGGGCCTCGGTACACATGCTTT
ACATGTGTTTAGTCGAGGTTAAAAAAACGTCTAGGCCCCCCGAACCACGGGGACGTGGTTTTCCTTTGAAAAACACGATGATAATA
TGGCCACAACCATGGTGAGCAAGGGCGAGGAGCTGTTCACCGGGGTGGTGCCCATCCTGGTCGAGCTGGACGGCGACGTAAACGGC
CACAAGTTCAGCGTGTCCGGCGAGGGCGAGGGCGATGCCACCTACGGCAAGCTGACCCTGAAGTTCATCTGCACCACCGGCAAGCT
GCCCGTGCCCTGGCCCACCCTCGTGACCACCCTGACCTACGGCGTGCAGTGCTTCAGCCGCTACCCCGACCACATGAAGCAGCACG
ACTTCTTCAAGTCCGCCATGCCCGAAGGCTACGTCCAGgtaagaagcaaggtttcatttaggggaagggaaatgattcaggacgag
agtctttgtgctgctgagtgcctgtgatgaagaagcatgttagtcctgggcaacgtagcgagaccccatctctacaaaaaatagaa
aaattagccaggtatagtggcgcacacctgtgattccagctacgcaggaggctgaggtgggaggattgcttgagcccaggaggttg
aggctgcagtgagctggaatcatgccactactccaacctgggcaacacagcaaggaccctgtctcaaaagctacttacagaaaaga
attaggctcggcacggtagctcacacctgtaatcccagcactttgggaggctgaggcgggcagatcacttgaggtcaggagtttga
gaccagcctggccaacatggtgaaaccttgtctctactaaaaatatgaaaattagccaggcatggtggcacattcctgtaatccca
gctactcgggaggctgaggcaggagaatcacttgaacccaggaggtggaggttgcagtaagccgagatcgtaccactgtgctctag
ccttggtgacagagcgagactgtcttaaaaaaaaaaaaaaaaaaagaattaattaaaaatttaaaaaaaaatgaaaaaaagctgcat
gcttgtttttttgtttttagttattctacattgttgtcattattaccaaatattggggaaaatacaacttacagaccaatctcagga
gttaaatgttactacgaaggcaaatgaactatgcgtaatgaacctggtaggcatta

#### pCIRCUS dTomato + mascRNA Mut 7 + Laccase2 Scarless-Split eGFP

GGCCACCATGGTGAGCAAGGGCGAGGAGGTCATCAAAGAGTTCATGCGCTTCAAGGTGCGCATGGAGGGCTCCATGAACGGCCACG
AGTTCGAGATCGAGGGCGAGGGCGAGGGCCGCCCCTACGAGGGCACCCAGACCGCCAAGCTGAAGGTGACCAAGGGCGGCCCCCTG
CCCTTCGCCTGGGACATCCTGTCCCCCCAGTTCATGTACGGCTCCAAGGCGTACGTGAAGCACCCCGCCGACATCCCCGATTACAA
GAAGCTGTCCTTCCCCGAGGGCTTCAAGTGGGAGCGCGTGATGAACTTCGAGGACGGCGGTCTGGTGACCGTGACCCAGGACTCCT
CCCTGCAGGACGGCACGCTGATCTACAAGGTGAAGATGCGCGGCACCAACTTCCCCCCCGACGGCCCCGTAATGCAGAAGAAGACC
ATGGGCTGGGAGGCCTCCACCGAGCGCCTGTACCCCCGCGACGGCGTGCTGAAGGGCGAGATCCACCAGGCCCTGAAGCTGAAGGA
CGGCGGCCACTACCTGGTGGAGTTCAAGACCATCTACATGGCCAAGAAGCCCGTGCAACTGCCCGGCTACTACTACGTGGACACCA
AGCTGGACATCACCTCCCACAACGAGGACTACACCATCGTGGAACAGTACGAGCGCTCCGAGGGCCGCCACCACCTGTTCCTGTAC
GGCATGGACGAGCTGTACAAGGATTCGTCAGTAGGGTTGTAAAGGTTTTTCTTTTCCTGAGAAAACAACCTTTTGTTTTCTCAGGT
TTTGCTTTTTGGCCTTTCCCTAGCTTTAAAAAAAAAAAAAAGCAAAAGACGCTGGTGGCTGGCACTCCTGGTTTCCAGGACGGGGTTC
AAGTCCCTGCGGTATCTTTGCTTACCGAGCTCGGATCCACTAGTCCAGTGTGGTGGAATTCcattgagaaatgactgagttccggt
gctctcaagtcattgatctttgtcgacttttatttggtctctgtaataacgacttcaaaaacattaaattctgttgcgaagccagt
aagctacaaaaagaaaaaacaagagagaatgctatagtcgtatagtatagtttcccgactatctgatacccattacttatctaggg
ggaatgcgaacccaaaattttatcagttttctcggatatcgatagatattggggaataaatttaaataaataaattttgggcgggt
ttagggcgtggcaaaaagtttttttggcaaatcgctagaaatttacaagacttataaaattatgaaaaaatacaacaaaattttaaa
cacgtgggcgtgacagttttgggcggttttagggcgttagagtaggcgaggacagggttacatcgactaggctttgatcctgatca
agaatatatatactttataccgcttccttctacatgttacctatttttcaacgaatctagtatacctttttactgtacgatttatg
ggtataataataagctaaatcgagactaagttttattgttatatatattttttttattttatgcagGAGCGCACCATCTTCTTCAA
GGACGACGGCAACTACAAGACCCGCGCCGAGGTGAAGTTCGAGGGCGACACCCTGGTGAACCGCATCGAGCTGAAGGGCATCGACT
TCAAGGAGGACGGCAACATCCTGGGGCACAAGCTGGAGTACAACCACAACAGCCACAACGTCTATATCATGGCCGACAAGCAGAAG
AACGGCATCAAGGTGAACTTCAAGATCCGCCACAACATCGAGGACGGCAGCGTGCAGCTCGCCGACCACTACCAGCAGAACACCCC
CATCGGCGACGGCCCCGTGCTGCTGCCCGACAACCACTACCTGAGCACCCAGTCCGCCCTGAGCAAAGACCCCAACGAGAAGCGCG
ATCACATGGTCCTGCTGGAGTTCGTGACCGCCGCCGGGATCACTCTCGGCATGGACGAGCTGTACAAGTAAGGATCTGAATTCTGC
AGTCGACGATCCGCCCCTCTCCCTCCCCCCCCCCTAACGTTACTGGCCGAAGCCGCTTGGAATAAGGCCGGTGTGCGTTTGTCTATA

TGTTATTTTCCACCATATTGCCGTCTTTTGGCAATGTGAGGGCCCGGAAACCTGGCCCTGTCTTCTTGACGAGCATTCCTAGGGGT
CTTTCCCCTCTCGCCAAAGGAATGCAAGGTCTGTTGAATGTCGTGAAGGAAGCAGTTCCTCTGGAAGCTTCTTGAAGACAAACAAC
GTCTGTAGCGACCCTTTGCAGGCAGCGGAACCCCCCACCTGGCGACAGGTGCCTCTGCGGCCAAAAGCCACGTGTATAAGATACAC
CTGCAAAGGCGGCACAACCCCAGTGCCACGTTGTGAGTTGGATAGTTGTGGAAAGAGTCAAATGGCTCTCCTCAAGCGTATTCAAC
AAGGGGCTGAAGGATGCCCAGAAGGTACCCCATTGTATGGGATCTGATCTGGGGCCTCGGTACACATGCTTTACATGTGTTTAGTC
GAGGTTAAAAAAACGTCTAGGCCCCCCGAACCACGGGGACGTGGTTTTCCTTTGAAAAACACGATGATAATATGGCCACAACCATG
GTGAGCAAGGGCGAGGAGCTGTTCACCGGGGTGGTGCCCATCCTGGTCGAGCTGGACGGCGACGTAAACGGCCACAAGTTCAGCGT
GTCCGGCGAGGGCGAGGGCGATGCCACCTACGGCAAGCTGACCCTGAAGTTCATCTGCACCACCGGCAAGCTGCCCGTGCCCTGGC
CCACCCTCGTGACCACCCTGACCTACGGCGTGCAGTGCTTCAGCCGCTACCCCGACCACATGAAGCAGCACGACTTCTTCAAGTCC
GCCATGCCCGAAGGCTACGTCCAGgtaagtattcaaaattccaaaatttttactagaaatattcgatttttaataggcagtttc
tatactattgtatactattgtagattcgttgaaaagtatgtaacaggaagaataaagcatttccgaccatgtaaagtatatatatt
cttaataaggatcaatagccgagtcgatctcgccatgtccgtctgtcttattgttttattaccgccgagacatcaggaactataaa
agctagaaggatgagttttagcatacagattctagagacaaggacgcagagcaagtttgttgatccatgctgccacgctttaactt
tctcaaattgcccaaaactgccatgcccacatttttgaactattttcgaaatttttcataattgtattactcgtgtaaatttcca
tcaatttgccaaaaaactttttgtcacgcgttaacgccctaaagccgccaatttggtcacgcccacactattgagcaattatcaaa
ttttttctcattttattccccaatatctatcgatatccccgattatgaaattattaaatttcgcgttcgcattcacactagctgag
taacgagtatctgatagttggggaaatcgacttatttttatatacaatgaaaatgaatttaatcatatgaatatcgattatagct
ttttatttaatatgaatatttatttgggcttaaggtgtaacctcctcgacataagactcacatggcgcaggcacattgaagacaaa
aatactcattgtcgggtctcgcaccctccagcagcacctaaaattatgtcttcaattattgccaacattggagacacaattagtct
gtggcacctcag

#### pCIRCUS dTomato + mascRNA Mut 7 + Tornado-Split eGFP

GGCCACCATGGTGAGCAAGGGCGAGGAGGTCATCAAAGAGTTCATGCGCTTCAAGGTGCGCATGGAGGGCTCCATGAACGGCCACG
AGTTCGAGATCGAGGGCGAGGGCGAGGGCCGCCCCTACGAGGGCACCCAGACCGCCAAGCTGAAGGTGACCAAGGGCGGCCCCCTG
CCCTTCGCCTGGGACATCCTGTCCCCCCAGTTCATGTACGGCTCCAAGGCGTACGTGAAGCACCCCGCCGACATCCCCGATTACAA
GAAGCTGTCCTTCCCCGAGGGCTTCAAGTGGGAGCGCGTGATGAACTTCGAGGACGGCGGTCTGGTGACCGTGACCCAGGACTCCT
CCCTGCAGGACGGCACGCTGATCTACAAGGTGAAGATGCGCGGCACCAACTTCCCCCCCGACGGCCCCGTAATGCAGAAGAAGACC
ATGGGCTGGGAGGCCTCCACCGAGCGCCTGTACCCCCGCGACGGCGTGCTGAAGGGCGAGATCCACCAGGCCCTGAAGCTGAAGGA
CGGCGGCCACTACCTGGTGGAGTTCAAGACCATCTACATGGCCAAGAAGCCCGTGCAACTGCCCGGCTACTACTACGTGGACACCA
AGCTGGACATCACCTCCCACAACGAGGACTACACCATCGTGGAACAGTACGAGCGCTCCGAGGGCCGCCACCACCTGTTCCTGTAC
GGCATGGACGAGCTGTACAAGGATTCGTCAGTAGGGTTGTAAAGGTTTTTCTTTTCCTGAGAAAACAACCTTTTGTTTTCTCAGGT
TTTGCTTTTTGGCCTTTCCCTAGCTTTAAAAAAAAAAAAAAGCAAAAGACGCTGGTGGCTGGCACTCCTGGTTTCCAGGACGGGGTT
CAAGTCCCTGCGGTATCTTTGCTTACCGAGCTCGGATCCACTAGTCCAGTGTGGTGGAATTCggccgcactcgccggtcccaagcc
cggataaaatgggagggggcgggaaaccgcctaaccatgccgagtgcggccgcGAGCGCACCATCTTCTTCAAGGACGACGGCAAC
TACAAGACCCGCGCCGAGGTGAAGTTCGAGGGCGACACCCTGGTGAACCGCATCGAGCTGAAGGGCATCGACTTCAAGGAGGACGG
CAACATCCTGGGGCACAAGCTGGAGTACAACCACAACAGCCACAACGTCTATATCATGGCCGACAAGCAGAAGAACGGCATCAAGG
TGAACTTCAAGATCCGCCACAACATCGAGGACGGCAGCGTGCAGCTCGCCGACCACTACCAGCAGAACACCCCCATCGGCGACGGC
CCCGTGCTGCTGCCCGACAACCACTACCTGAGCACCCAGTCCGCCCTGAGCAAAGACCCCAACGAGAAGCGCGATCACATGGTCCT
GCTGGAGTTCGTGACCGCCGCCGGGATCACTCTCGGCATGGACGAGCTGTACAAGTAAGGATCTGAATTCTGCAGTCGACGATCCG
CCCCTCTCCCTCCCCCCCCCCTAACGTTACTGGCCGAAGCCGCTTGGAATAAGGCCGGTGTGCGTTTGTCTATATGTTATTTTCCA
CCATATTGCCGTCTTTTGGCAATGTGAGGGCCCGGAAACCTGGCCCTGTCTTCTTGACGAGCATTCCTAGGGGTCTTTCCCCTCTC
GCCAAAGGAATGCAAGGTCTGTTGAATGTCGTGAAGGAAGCAGTTCCTCTGGAAGCTTCTTGAAGACAAACAACGTCTGTAGCGAC
CCTTTGCAGGCAGCGGAACCCCCCACCTGGCGACAGGTGCCTCTGCGGCCAAAAGCCACGTGTATAAGATACACCTGCAAAGGCGG
CACAACCCCAGTGCCACGTTGTGAGTTGGATAGTTGTGGAAAGAGTCAAATGGCTCTCCTCAAGCGTATTCAACAAGGGGCTGAAG
GATGCCCAGAAGGTACCCCATTGTATGGGATCTGATCTGGGGCCTCGGTACACATGCTTTACATGTGTTTAGTCGAGGTTAAAAAA
ACGTCTAGGCCCCCCGAACCACGGGGACGTGGTTTTCCTTTGAAAAACACGATGATAATATGGCCACAACCATGGTGAGCAAGGGC
GAGGAGCTGTTCACCGGGGTGGTGCCCATCCTGGTCGAGCTGGACGGCGACGTAAACGGCCACAAGTTCAGCGTGTCCGGCGAGGG
CGAGGGCGATGCCACCTACGGCAAGCTGACCCTGAAGTTCATCTGCACCACCGGCAAGCTGCCCGTGCCCTGGCCCACCCTCGTGA
CCACCCTGACCTACGGCGTGCAGTGCTTCAGCCGCTACCCCGACCACATGAAGCAGCACGACTTCTTCAAGTCCGCCATGCCCGAA
GGCTACGTCCAGgtggccgcggtcggcgtggactgtagaacactgccaatgccggtcccaagcccggataaaagtggagggtacag
tccacgc

#### pCIRCUS dTomato + mascRNA Mut 7 + Tornado-RAB7A Guide

GGCCACCATGGTGAGCAAGGGCGAGGAGGTCATCAAAGAGTTCATGCGCTTCAAGGTGCGCATGGAGGGCTCCATGAACGGCCACG
AGTTCGAGATCGAGGGCGAGGGCGAGGGCCGCCCCTACGAGGGCACCCAGACCGCCAAGCTGAAGGTGACCAAGGGCGGCCCCCTG
CCCTTCGCCTGGGACATCCTGTCCCCCCAGTTCATGTACGGCTCCAAGGCGTACGTGAAGCACCCCGCCGACATCCCCGATTACAA
GAAGCTGTCCTTCCCCGAGGGCTTCAAGTGGGAGCGCGTGATGAACTTCGAGGACGGCGGTCTGGTGACCGTGACCCAGGACTCCT
CCCTGCAGGACGGCACGCTGATCTACAAGGTGAAGATGCGCGGCACCAACTTCCCCCCCGACGGCCCCGTAATGCAGAAGAAGACC
ATGGGCTGGGAGGCCTCCACCGAGCGCCTGTACCCCCGCGACGGCGTGCTGAAGGGCGAGATCCACCAGGCCCTGAAGCTGAAGGA
CGGCGGCCACTACCTGGTGGAGTTCAAGACCATCTACATGGCCAAGAAGCCCGTGCAACTGCCCGGCTACTACTACGTGGACACCA
AGCTGGACATCACCTCCCACAACGAGGACTACACCATCGTGGAACAGTACGAGCGCTCCGAGGGCCGCCACCACCTGTTCCTGTAC
GGCATGGACGAGCTGTACAAGGATTCGTCAGTAGGGTTGTAAAGGTTTTTCTTTTCCTGAGAAAACAACCTTTTGTTTTCTCAGGT

TTTGCTTTTTGGCCTTTCCCTAGCTTTAAAAAAAAAAAAAAGCAAAAGACGCTGGTGGCTGGCACTCCTGGTTTCCAGGACGGGGTT
CAAGTCCCTGCGGTATCTTTGCTTACCGAGCTCGGATCCACTAGTCCAGTGTGGTGGAATTCggccgcactcgccggtcccaagcc
cggataaaatgggagggggcgggaaaccgcctaaccatgccgagtgcggccgcCATGTTGTTCTCGTCTCCTCGACACCCTGCCGC
CAGCTGGATTTCCAAAATTAAACACATAATCCAAGAAAAATTGCATATATTAACATGTACAAACCCTGGAGAGATGAAAAGCTAAA
AAAAGGCGTACATAATTCTTAAAAAGGTGTCGAGAAGAGGAGAACAATATCTTTgtggccgcggtcggcgtggactgtagaacact
gccaatgccggtcccaagcccggataaaagtggagggtacagtccacgc

#### pCIRCUS dTomato + mascRNA Mut 7 + Tornado-Control Guide

GGCCACCATGGTGAGCAAGGGCGAGGAGGTCATCAAAGAGTTCATGCGCTTCAAGGTGCGCATGGAGGGCTCCATGAACGGCCACG
AGTTCGAGATCGAGGGCGAGGGCGAGGGCCGCCCCTACGAGGGCACCCAGACCGCCAAGCTGAAGGTGACCAAGGGCGGCCCCCTG
CCCTTCGCCTGGGACATCCTGTCCCCCCAGTTCATGTACGGCTCCAAGGCGTACGTGAAGCACCCCGCCGACATCCCCGATTACAA
GAAGCTGTCCTTCCCCGAGGGCTTCAAGTGGGAGCGCGTGATGAACTTCGAGGACGGCGGTCTGGTGACCGTGACCCAGGACTCCT
CCCTGCAGGACGGCACGCTGATCTACAAGGTGAAGATGCGCGGCACCAACTTCCCCCCCGACGGCCCCGTAATGCAGAAGAAGACC
ATGGGCTGGGAGGCCTCCACCGAGCGCCTGTACCCCCGCGACGGCGTGCTGAAGGGCGAGATCCACCAGGCCCTGAAGCTGAAGGA
CGGCGGCCACTACCTGGTGGAGTTCAAGACCATCTACATGGCCAAGAAGCCCGTGCAACTGCCCGGCTACTACTACGTGGACACCA
AGCTGGACATCACCTCCCACAACGAGGACTACACCATCGTGGAACAGTACGAGCGCTCCGAGGGCCGCCACCACCTGTTCCTGTAC
GGCATGGACGAGCTGTACAAGGATTCGTCAGTAGGGTTGTAAAGGTTTTTCTTTTCCTGAGAAAACAACCTTTTGTTTTCTCAGGT
TTTGCTTTTTGGCCTTTCCCTAGCTTTAAAAAAAAAAAAAAGCAAAAGACGCTGGTGGCTGGCACTCCTGGTTTCCAGGACGGGGTT
CAAGTCCCTGCGGTATCTTTGCTTACCGAGCTCGGATCCACTAGTCCAGTGTGGTGGAATTCggccgcactcgccggtcccaagcc
cggataaaatgggagggggcgggaaaccgcctaaccatgccgagtgcggccgcACGGGCAGGTGTTGTGTACGTCACCTGCCTCGg
tggccgcggtcggcgtggactgtagaacactgccaatgccggtcccaagcccggataaaagtggagggtacagtccacgc

**Summary of plasmid usage**

| Plasmid Name | Figure Panels | Supplementary Figure Panels |
|---|---|---|
| pcDNA3.1(+) CircRNA Mini Vector | | |
| pcDNA3.1(+) circRNA Mini Scarless MCS | | |
| pcDNA3.1(+) ZKSCAN1 Scarless MCS | | |
| pcDNA3.1(+) Laccase2 Scarless MCS | | |
| pcDNA3.1(+) POLR2A Scarless MCS | | |
| pcDNA3.1(+) ciRS-7 Scarless MCS | | |
| pcDNA3.1(+) IGF2BP1 Scarless MCS | | |
| pcDNA3.1(+) Tornado PaqCI MCS | | |
| pcDNA3.1(+) circRNA Mini-Split eGFP | 2B, 2C, 2D, 2E, 2F, 2G, 4A, 4B, 4C, 4D, 4E, 4F, 4G, 4H, 4I | S2B, S3, S4B, S4C, S7, S11A, S11B, S11C |
| pcDNA3.1(+) circRNA Mini Scarless-Split eGFP | 2B, 2C, 2D, 2E, 2F, 2G, 4A, 4B, 4C, 4D, 4E, 4F, 4G, 4H, 4I | S2C, S3, S4B, S4C, S7, S11A, S11B, S11C |
| pcDNA3.1(+) ZKSCAN1 Scarless-Split eGFP | 2B, 2C, 2D, 2E, 2F, 2G, 4A, 4B, 4C, 4D, 4E, 4F, 4G, 4H, 4I, 5B, 5C | S2C, S3, S4B, S4C, S5B, S5C, S6A, S6B, S7, S11A, S11B, S11C, S13, S14D |
| pcDNA3.1(+) Laccase2 Scarless-Split eGFP | 2B, 2C, 2D, 2E, 2F, 2G, 4A, 4B, 4C, 4D, 4E, 4F, 4G, 4H, 4I, 5B, 5C | S2C, S3, S4B, S4C, S5B, S5C, S6A, S6B, S7, S11A, S11B, S11C, S13, S14D |
| pcDNA3.1(+) POLR2A Scarless-Split eGFP | 2B, 2C, 2D, 2E, 2F, 2G, 4A, 4B, 4C, 4D, 4E, 4F, 4G, 4H, 4I, 5B, 5C | S2C, S3, S4B, S4C, S5B, S5C, S7, S11A, S11B, S11C, S13 |
| pcDNA3.1(+) ciRS-7 Scarless-Split eGFP | 2B, 2C, 2D, 2E, 2F, 2G, 4A, 4B, 4C, 4D, 4E, 4F, 4G, 4H, 4I | S2C, S3, S4B, S4C, S6A, S6B, S7, S11A, S11B, S11C |
| pcDNA3.1(+) IGF2BP1 Scarless-Split eGFP | 2B, 2C, 2D, 2E, 2F, 2G, 4A, 4B, 4C, 4D, 4E, 4F, 4G, 4H, 4I, 5B, 5C | S2C, S3, S4B, S4C, S5B, S5C, S7, S11A, S11B, S11C, S12B, S12C, S13 |
| pcDNA3.1(+) Tornado PaqCI-Split eGFP | 2B, 2C, 2D, 2E, 2F, 2G, 4A, 4B, 4C, 4D, 4E, 4F, 4G, 4H, 4I | S1B, S3, S4B, S4C, S5B, S7, S8B, S8C, S8D, S11A, S11B, S11C, S14D |
| pcDNA3.1(+) circRNA Mini-Split eGFP Delta Repeat | | S4B, S4C |
| pcDNA3.1(+) circRNA Mini Scarless-Split eGFP Delta Repeat | | S4B, S4C |
| pcDNA3.1(+) ZKSCAN1 Scarless-Split eGFP Delta Repeat | | S4B, S4C |
| pcDNA3.1(+) Laccase2 Scarless-Split eGFP Delta Repeat | | S4B, S4C |
| pcDNA3.1(+) POLR2A Scarless-Split eGFP Delta Repeat | | S4B, S4C |
| pcDNA3.1(+) ciRS-7 Scarless-Split eGFP Delta Repeat | | S4B, S4C |
| pcDNA3.1(+) IGF2BP1 Scarless-Split eGFP Delta Repeat | | S4B, S4C |
| pcDNA3.1(+) ciRS-7 Scarless-Split eGFP (Laccase2 ss) | | S6B |
| pcDNA3.1(+) ciRS-7 Scarless-Split eGFP (ZKSCAN1 ss) | | S6B |
| pcDNA3.1(+) circRNA Mini-ciRS-7 | 3B, 3C | S9A, S9C |
| pcDNA3.1(+) circRNA Mini Scarless-ciRS-7 | 3B, 3C | S9A, S9C |
| pcDNA3.1(+) ZKSCAN1 Scarless-ciRS-7 | 3B, 3C | S9A, S9C |
| pcDNA3.1(+) Laccase2 Scarless-ciRS-7 | 3B, 3C | S9A, S9C |
| pcDNA3.1(+) POLR2A Scarless-ciRS-7 | 3B, 3C | S9A, S9C |
| pcDNA3.1(+) ciRS-7 Scarless-ciRS-7 | 3B, 3C | S9A, S9C |
| pcDNA3.1(+) Tornado PaqCI-ciRS-7 | 3B, 3C | S9A, S9C |
| pcDNA3.1(+) circRNA Mini-ZKSCAN1 Exons 2-3 | 3E, 3F | S9A, S9E |
| pcDNA3.1(+) circRNA Mini Scarless-ZKSCAN1 Exons 2-3 | 3E, 3F | S9A, S9E |
| pcDNA3.1(+) ZKSCAN1 Scarless-ZKSCAN1 Exons 2-3 | 3E, 3F | S9A, S9E |
| pcDNA3.1(+) Laccase2 Scarless-ZKSCAN1 Exons 2-3 | 3E, 3F | S9A, S9E |
| pcDNA3.1(+) POLR2A Scarless-ZKSCAN1 Exons 2-3 | 3E, 3F | S9A, S9E |
| pcDNA3.1(+) ciRS-7 Scarless-ZKSCAN1 Exons 2-3 | 3E, 3F | S9A, S9E |
| pcDNA3.1(+) Tornado PaqCI-ZKSCAN1 Exons 2-3 | 3E, 3F | S9A, S9E |
| pcDNA3.1(+) circRNA Mini-NFASC Exons 26-27 | | S10B |
| pcDNA3.1(+) circRNA Mini Scarless-NFASC Exons 26-27 | | S10B |
| pcDNA3.1(+) ZKSCAN1 Scarless-NFASC Exons 26-27 | | S10B |
| pcDNA3.1(+) Laccase2 Scarless-NFASC Exons 26-27 | | S10B |
| pcDNA3.1(+) POLR2A Scarless-NFASC Exons 26-27 | | S10B |
| pcDNA3.1(+) ciRS-7 Scarless-NFASC Exons 26-27 | | S10B |
| pcDNA3.1(+) Tornado PaqCI-NFASC Exons 26-27 | | S10B |
| pcDNA3.1(+) IGF2BP1 Scarless-Split eGFP (Upstream miR-92a) | | S12B, S12C |
| pcDNA3.1(+) IGF2BP1 Scarless-Split eGFP (Upstream miR-92a Mut) | | S12B, S12C |
| pcDNA3.1(+) IGF2BP1 Scarless-Split eGFP (Downstream miR-92a) | | S12B, S12C |
| pcDNA3.1(+) IGF2BP1 Scarless-Split eGFP (Downstream miR-92a Mut) | | S12B, S12C |
| pcDNA3.1(+) IGF2BP1 Scarless-Split eGFP (Up & Downstream miR-92a) | | S12B, S12C |
| pcDNA3.1(+) IGF2BP1 Scarless-Split eGFP (Up & Downstream miR-92a Mut) | | S12B, S12C |
| pcDNA3.1(+) ZKSCAN1 Scarless-Split eGFP (mascRNA WT) | 5B, 5C | S13 |
| pcDNA3.1(+) ZKSCAN1 Scarless-Split eGFP (mascRNA Mut 7) | 5B, 5C | S13 |
| pcDNA3.1(+) ZKSCAN1 Scarless-Split eGFP (mascRNA Mut 10) | 5B, 5C | S13 |
| pcDNA3.1(+) Laccase2 Scarless-Split eGFP (mascRNA WT) | 5B, 5C | S13 |
| pcDNA3.1(+) Laccase2 Scarless-Split eGFP (mascRNA Mut 7) | 5B, 5C | S13 |
| pcDNA3.1(+) Laccase2 Scarless-Split eGFP (mascRNA Mut 10) | 5B, 5C | S13 |
| pcDNA3.1(+) POLR2A Scarless-Split eGFP (mascRNA WT) | 5B, 5C | S13 |
| pcDNA3.1(+) POLR2A Scarless-Split eGFP (mascRNA Mut 7) | 5B, 5C | S13 |
| pcDNA3.1(+) POLR2A Scarless-Split eGFP (mascRNA Mut 10) | 5B, 5C | S13 |
| pcDNA3.1(+) IGF2BP1 Scarless-Split eGFP (mascRNA WT) | 5B, 5C | S13 |
| pcDNA3.1(+) IGF2BP1 Scarless-Split eGFP (mascRNA Mut 7) | 5B, 5C | S13 |
| pcDNA3.1(+) IGF2BP1 Scarless-Split eGFP (mascRNA Mut 10) | 5B, 5C | S13 |
| pCIRCUS dTomato + WT mascRNA + ZKSCAN1 Scarless-Split eGFP | | S14B, S14C, S14D |
| pCIRCUS dTomato + WT mascRNA + Laccase2 Scarless-Split eGFP | | S14B, S14C, S14D |
| pCIRCUS dTomato + WT mascRNA + Tornado-Split eGFP | | S14B, S14C, S14D |
| pCIRCUS dTomato + mascRNA Mut 7 + ZKSCAN1 Scarless-Split eGFP | 6B, 6C | |
| pCIRCUS dTomato + mascRNA Mut 7 + Laccase2 Scarless-Split eGFP | 6B, 6C | |
| pCIRCUS dTomato + mascRNA Mut 7 + Tornado-Split eGFP | 6B, 6C | |
| pCIRCUS dTomato + mascRNA Mut 7 + Tornado-RAB7A Guide | 7C, 7D, 7E | S15B |
| pCIRCUS dTomato + mascRNA Mut 7 + Tornado-Control Guide | 7D, 7E | S15B |
